# VTA dopamine neuron activity produces spatially organized stimulus and action value representations through conditioned reinforcement

**DOI:** 10.1101/2025.11.04.685995

**Authors:** Alejandro Pan-Vazquez, Christopher A. Zimmerman, Brenna McMannon, Julie M. J. Fabre, Miranta Louka, Tingying Jia, Yotam Sagiv, Steven J. West, Mayo Faulkner, International Brain Laboratory, Peter Dayan, Ilana B. Witten

**Affiliations:** Princeton Neuroscience Institute, Princeton University, Princeton, NJ, USA; Department of Neurobiology, University of Utah, Salt Lake City, UT, USA; Sainsbury Wellcome Centre, University College London, London, UK; Institute of Neurology, University College London, London, UK; Max Planck Institute for Biological Cybernetics, Tübingen, Germany; University of Tübingen,Tübingen, Germany; Howard Hughes Medical Institute, Princeton University, Princeton, NJ, USA

## Abstract

What is the neural architecture by which dopamine (DA) determines choice? Reinforcement learning (RL) has suggested an algorithmic chain: prediction errors based on reward train predicted values for stimuli and actions, and thereby determine choice. Elements of this chain have been identified, but have yet to be convincingly assembled. Here we used optogenetic stimulation of putative prediction error neurons at the top of this chain – i.e., DA neurons in the ventral tegmental area (VTA) – and recorded the activity of thousands of neurons in their target structure – the striatum – in mice performing a task that dissociates VTA DA signalling, stimulus value, action value, and choice. Conventional RL models captured some features of the data, including DA-driven choice reinforcement and value correlates, but failed on two key observations: i) action and state value correlates were relatively weak in the VTA’s main target, the nucleus accumbens (NAc), and ii) VTA DA could not reinforce actions in the absence of a reward-predictive cue (CS+). Instead, we find that VTA DA produces stimulus value representations in the NAc, which in turn generates action value representations downstream (e.g. the CP). We formalize these observations in a Conditioned Reinforcer model, where VTA DA specifically acts on stimuli rather than actions, and these stimulus value representations are used to drive choice reinforcement.

## INTRODUCTION

How do animals make choices that allow them to obtain rewards? DA neurons are thought to contribute critically to this process by encoding an error signal for predicting rewards^1–4^. This error signal is hypothesized to instruct representations in downstream target neurons of the rewards predicted by different stimuli or actions (i.e., values), which animals can then use to make the most appropriate or valuable choice^5–7^.

There is extensive support for aspects of this model. Behaviorally, optogenetic stimulation of VTA DA neurons following an action can reinforce it^8–13^, which implies that VTA DA neurons can make actions valuable. Concomitantly, inhibiting these neurons can rob natural rewards of their reinforcing power^14–16^. More mechanistically, DA can generate synaptic plasticity at target sites in the striatum^17–21^ and elsewhere^22–24^ across vertebrates^25^ and invertebrates^26^, which is consistent with DA potentially acting as a teaching signal that is used to sculpt downstream representations of action value.

However, the core idea that DA release produces value representations in downstream neurons remains untested. This is because most previous work involving artificial activation of DA neurons has employed behavioral rather than neural readouts of value^9,27–34^. Conversely, previous work identifying value representations during behavior has not employed the direct activation of DA neurons^33,35–53^. In fact, despite the important characterizations of the activity patterns of DA neurons^1,4,15,34,54–79^, how such activity in turn alters downstream neural dynamics to mediate adaptive behavior remains an open question.

Thus, here we ask: is VTA DA neuron activity sufficient to produce downstream value correlates? And, if so, are they localized to the target sites of the VTA DA neurons, as might be expected if DA serves as a teaching signal for producing such representations?

## RESULTS

### Mice adjust their behavior using trial-and-error learning to obtain VTA DA stimulation

To address these questions, we developed a head-fixed probabilistic reversal learning task in which, rather than working for a palatable reward^13,15,80,81^, mice work to receive optogenetic stimulation of VTA DA neurons (Fig. 1a,b and Extended Data Fig. 1a). After an auditory “go cue”, the mice rotated a wheel positioned under their forepaws left or right. This choice resulted in the presentation of one of two auditory cues (CS+ or CS–). The CS+ signaled the opportunity to lick a spout to receive a 1 s burst of VTA DA stimulation. The CS– signalled the absence of stimulation. For a block of trials, turning the wheel in one direction led to the CS+ with a probability of 70%, whereas the other direction led to the CS+ with only a 10% probability. After a variable number of trials, and without any indication to the mouse, the outcome contingencies reversed so that the other choice led to the CS+ with higher probability.

**Figure 1.**
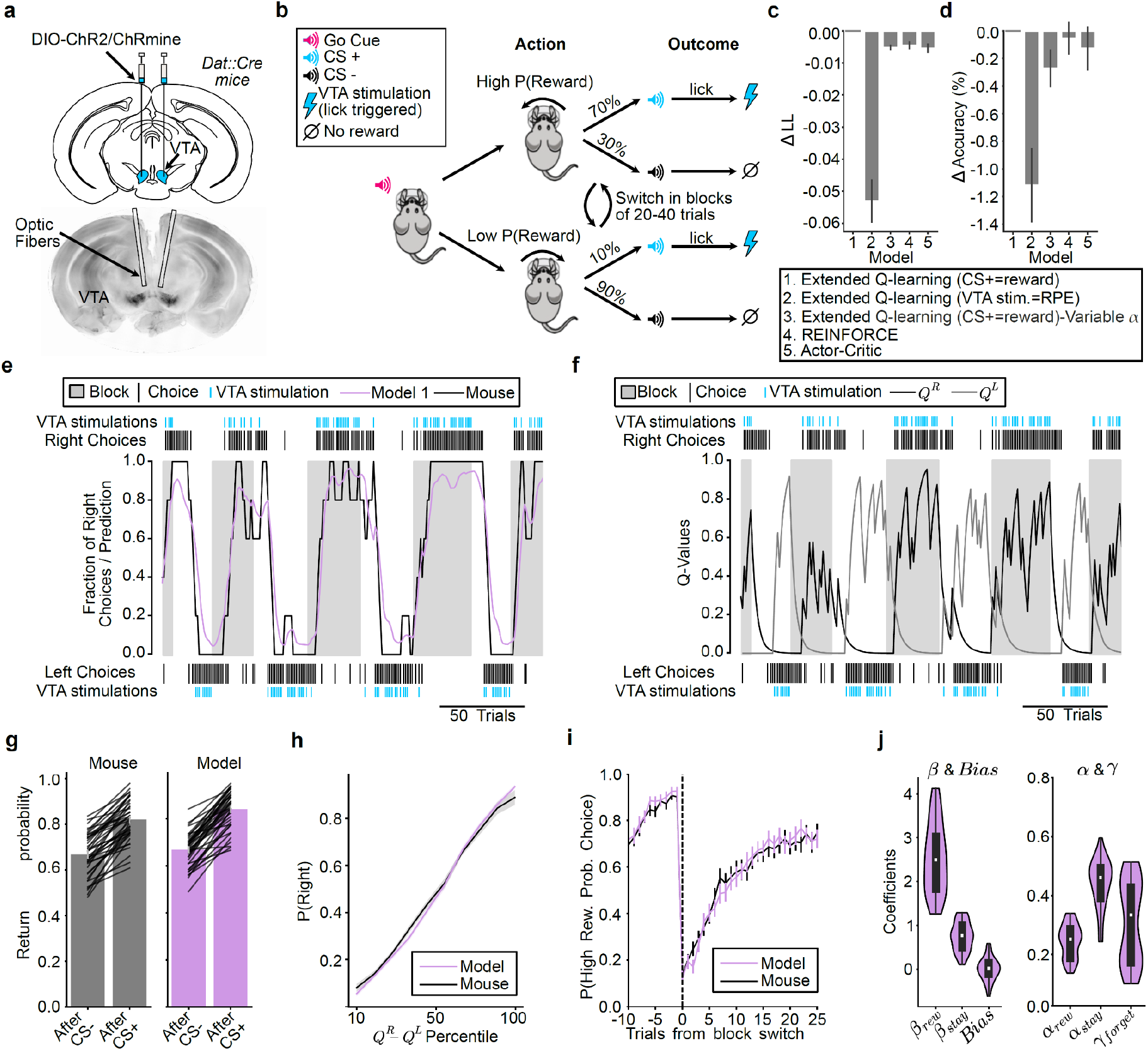
Mice adjust their behavior based on trial-and-error to obtain VTA DA stimulation. **a**, *Top:* Experimental strategy for optogenetic stimulation of VTA DA neurons. *Bottom:* Representative coronal brain slice showing virus expression and optical fiber placement. **b**, Probabilistic reversal learning task. VTA stimulation consisted of a 20 Hz burst of 5 ms light pulses for 1 s (ChR2: 447 nm, ∼8 mW; ChRmine: 532nm, ∼0.5 mW). **c**, Mean accuracy of different RL models in relation to the best performing extended Q-learning model (Model 1, Extended Q-learning (CS+ as a reward), 76.8% accuracy, -0.475 LL). Error bars represent mean ± s.e.m. across sessions (n=41 sessions). **d**, Same as c, but for log-likelihood (LL) per choice. **e**, Data and Q-learning model predictions in a representative behavioral session. The fraction of right choices (black line, 5-trial moving average) and the model’s predicted probability of choosing right (purple line, 5-trial moving average) are each plotted. Shaded grey areas indicate blocks of trials where the right choice had the higher probability of reward. Black vertical ticks at the top and bottom represent individual right and left choices. Blue vertical ticks indicate VTA DA stimulation. **f**, The Q-learning model’s action value estimates for the left (*Q*^*L*^) and right (*Q*^*R*^) choices over the same session as in (**e**). **g**, Calibration of the Q-learning model’s win-stay behavior. Bar plot showing the probability of repeating a choice following an CS+ or CS– trial for the mice (grey) and for the Q-learning model’s on-policy simulations (purple). Error bars represent the mean ± s.e.m. across sessions. **h**, Calibration of the Q-learning model’s action selection policy. The probability of choosing the right action is plotted as a function of the percentile of the difference between the model-derived action values (*Q*^*R*^ − *Q*^*L*^) for the mice (grey) and for the Q-learning model’s on-policy simulations (purple). Lines and shading represent the mean ± s.e.m. across sessions. **i**, Calibration of the Q-learning model’s reward-contingency reversal (block switch) behavior. The probability of choosing the high reward probability action is plotted as a function of trials from block switches for the mice (grey) and for the Q-learning model’s on-policy simulations (purple). Lines and error bars represent the mean ± s.e.m. across sessions. **j**, Distributions of the best-fit parameters for the Q-learning model across all sessions. ***β***_***Reward***_,: weight of the reward-seeking component of the decision, ***β***_*stay*_: weight of the perseveration component of the decision, : choice bias for left or right actions, *α*_***Reward***_,: learning rate for the reward-seeking component of the chosen action, ***α*_*stay*_**: : learning rate for the perseveration component, *γ*_*forget*_: forgetting rate for the reward-seeking component of the unchosen action. Violin plots illustrate the distribution of the mean of each parameter across sessions; the limits of the violin represent the extreme of the fit parameters, and the boxplot within the violin marking the median (white) and interquartile range (IQR; thick black line). *n* = 41 sessions from 8 animals for all panels. In panels **e-j**, results are from the best performing model: Model 1, Q-learning (CS+ as a reward)).

The task was optimized to produce ongoing value-based adaptation of choice behavior based on recent choices and their outcomes, but not to distinguish among the diverse family of reinforcement learning models^82^. To capture the animals’ decision-making strategy, we fit paradigmatic value-based (variants of Q-learning) and policy-based (REINFORCE and Actor-Critic) models to the choices (Fig. 1c,d; Extended Data Fig. 2). To be comprehensive, for Q-learning we considered a version where the CS+ acted as the reward (given that it signaled the availability of VTA DA stimulation^83^), a version where the VTA DA stimulation was the reward prediction error (given that DA is thought to signal a reward prediction error^1,6^), as well as a version with a variable learning rate^84,85^. All the models fit similarly (Fig. 1c,d, Extended Data Fig. 2a), produced similar trial-by-trial decision variables (Extended Data Fig. 2b), and, as expected, could not be distinguished from each other in a recovery analysis (Extended Data Fig. 2c,d). Therefore, for convenience, for subsequent analyses we used the single model which numerically fit best (Model 1: Extended Q-learning model (CS+ as reward), Extended Data Fig. 2a).

The model, which estimates the value of both the left (*Q*^*L*^) and right (*Q*^*R*^) actions to arrive at a choice, was able to accurately predict the animals’ choice trajectories across blocks (Fig. 1e,f). This model also successfully recapitulated key aspects of the animals’ decision-making behavior when run in simulation (“on-policy”), including the higher return probability after rewarded versus unrewarded choices (Fig. 1g), the relationship between relative value (Δ*Q = Q* ^*R*^ − *Q*^*L*^) and choice probability (*p*^(Right)^; Fig. 1h), and the gradual reversal in choice following block switches (Fig. 1i).

The fitted model coefficients (Fig. 1j; Extended Data Fig. 2e-h) revealed that the animals primarily relied on the reward (*β*_*rew*_), rather than perseveration (*β*_*stay*_) or bias to select their choice (Fig. 1j).Moreover, the animals integrated rewards over a number of trials, as evidenced by a slow learning rate (Fig. 1j; *α*_*rew*_= 0.245 ± 0.009; mean ± s.e.m.across sessions).

Together, this demonstrates that mice will perform a probabilistic reversal learning task to obtain VTA DA stimulation, and that their decision-making can be well-explained by a reinforcement learning model that estimates the trial-by-trial values of each possible action.

### VTA DA stimulation is sufficient to generate action value representations in the striatum, particularly in intermediate CP

We next set out to test the long-standing hypothesis that VTA DA neuron activity generates action value representations in the striatum^1,6,18,29,36,37,86,87^. To accomplish this, we surveyed neural activity across the striatum using high-density Neuropixels probes^88^ as mice performed our task (Fig. 2a and Extended Data Fig. 3a–e; 5,109 single units from 37 sessions from 8 mice).

**Figure 2.**
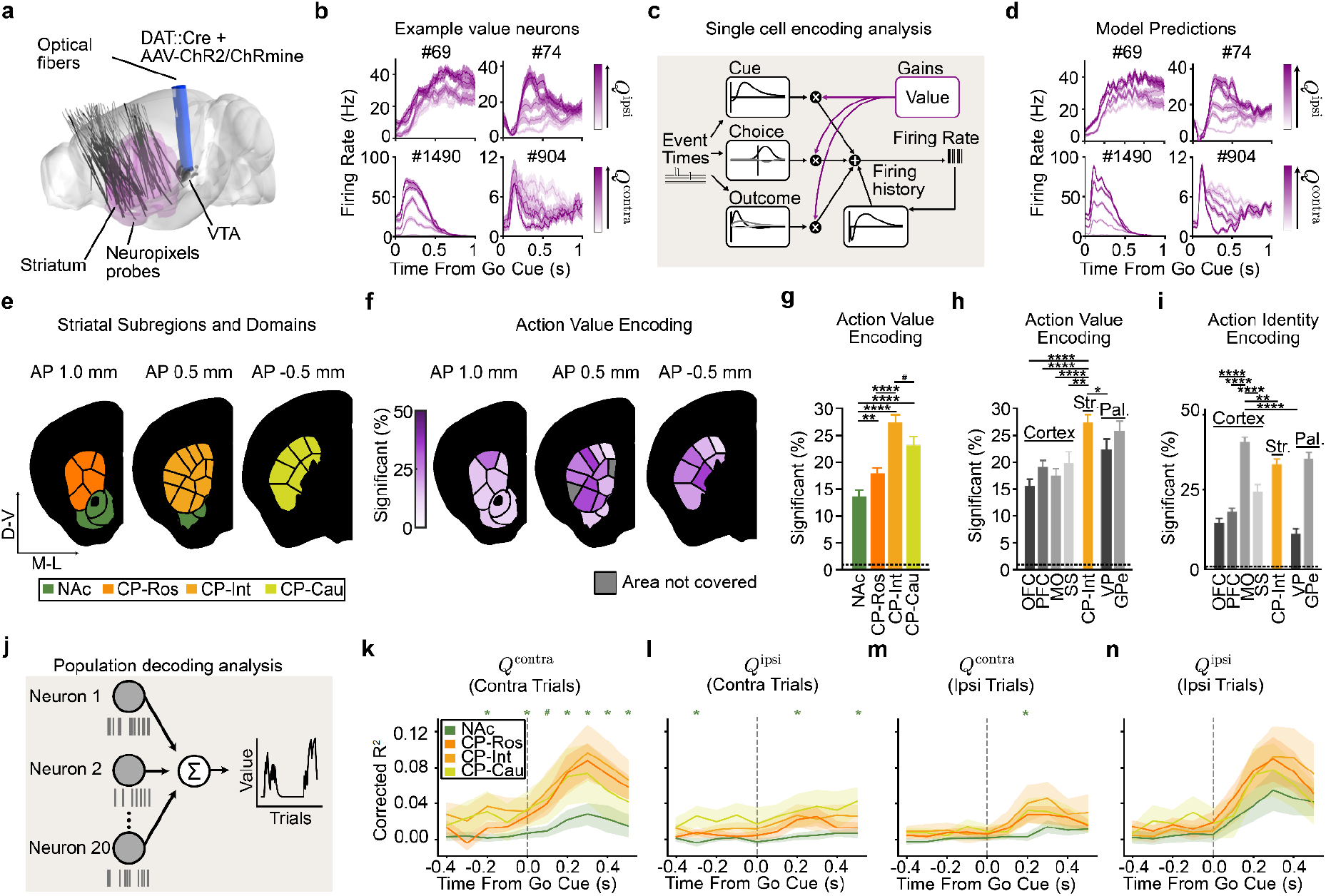
VTA DA stimulation is sufficient to generate action value representations, most prominently in the intermediate CP. **a**, 3D rendering of the Allen CCF showing optogenetic fiber (blue) placement over the VTA (black) and reconstructions of all Neuropixels probe trajectories (grey) in the striatum (purple) and associated brain structures. Recordings were bilateral but reconstructions are projected onto the left hemisphere. **b**, Example peri-stimulus time histograms (PSTHs) for four CP neurons (#69:CP-Int, #98:CP-Int, #1490:CP-Cau, #904:CP-Cau), showing their firing rate after the go cue. Trials are divided based on quintiles of *Q*^*ipsi*^(top) or *Q*^*contra*^(bottom) action values. Lines and shading represent the mean ± s.e.m. across trials. Only ipsilateral trials are plotted for *Q*^*ipsi*^ and only contralateral trials for . *Q*^*contra*^ **c**, Schematic of the single-neuron encoding analysis (see Methods for details). **d**, PSTH predictions from the encoding model for the same example neurons as in (**b**). **e**, Coronal sections illustrating the locations of analyzed striatal subregions and domains within three anterior-posterior (AP) subregions. Each striatal subregion (rostral, intermediate, and caudal CP and the NAc) has a unique color, and the striatal domains within those subregions are separated by black lines. **f**, Coronal sections identical to (**e**) with color saturation indicating the percentage of neurons significantly encoding action value in each striatal domain. Grey areas were not covered by the recordings. **g**, Percentage of neurons significantly encoding action values (either *Q*^*contra*^or *Q*^*ipsi*^) in recorded striatal subregions. **h**, Percentage of neurons significantly encoding action values (*Q*^*contra*^ or *Q*^*ipsi*^ in recorded cortical and pallidal subregions along with CP-Int from **(g).** *n* = 829 NAc, 1529 CP-Ros, 958 CP-Int, 720 CP-Cau, 733 OFC, 996 PFC, 1088 MO, 402 SS, 522 VP, and 615 GPe task-modulated single units, see Extended Data Table 3 for mapping between these names and Allen CCF regions. **i**, Same as (**h**) but for action identity encoding (choice). **j**, Schematic of the population decoding analysis (see Methods for details). **k–n**, Time-resolved population decoding performance (corrected R^2^) in 100 ms bins for *Q*^*contra*^ (**k,m**) or *Q*^*ipsi*^ (**l,n**) action values in trials with contralateral (**k,l**) or ipsilateral trials (**m,n**) choices. Decoding was based on groups of 20 randomly chosen neurons. Lines and shading represent the mean ± s.e.m. across sessions (*n* = 18 NAc, 20 CP-Ros, 15 CP-Int, and 13 CP-Cau sessions). The corrected R^2^ is computed by subtracting the mean R^2^ of a session-specific null distribution from the mean R^2^ of decoders applied to the observed behavioral data (see Methods for details). * indicate time points where decoding performance for CP was significantly different from NAc (*P* ≤ 0.05, from two-tailed Mann–Whitney U test at each timepoint). In all bar plots, bars denote the mean ± bernoulli s.e.m. across all recorded neurons and the dashed lines indicate the chance level for significance (1%; see Methods for details on significance testing). In g-h *P < 0.05, **P < 0.01, ***P < 0.001, ****P < 0.0001 from χ^2^ tests highlighted in the results, for all comparison see Extended Data Table 7.

The activity of most striatal units was significantly modulated by task events (79.0%). Of these task-modulated neurons, 67.3% were modulated by the go cue, 61.0% by the actions, and 87.0% by the outcomes (*P* ≤ 0.01 based on a linear encoding model; see Methods for details; Extended Data Fig. 3f–k).

For a subset of these neurons, the responses to these task events were further modulated by trial-by-trial action values (as estimated from the Q-learning model). For example, the event-related firing of the example neurons shown in Fig. 2b increased or decreased monotonically as a function of contralateral (*Q*^*contra*^) or ipsilateral (*Q*^*isit*^) action value (contralateral and ipsilateral are defined relative to the recording hemisphere, see Methods).

This implies that VTA DA neuron stimulation is indeed sufficient to create action value representations in at least a subset of striatal neurons. Our high-density Neuropixels recordings allowed us to map where in the striatum such DA-driven action value representations were located.

First, to quantify if and how each neuron’s event-related activity was modulated by the trial-by-trial *Q*^*contra*^ and *Q*^*ipst*^ action values, we used a bilinear encoding model^42,80^ (Fig. 2c,d; see Methods for details). Briefly, each neuron’s spiking activity was modeled using a sum of temporal kernels aligned to the task events (go cue, contralateral/ipsilateral action (relative to the recording hemisphere), CS+/–, and VTA DA stimulation) that were each modulated multiplicatively by trial-by-trial action values inferred by the behavioral model. We then quantified the strength and significance of action value coding by calculating how much the inclusion of the action values in the model improved our ability to explain each neuron’s spiking activity relative to null distributions that preserved the temporal correlations in the data^89,90^ (see Methods for details). Across the entire striatum, our encoding model identified 20.2% of neurons as significantly encoding trial-by-trial action values.

We then examined the strength of action value encoding across the striatum based on the divisions from the mouse cortico-striatal projectome^91,92^, which organizes the striatum into four higher-order divisions (the rostral, intermediate, and caudal CP and the NAc; referred to here as “subregions”) that are then further parcellated into thirty-one lower-order subdivisions (referred to here as “domains”; Fig. 2e).

Within the four subregions, we found that modulation by action value was most common in the intermediate CP (27.3% of neurons) and least common in the NAc (13.6% of neurons; Fig. 2f,g; similar conclusions from ΔR^2^ in Extended Data Fig. 4a). Intermediate CP had significantly more neurons encoding action value than NAc (*P* < 1e-11 by χ^2^ test) and rostral CP (*P* < 1e-7 by χ^2^ test) with a trending effect relative to caudal CP (*P* = 0.054 by χ^2^ test; *P* < 1e-6 when comparing ΔR^2^ by Mann–Whitney U test). In contrast, the NAc had fewer neurons encoding action value than any other subregion (*P* < 0.01 vs. CP-Ros, *P* < 1e-11 vs. CP-Int, and *P* < 1e-5 vs. CP-Cau by χ^2^ tests). Furthermore, within the intermediate CP, individual domains had as many as 35.9% of neurons significantly encoding action value (Fig. 2f; see also Extended Data Fig. 4b–e).

The stronger value coding in intermediate CP than NAc was the case despite firing rates, and therefore the statistical power of our encoding model^93,94^, being significantly higher in the NAc when compared to the intermediate CP (*P* < 0.001 by Mann–Whitney U test; Extended Data Fig. 3e).

### Stronger encoding of action identity in MO but action value in intermediate CP

We compared these encoding model results in striatum to those in cortex, because the cortex provides a major monosynaptic input to the striatum (Fig. 2h). We performed cortical recordings in the somatosensory cortex (SS), motor cortex (MO), prefrontal cortex (PFC), and orbitofrontal cortex (OFC; see Extended Data Table 3 for Allen region groupings). In the cortex, neurons encoding action values were less common compared to the striatal subregion where action value was most strongly encoded (intermediate CP; *P* < 1e-8 vs. OFC, *P* < 1e-4 vs. PFC, *P* < 1e-6 vs. MO, and *P* < 0.01 vs. SS by χ^2^ tests). In contrast, the action identity (contralateral versus ipsilateral choice) was most strongly encoded in the MO compared not only to other cortical regions (*P* < 1e-7 vs. OFC, PFC, and SS by χ^2^ tests) but also to all striatal subregions, including the intermediate CP (*P* < 0.01 by χ^2^ test) (Fig. 2i). Thus, action encoding dominates in MO while action value encoding dominates in intermediate CP.

We also recorded from output structures of the CP and NAc: the globus pallidus externus (GPe) and ventral pallidum (VP), respectively (Fig. 2h). In the GPe, we observed a high fraction of neurons encoding action value (*P* = 0.51 by χ^2^ test) and action identity (*P* = 0.50 by χ^2^ test), at levels that were comparable to the intermediate CP. In contrast, the fraction of neurons encoding action value (*P* < 0.05 by χ^2^ test) or action identity (*P* < 1e-15 by χ^2^ test) was significantly lower in the VP when compared to the intermediate CP . These results suggest that value representations in the striatum are projected downstream to the pallidum.

### Population analyses corroborate stronger representations of action value in CP relative to NAc

The analyses above relied on single neuron encoding models. Population-level decoding corroborated the conclusion that action value representations generated by VTA DA stimulation are preferentially localized to the CP relative to the NAc. We trained linear decoders to predict the trial-by-trial estimates *Q*^*contra*^ of *Q*^*ipsi*^ or using the activity of groups of 20 randomly chosen, simultaneously recorded units from each striatal subregion around the time of the go cue (Fig. 2j–n and Extended Data Fig. 4f–i). In order to control for neural activity related to the different actions, we trained separate action value decoders for contralateral versus ipsilateral choices.

On trials in which the mouse turned the wheel in the contralateral direction, the value of both chosen (contralateral) and unchosen (ipsilateral) actions was better decodable from CP subregions than from the NAc in multiple 100 ms bins before and after the go cue (Fig. 2k,l; *P* < 0.05 by Mann–Whitney U tests). For both contralateral and ipsilateral choices, the value of non-chosen actions was less strongly decoded than the chosen action (compare Fig. 2k to Fig. 2l, and Fig. 2m to Fig. 2n; statistics in Extended Data Table 7), although decoding of the non-chosen value was still above chance across CP subregions (Fig. 2l). Decoding in the CP tended to be stronger than NAc on ipsilateral choice trials as well, although the difference was not significant in this case (Fig. 2m,n).

Downstream of the striatum, action value decoders trained using population activity from the GPe, a major output of the CP, performed much better than decoders trained using activity from the VP, an output of the NAc (Extended Data Fig. 5a–d). Indeed, action value decoding in the VP was hardly better than chance, despite the presence of single neurons that significantly encoded this variable (Fig. 2h). Thus, action value decoding is stronger in the CP relative to the NAc, as well as in the respective downstream targets of these structures.

### VTA DA stimulation generates relative and state value representations, most prominently in intermediate CP

We were surprised that action value representations generated by VTA DA stimulation were localized to the CP (relative to the NAc; Fig. 2), given that the NAc is the major target of DA release by VTA DA neurons^95,96^, a fact which we confirmed in our own stimulation experiments (Extended Data Fig. 6a–e).

However, so far we have only considered modulation of neural activity by the value of each action. A prominent hypothesis (the Actor-Critic model^37,48,87^) instead suggests that the NAc might compute a different form of value: state value (the trial-by-trial reward expectation,*V*), which could be used for calculating reward prediction errors and also serve as a motivational signal. The CP, in turn, might encode relative value (the difference in value between the two actions, Δ*Q*), which could be used to select a choice.

Therefore, we next investigated whether state and relative values were differentially encoded across striatal subregions and domains, and whether state value would be preferentially localized to the NAc. To this end, we again used a bilinear encoding model to test how strongly each neuron’s task-related activity was modulated by trial-by-trial estimates of value, but this time considered modulation by relative value (Δ*Q* = *Q*^*contra*^ − *Q*^*ipsi*^) or state value (*V* ; Fig. 3a), rather than *Q*^*contra*^ or *Q*^*ipsi*^) (Fig. 2c–h). We identified many neurons whose activity was correlated with one or both of these variables (example neurons: Fig. 3b,c).

**Figure 3.**
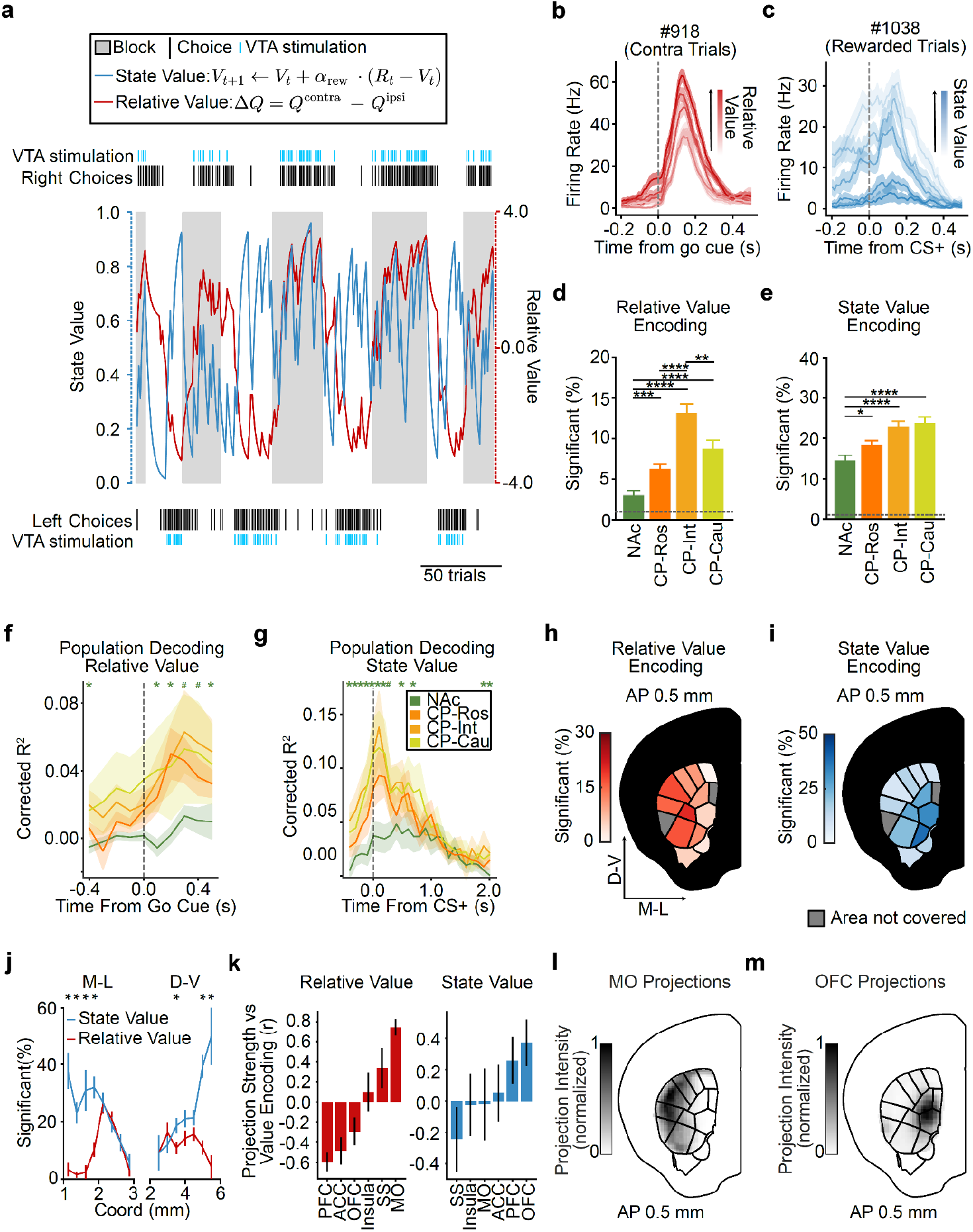
VTA DA stimulation generates relative and state value representations in the intermediate CP, with different spatial organizations. **a**, Estimated state (blue) and relative (red) value for a representative behavioral session. Black vertical ticks at the top and bottom represent individual right and left choices. Shaded grey indicates blocks where the right choice had the higher probability of reward. Blue vertical ticks indicate VTA stimulation. **b**, Example PSTH for a striatal neuron (#918:CP-Ros) in contralateral trials for different relative values. **c**, Example PSTH for a different striatal neuron (#1038: CP-Int) for rewarded trials with different state values. In (**b,c**), trials are divided based on value quintiles of relative and state value. Lines and shading represent mean ± s.e.m. **d**, Percentage of neurons in each striatal subregion significantly encoding relative value. Bar plots represent mean ± s.e.m. (*n* = 829 NAc, 1529 DS-Ros, 958 DS-Int, 720 DS-Cau). The dashed line indicates the chance level for significance (1%). *P < 0.05, **P < 0.01, ***P < 0.001, ****P < 0.0001 from χ^2^ tests highlighted in the results, for all comparison see Extended Data Table 7. **e**, Same as (**d**) but for state value. **f**, Time-resolved population decoding performance (corrected R^2^) for relative value in trials with contralateral choices (based on groups of 20 randomly chosen neurons). The corrected R^2^ is computed by subtracting the mean R^2^ of a session-specific null distribution from the mean R^2^ of decoders applied to the observed behavioral data (see Methods for details). Lines and shading represent the mean ± s.e.m. across sessions (*n* = 18 NAc, 20 CP-Ros, 15 CP-Int, and 13 CP-Cau sessions). * indicates time points where decoding performance for CP was significantly different from NAc (*P* ≤ 0.05, from two-tailed Mann–Whitney U test at each timepoint). **g**, Same as (**f**) but for state value around the time of the CS+, on rewarded trials only. **h**, Coronal sections illustrating analyzed striatal domains within CP-Int. Saturation indicates the local percentage of neurons significantly encoding relative value. Grey areas were not covered by the recordings. **i**, Same as **(h)** but for state value encoding. **j**, Average percentage of neurons encoding state (blue) and relative (red) value along the mediolateral (left, ML) and dorsoventral (right, DV) axes within intermediate CP. Lines and error bars represent mean ± binomial s.e.m. (\**P* < 0.05, χ^2^ test) **k**, Correlation coefficients (Pearson r) between the normalized cortical projection intensity to each striatal domain from the Allen Connectivity Atlas^97^ and that domain’s encoding of relative (red) and state (blue) value. Bar plots indicate mean ± bootstrap standard deviation (s.d.). From left to right: cortical projection intensities were obtained from 9 pre-frontal cortex (PFC), 34 anterior cingulate cortex (ACC), 16 orbito-frontal cortex (OFC), 16 Insula, 70 somatosensory cortex (SS), and 60 motor cortex (MO) anterograde injections (see Extended data table 6 for details). **l,m**, Mean normalized intensity of projections from primary and secondary motor cortex (**l**, MO) and orbito-frontal cortex (**m**, OFC) to intermediate striatum.

Similar to *Q*^*contra*^ and *Q*^*ipsi*^ (Fig. 2), relative (Fig. 3d and Extended Data Fig. 7a–c,g) and state (Fig. 3e and Extended Data Fig. 7d–f,h) values were preferentially encoded in the CP relative to the NAc. Intermediate CP had significantly more neurons encoding relative value than all other striatal subregions (*P* < 1e-13 vs. NAc, *P* < 1e-8 vs. CP-Ros, and *P* < 0.01 vs. CP-Cau by χ^2^ tests), whereas NAc had fewer neurons encoding relative value than any other subregion (*P* < 0.001 vs. CP-Ros, *P* < 1e-13 vs. CP-Int, and *P* < 1e-5 vs. CP-Cau by χ^2^ tests; Fig. 3d). A similar proportion of neurons significantly encoded state value in intermediate and caudal CP (*P* = 0.67 by χ^2^ test; Fig. 3e), however the magnitude of this state value coding, based on the explained variance (ΔR^2^) of neural activity across the entire population, was significantly stronger in intermediate CP (*P* < 0.01 by Mann–Whitney U test; Extended Data Fig. 7e). Once again, NAc had fewer neurons encoding state value than any other striatal subregion (*P* < 0.05 vs. CP-Ros and *P* < 1e-5 vs. CP-Int and CP-Cau by χ^2^ tests; Fig. 3e). This was true also when considering other methods of calculating state value (based on the sum or probability-weighted sums of *Q*^*contra*^ and *Q*^*ipsi*^); Extended Data Fig. 7n,o).

We next trained population-level decoders for both relative and state value (Fig. 3f,g and Extended Data Fig. 7i,j). Relative value decoders were trained using only contralateral trials to control for representations of the action itself, while state value decoders were trained using only rewarded trials to control for the response to the outcome itself. These population-level analyses supported the preferential localization of both relative (Fig. 3f and Extended Data Fig. 7i) and state (Fig. 3g and Extended Data Fig. 7j,p,q) values to the CP in comparison to the NAc. Moreover, state value decoding in the CP was apparent earlier in the trial than in the NAc (from the first timepoint tested, 500–400 ms before CS+ onset; Fig. 3g). Thus, not only action values (*Q*^*contra*^ and *Q*^*ipsi*^), but also relative and state values (Δ *Q* and *V*), are most prominently represented in the CP relative to the NAc.

Within the intermediate CP, relative and state value representations had distinct spatial organizations. Specifically, relative values were more prominently encoded in dorsolateral domains of the intermediate CP, whereas state values were more strongly represented in ventromedial domains (Fig. 3h–j and Extended Data Fig. 7k–m). To investigate how these spatial patterns relate to the topography of cortical inputs to the striatum, we turned to the Allen Projection Atlas^97^ (Fig. 3k–m). Interestingly, the encoding of relative value — a variable which could be used for action selection — was positively correlated with the projection pattern from the MO (*r* = 0.715 Pearson correlation; Fig. 3k,l). The MO is of course important for generating actions^98–108^ and, in our data, was the cortical region that best encoded the animals’ choice (Fig. 2i). In contrast, the encoding of state value was positively correlated with the OFC projection pattern (*r* = 0.317 Pearson correlation; Fig. 3k,m), a cortical region implicated in value^38,39,109^ and state^110–112^ representations.

### Value coding is strongest in intermediate CP across learning and forgetting rates

We next explored the hypothesis that a mismatch in the timescale of value computation between the NAc and behavior could potentially explain why value representations appear relatively weak in the NAc (Figs. 2,3). Action values depend on parameters that determine the timescale over which previous rewards are integrated (learning rate: α_*rew*_Fig. 4a) and forgotten (forgetting rate: γ_*forget*_; Fig. 4b). We so far have considered neural correlates of action values (Figs. 2,3) based on the values of these parameters that best explain the animals’ behavior (Fig. 1). However, previous fiber photometry recordings suggest that DA release in the NAc and CP may reflect lower and higher learning rates, respectively^62^. Inspired by this result and other findings about temporal scales across the dopamine system^67,113,114^, we wondered whether individual neurons in the NAc may in fact integrate (or forget) rewards more slowly than the animals’ behavioral policy — and than neurons in the CP.

**Figure 4.**
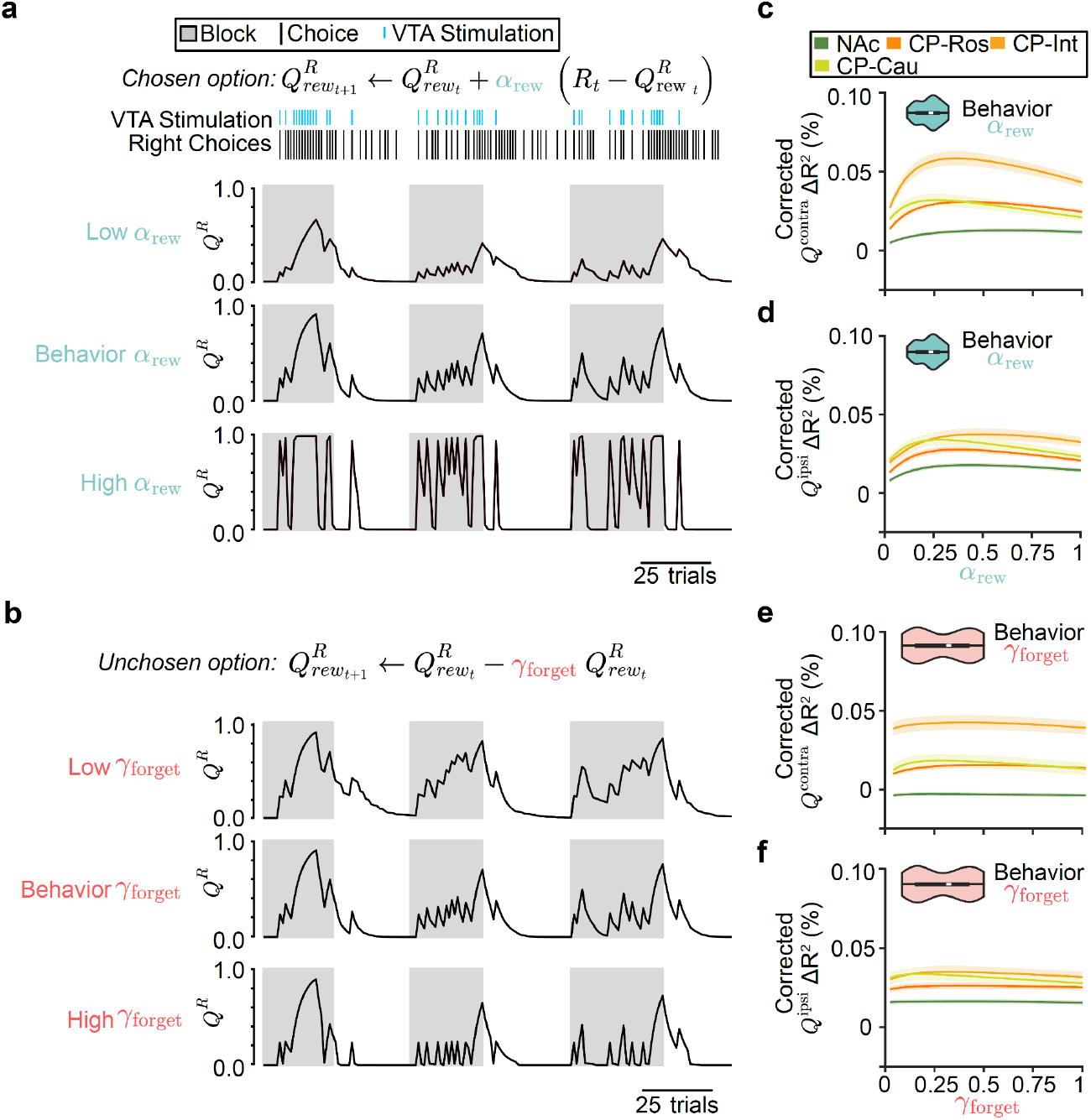
Across learning and forgetting rates, action value coding is strongest in intermediate CP. **a**, Right action values (*Q*^*R*^) of an example session for the behaviorally fit learning rate (*α*_*rew*_, middle panel), a high learning rate (0.95; bottom panel), and a low learning rate (0.1; top panel). Black vertical ticks represent individual right choices. Blue vertical ticks indicate VTA stimulation. Shaded grey areas indicate blocks where the right choice had the higher probability of reward. **b**, Same as (**a**) but for forgetting rates (*γ*_*forget*_). **c**, Average change in corrected explained variance (corrected ΔR^2^) for the single-neuron encoding model as a function of the learning rate parameter (*α*_*rew*_) for . *Q*^*contra*^ Colored lines indicate different striatal subregions. Violin plots illustrate the distribution of *α*_*rew*_ obtained from fitting the behavior (plot limits described in Fig. 1j) . Corrected ΔR^2^ computed by subtracting the mean ΔR^2^ of pseudosession-based null distribution from the ΔR^2^ of the encoding model applied to the observed behavioral data (see Methods for details). **d**, Same as (**c**) but for *Q*^*ipsi*^, action values. **e,f**, Same as (**c,d**) but for the forgetting parameter (*γ*_*forget*_) instead of *α*_*rew*_ Lines and shading represent mean ± s.e.m. across sessions.

To test this idea, we asked what learning rate best explains individual neurons’ activity across the striatum. To do this, we refit encoding models to the neural activity using trial-by-trial action values for different learning rates (α_*rew*_ varying between 0 and 1). This revealed that the CP — especially the intermediate CP — more strongly encoded both contralateral and ipsilateral action values across all learning rates (Fig. 4c,d and Extended Data Fig. 8a,b).

Within the CP, in particular for contralateral action value (Fig. 4c), the learning rate that best explained trial-by-trial variations in neural activity was similar to the learning rate that best described the animals’ choice behavior (α_*rew*_ = 0.245 ± 0.009; mean ± s.e.m. across sessions). We repeated the same analysis for relative and state value, instead of action values, and observed the same phenomenon (Extended Data Fig. 8e–g).

We also performed an equivalent analysis in which we varied the forgetting rate (γ_*forget*_;), rather than the learning rate (Fig. 4b). Again, we found that the CP more strongly encoded both contralateral and ipsilateral action values across all potential forgetting rates (Fig. 4e,f and Extended Data Fig. 8c,d). However, we observed that varying the Q-learning model’s forgetting rate had very little impact on the neural encoding of action values^115^, implying that learning rate is better reflected in neural activity than forgetting rate.

Together, these analyses suggest that the relatively weak DA-driven action value correlates in the NAc is not because neurons in this part of the striatum are representing value using a slower learning rate than the animal’s behavioral policy. Instead, across learning and forgetting rates, value coding is strongest in the intermediate CP, with neurally estimated learning rates roughly matching the behaviorally estimated learning rates.

### VTA DA stimulation endows the CS+ with value, which the mice then work for in our task

So far, we have shown that across different types of value and different learning and forgetting rates, VTA DA stimulation produces value representations that are strongest in the intermediate CP. The fact that these representations are strongest in this subregion — which is not the major direct target of the stimulated VTA DA neurons^95,96^ (Extended Data Fig. 6a–e) — suggests an indirect path between VTA DA stimulation and the formation of action value representations.

What could mediate an indirect path between VTA DA stimulation and action values? We hypothesized that rather than directly producing action values, this stimulation could contribute to learning the value of a stimulus^28,29,116,117^ — the CS+ in our task — and this valuable stimulus could in turn serve as a conditioned reinforcer to produce valuable actions. This hypothesis was inspired by previous work on conditioned reinforcement, in which repeatedly pairing a stimulus with DA stimulation in the NAc (but not the CP) leads to the stimulus itself being able to reinforce actions^28^, as well as the observation that a sensory stimulus can potentiate self-stimulation of VTA DA neurons^118^.

This hypothesis makes two core predictions, which we test below: 1) That even though action value representations are relatively sparse in the NAc, responses to the CS+ will be readily apparent; and 2) That the mice are working for the CS+ itself in our task, and not directly for the VTA DA stimulation.

Indeed, even though the NAc had relatively few neurons significantly encoding action values (13.6%, Fig. 2f,g), encoding of the CS+ was very common (55.9%; see Methods for details). Based on the kernels of our encoding model, which distinguish neural responses between task events (including the CS+ from VTA DA stimulation), the plurality of NAc neurons (35.7%) were most strongly activated by the CS+ relative to other task events (Fig. 5a). This predominant response to the CS+ was widespread across striatum: the plurality of neurons in all subregions were most strongly activated by the CS+ (Fig. 5b and Extended Data Fig. 9a–c).

**Figure 5.**
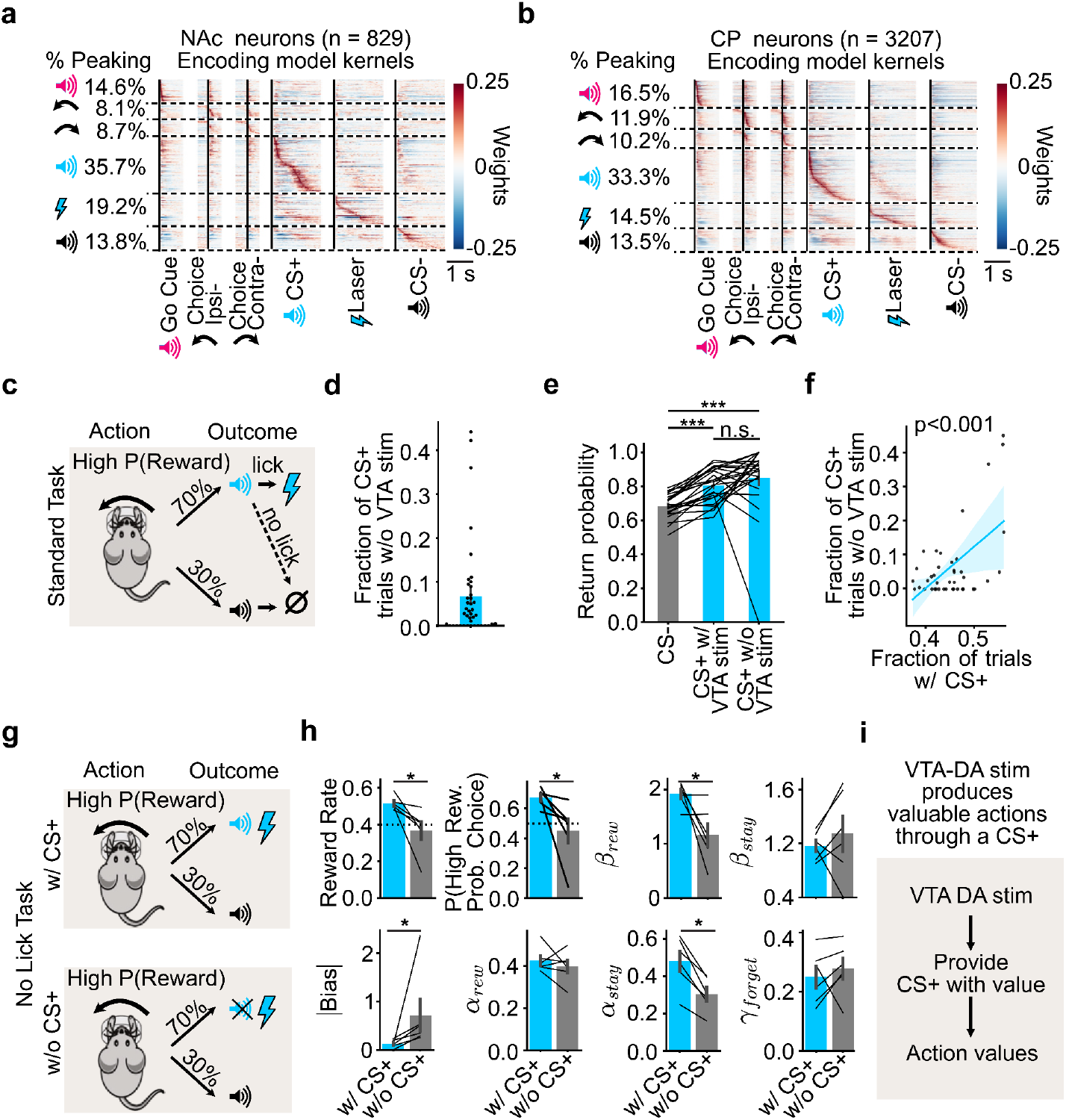
VTA DA stimulation endows the CS+ with value, which the mice then work for in our task. **a**, Encoding model kernels for individual neurons in the NAc, with neurons sorted by the peak across their kernels. “% peaking” on the left represents the fraction of neurons with the peak kernel response for each task event. **b**, Same as (**a**) but for CP. **c**, Schematic of the outcome contingency in the probabilistic reversal learning task, highlighting the licking requirement for the acquisition of the VTA stimulation. **d**, Fraction of CS+ trials without VTA DA stimulation. Individual points represent sessions. *n* = 41 sessions from 7 mice. **e**, Average probability of repeating the previous choice (left or right) as a function of trial outcome. Individual lines represent sessions where there were at least 5 CS+ trials without VTA DA stimulation (***P < 0.001, paired Wilcoxon signed-ranked test with Bonferroni-Holm’s correction). *n* = 24 sessions from 7 mice. **f**, Correlation across sessions between the fraction of CS+ trials without VTA stimulations (CS+ trials without VTA stimulations/total CS+ trials) and overall performance (CS+ trials/total trials). Line represents linear regression. *n* = 41 sessions from 8 mice. **g**, “No Lick” task for examining the role of the CS+ on VTA DA seeking behavior, in which the licking requirement is removed and the CS+ and VTA DA are presented simultaneously (w/ CS+), or else the CS+ is omitted (w/o CS+). **h**, On sessions with and without the CS+ presentation, bar plots of the average reward rate, fraction of high reward probability choices (P(high reward probability choice)), and behavioral model parameters (*β*_*rew*_,*β*_*stay*_, *Bias α*_*rew*_,*α*_*stay*_, *γ*_*foregt*_). Lines represent individual mice (\**P* < 0.05,. n = 6 animals. **i**, Overall, our data points towards a model where VTA DA stim produces action values indirectly, by first providing a CS+ with value (i.e., making a “conditioned reinforcer”), which is then used to generate action value representations in the CP. All bar plots show mean ± s.e.m. n.s. signifies p>0.05. See Extended Data Table 7 for statistical details

Next, we tested the idea that mice in our task are working for the CS+ itself, rather than for VTA DA stimulation. Upon hearing the CS+ on rewarded trials, VTA DA stimulation must be “collected” by licking a dry spout (Fig. 5c,d). We hypothesized that if animals were working for the CS+, and not for VTA DA stimulation itself, then they might at times forgo collecting the VTA DA stimulation and that, when doing so, their subsequent choice behavior would be unaffected. We found support for this hypothesis, following CS+ trials with versus without VTA DA stimulation, the animals’ behavior was similar, which implies that VTA DA stimulation did not impact their next trial behavior. This was true for the probability of repeating a choice (Fig. 5e) as well as the next trial latency (a readout of motivation; Extended Data Fig. 10a). Moreover, in the sessions with the highest performance, mice tended to have a lower VTA DA stimulation collection rate (*P* < 0.001; Fig. 5f). This may also suggest that high-performing mice may indeed be working to receive the VTA DA stimulation-paired CS+, rather than the stimulation itself.

The fact that behavior was similar after a CS+ trial regardless of whether VTA DA stimulation was received (Fig. 5f) suggests that action values may be updated in a similar manner across these outcomes. To test this, we re-fit our Q-learning model, this time leaving the value of the CS+ without VTA DA stimulation as a free parameter. If the CS+ is rewarding (without VTA DA stimulation), we would expect the fitted value of this parameter to be positive. Furthermore, if its value is similar to the value of the CS+ with VTA DA stimulation (which is preset to 1), then we would expect this parameter to be close to 1 as well. We found that this was indeed the case. The fitted value of the CS+ without VTA DA stimulation was positive and extremely close to 1 both on average (Extended Data Fig. 10b) and on individual sessions (Extended Data Fig. 10c).

Together, these analyses indicate that mice update their choice behavior and action values equally after CS+ trials regardless of whether VTA DA stimulation was actually collected. This suggests that the animals were working for the CS+ and not for VTA DA stimulation itself.

To test this idea more directly, we next examined the effect on performance of experimentally removing the CS+. We trained a separate cohort of mice on a variant of our task in which VTA DA stimulation and the CS+ co-occured immediately after the choice (i.e., we removed the licking requirement to obtain VTA DA stimulation so it began simultaneously with the CS+). Then, we compared performance on sessions with and without the CS+ (“No Lick Task with CS+” versus “No Lick Task without CS+”; Fig. 5g). Removing the CS+ led to a dramatic reduction in task performance, which was evident both from the animals’ reward rates and from the estimated weight of reward-seeking (β_*rew*_) in our behavioral model (*P* < 0.05 by Mann–Whitney U test; Fig. 5h). Note that the CS+ and the VTA DA stimulation occurred simultaneously in this task variant, and therefore the importance of the CS+ to performance cannot be attributed to a shorter or stronger temporal relationship between the action and the CS+ (compared to the VTA DA stimulation).

Thus, our behavioral data (Fig. 5c–h) are consistent with a model in which mice are working for the VTA DA stimulation-associated CS+ rather than for VTA DA stimulation itself. Together with our neural data (Figs. 2–4 and Fig. 5a,b), this suggests that VTA DA activity may support action learning indirectly. First, DA release (potentially in the NAc) endows stimuli with value as animals discover which stimuli are temporally associated with VTA DA neuron activation. Then, these valuable stimuli in turn produce action value representations in downstream sites, such as the intermediate CP (Fig. 5i).

### A new Conditioned Reinforcer model quantitatively explains behavioral sensitivity to the CS+

Our behavioral and neural results suggest that VTA DA stimulation may support action learning indirectly, by updating the value of the CS+, which in turn serves to update the value of actions that lead to the CS+ (“conditioned reinforcement”). To interrogate this hypothesis further, we built a behavioral model that instantiates such a conditioned reinforcement framework to test if it could quantitatively capture the role of the CS+ in behavioral updating. Moreover, by explicitly modeling the CS+ valuation process, we could also determine if CS+ value is preferentially encoded at the primary target site of VTA DA neurons (NAc), which would support a direct role for these neurons in updating the CS+ value.

The striking finding that the mice could not do the task without the CS+, even if VTA DA stimulation was presented immediately after the wheel turn (Fig. 5g,h) implies that a traditional Actor-Critic model, with a dopaminergically-coded TD error that updates the actor, would not be able to model our data Extended Data Fig. 11a). Thus we built a new model, called the “Conditioned Reinforcer” model (Fig. 6a) that comprises two interconnected Actor-Critic systems. One Actor-Critic calculates the value of the CS state and controls the licking decision, and the other calculates the value of the choice state and the decision to turn the wheel left or right. The CS critic uses VTA stimulation as its error signal. The choice critic uses an error signal calculated by the CS critic.

**Figure 6.**
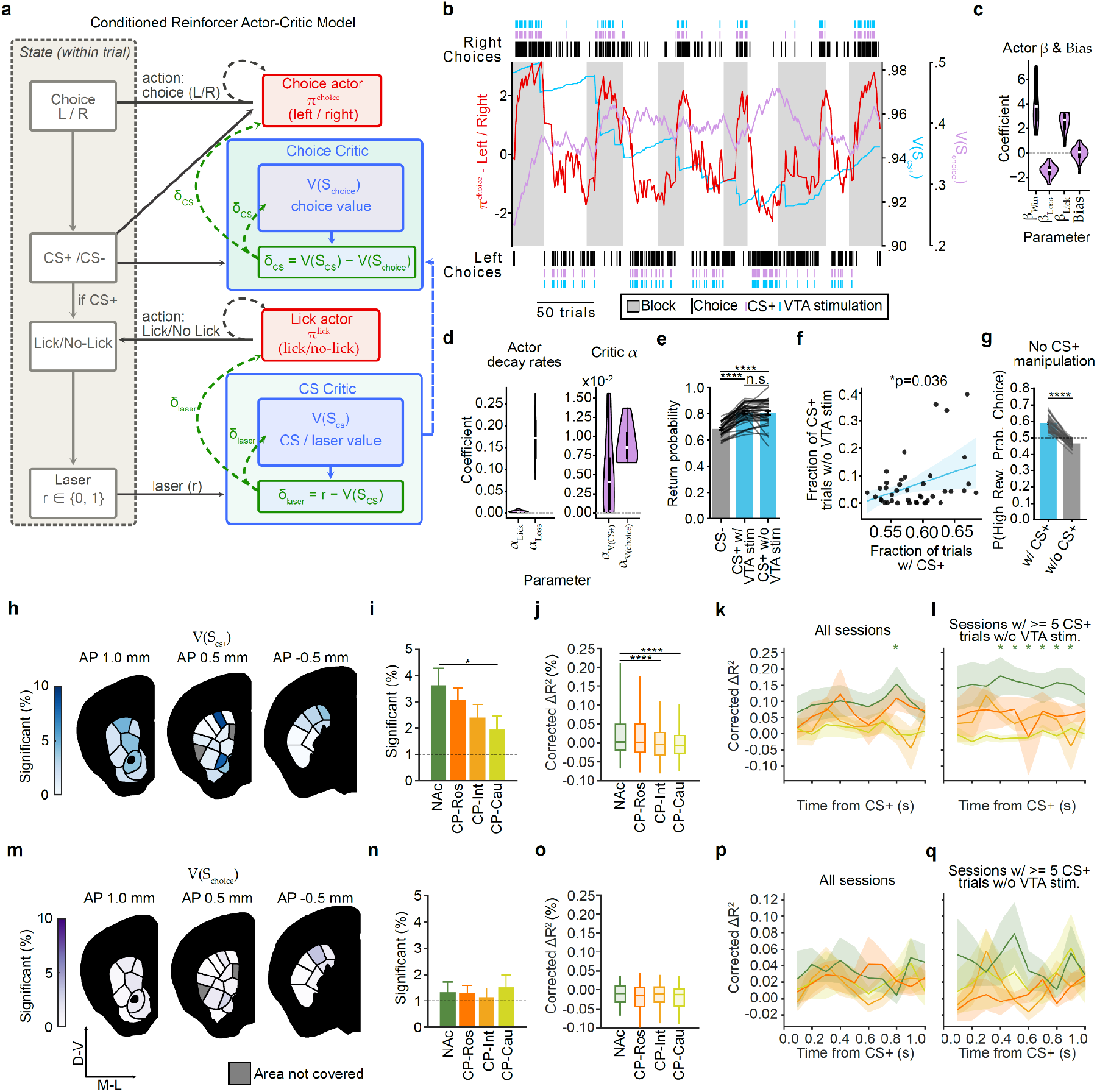
The Conditioned Reinforcer model reproduces the dependence of behavior on the CS+ by computing a CS+ value that is preferentially represented in NAc. **a**, Schematic of the Conditioned Reinforcer model (see Methods for details). Within a trial, the animal transitions through a choice state (*S*_*choice*_), an outcome state signalled by the CS+ or CS– (*S*_*cs*_), and, on CS+ trials, a lick / no-lick state gating delivery of the VTA DA stimulation (r ∈ {0,1}). The value of the CS is estimated by the CS critic (blue). On CS − trials,*V* (*S*_*cs*_) = *V*(*S*_*cs-*_) =0; on CS+ trials, *V* (*S*_*cs*_) = *V*(*S*_cs+_, is a learned quantity. The CS+ acquires its value from the stimulation itself, via the prediction error *δ*_*laser*_.=r − *V*(*S*_*cs+*_) The same error trains the lick actor (red), *π*^*lick*^, so that licking is reinforced where it obtains stimulation. Having acquired value, the CS+ is a conditioned reinforcer and can act on what precedes it. Earlier in the trial the choice actor (red) selects the left or right port, *π*^*choice*^, and the choice critic (blue) estimates *V*(*S*_*choice*_). Both are updated by*δ*_*cs*_ =V(*S*_*cs*_−*V*(*S*_*choice*_), so choices followed by the CS+ are reinforced and those followed by the CS− are not. **b**, Model variables for a representative behavioral session: *π*^*choice*^(red), *V*(*S*_*cs+*_) (cyan) and *V(S_choice_)* (purple). Black vertical ticks at the top and bottom represent individual right and left choices, magenta ticks indicate CS+ trials and blue ticks indicate VTA DA stimulation. Shaded grey areas indicate blocks of trials where the right choice had the higher probability of reward. **c**, Distributions of the best-fit parameters across sessions: actor weights and bias (*β*_*win*_, *β*_*Loss*,_ *β*_*Lick*,_ *Bias*). The limits of the violin represent the extreme of the fit parameters, and the boxplot within the violin marks the median (white) and interquartile range (IQR; thick black line). **d**, Same as (c) for critic learning rates (*αV(cs), αV* _*(choice)*_) and actor decay rates (*α*_*Lick*_, *α*_*choice*_). **e**, Average probability of repeating the previous choice (left or right) as a function of trial outcome from online simulations. Individual lines represent simulated sessions (***P < 0.001, paired Wilcoxon signed-ranked test with Bonferroni-Holm’s correction). Same analysis as Fig. 5e. **f**, Correlation across simulated sessions between the fraction of CS+ trials without VTA stimulations (CS+ trials without VTA stimulations/total CS+ trials) and overall performance (CS+ trials/total trials). Line represents linear regression . Same analysis as Fig. 5f. **g**, Bar plot representing the effect of removing CS+ from the model in simulation on performance (P(high reward probability choice)). Lines represent individual session (\**P* < 0.0001, Wilcoxon signed-ranked test). Same analysis as Fig. 5h. For simulations in (**e-g**) the parameters were obtained from model fits to the behavior of animals and sessions in Fig. 1-5f. **h**, Coronal sections with color saturation indicating the percentage of neurons significantly encoding *V*(*S*_*cs+*_) in each striatal domain. Grey areas were not covered by the recordings. **i**, Percentage of neurons significantly encoding *V*(*S*_*cs+*_)in each striatal subregion (*n* = 829 NAc, 1529 DS-Ros, 958 DS-Int, 720 DS-Cau). **j**, Distribution of corrected ΔR^2^ values from removing the *V*(*S*_*cs+*_) gain from the encoding model, computed by subtracting the mean ΔR^2^ of a pseudosession-based null distribution from the ΔR^2^ of the encoding model applied to the observed behavioral data (see Methods for details).*P < 0.05, **P < 0.01, ***P < 0.001, ****P < 0.0001 from χ^2^ tests. **k**, Time-resolved population decoding performance (corrected R^2^) *V*(*S*_*cs+*_) for in rewarded trials (based on groups of 40 randomly chosen neurons). Corrected R^2^ is computed by subtracting the mean R^2^ of a session-specific null distribution from the mean R^2^ of decoders applied to the observed behavioral data (see Methods for details). Lines and shading represent the mean ± s.e.m. across sessions (*n* = 9 NAc, 12 CP-Ros, 8 CP-Int, and 7 CP-Cau sessions). * indicates time points where decoding performance for CP was significantly different from NAc (*P* ≤ 0.05, from two-tailed Mann–Whitney U test at each timepoint). **l**, Same as k but for trials with >= than 5 CS+ trials w/o VTA stim. **m–q**, Same as (**h–l**) but for the choice value, *V*(*S*_*choice*_. Bar plots in (**e, g**) denote mean ± s.e.m. Bar plots in (**i, n**) denote the mean ± Bernoulli s.e.m. across all recorded neurons and the dashed lines indicate the chance level for significance (1%; see Methods for details on significance testing). Box plots represent the 10th, 25th, 50th, 75th, and 90th percentiles. For panels (**c,d,f,g**) *n* = 41 simulated sessions from *n* = 8 simulated mice. For panel **e** *n* = 24 simulated sessions from n = 7 simulated mice (sessions with >= 5 CS+ trials w/o VTA stim. For all panels see Extended Data Table 7 for statistical details and additional comparisons.

The model could be fit to the animals’ behavioral data (Fig. 6c,d, Extended Data Fig. 11b), was identifiable (Extended Data Fig. 11c,d), and could generate trial-by-trial estimates of choice and licking propensities (π^*choice*^, π^*lick*^) and of CS+ and choice state values (*V*(*S*_cs+_),*V*(*S*_*choice*_) (Fig. 6b). The choice propensities were nearly identical to the relative action values from the Q-learning model used in previous analyses (Extended Data Fig. 11e). Hence, it produced very similar performance in choice prediction compared to the Q-learning model (Extended Data Fig. 11f-i). However, licking behavior and CS+ value were not captured by the previous model (Extended Data Fig. 11e,j).

This model recapitulated key aspects of the animals’ conditioned reinforcement behavior described in Fig.5., including the similar updating of the next trial’s choice by the CS+ regardless of whether DA stimulation was delivered (Fig. 6e), the inverse relationship between good task performance and the tendency to forgo collection of DA stimulation (Fig. 6f), and the animals’ inability to do the task in the absence of the CS+ (Fig. 6g).

The most important property that allowed the model to recapitulate these features was the necessary role that CS+ value had in updating the choice propensities. This explains the dramatic deterioration in the behavior in the absence of the CS+ (Fig. 5g,h), and the lack of additional reinforcement from the CS+ with vs. without the laser (Fig. 5e, Extended Data Fig. 10).

As for the inverse relationship between performance and VTA DA stimulation collection (Fig. 5f), this arises from the model (Fig. 6f) because, in high-performing sessions, the initial value of is (*V*(*S*_*cs*+_)high and so the CS critic’s prediction error on each stimulation *(δ*_*laser*_) is small and reinforcement of the lick policy is weak relative to its passive decay. This causes licking probability to drift down and forgoing the laser collection to increase. Meanwhile, the low*α*_*cs*_ keeps (*V*(*S*_*cs*+_)high in the absence of VTA stimulation, allowing the choice Actor-Critic to operate normally. Consistent with this mechanism, we found that *across sessions* high (*V*_*0*_(*S*_*cs*+_) (initialization parameter for) significantly correlates with the probability of forgoing VTA stimulations (Extended Data Fig. 11k). Conversely, *within a session* (*V*(*S*_*cs*+_)correlated with VTA stimulation collection (Extended Data Fig. 11l), as expected based on the fact that high and (*V*(*S*_*cs*+_)high licking propensity both arise from the same error signal (VTA DA stimulation).

Finally, the model makes experimental predictions. While removing the CS+ in simulation dropped performance rapidly to chance (Fig. 6g, Extended Data Fig. 11n), removing the VTA-DA stimulation dropped performance more slowly (Extended Data Fig. 11m,n) due to the slow update of the CS+ value (*α*_*cs*+_= 0.005 ± 0.002) relative to the choice policy, which updates rapidly under a high decay-and-gain regime (*α*_*loss*_ = 0.17 ± 0.008,*β*_*win*_ = 3.98 ± 0.265; mean ± s.e.m.).

### CS+ value is encoded in the NAc

Having established that the Conditioned Reinforcer model (Fig. 6a) recapitulates the animals’ behavioral dependence on the CS+ (Fig. 5), we next examined how the latent variables in the model are encoded in the brain.

As expected based on the high correlation between the choice propensities of the new Conditioned Reinforcer model and the relative action values in the previous Q-learning model (*r*=0.99, Extended Data Fig. 11e), choice propensity was preferentially encoded in (and decoded from) the intermediate CP (*P* < 1e-13 vs. NAc, *P* < 1e-7 vs. CP-Ros, and *P* < 0.001 vs. CP-Cau by χ^2^ tests by χ^2^ tests in Extended Data Fig. 12b; *P* < 1e-13 vs. NAc, *P* < 1e-4 vs. CP-Ros, and *P* < 0.05 vs. CP-Cau by Mann-Whitney U tests in Extended Data Fig. 12c), mirroring our earlier results from relative value (Fig. 3d,f).

Next, we used the model’s trial-by-trial *V*(*S*_*cs+*_) estimates to test whether the CS+ value is represented, and if so, where this representation may be preferentially located. In striking contrast to action, relative, and state values (Figs. 2,3), which were all preferentially encoded in the *V*(*S*_*cs+*_)CP, was preferentially encoded in the NAc. NAc had more neurons encoding than *V*(*S*_*cs+*_)any other striatal subregion (Fig. 6h,i), and the magnitude of this encoding, based on the improvement in explained variance (ΔR^2^) of neural activity driven by this variable, was likewise greatest in NAc (Fig. 6j). This higher encoding in NAc was significant relative to caudal CP (*P* < 0.05 by χ^2^ test, Fig. 6i), and the ΔR^2^ of neural activity was significant relative to both intermediate and caudal CP (*P* < 1e-5 vs. CP-Int, *P* < 1e-6 vs. CP-Cau by Mann–Whitney U tests, Fig. 6j).

We examined whether population-level decoding of CS+ value corroborated this encoding result. While the encoding model accounts for the correlation between the decision variables in the behavioral model, such as state and choice value (Extended Data Fig. 11e), decoding directly from the neural data does not. To account for these correlations in our decoding analysis, we decoded each value from the residuals of an encoding model built from all other variables (see Methods). This decoding analysis confirmed the preferential localization of CS+ value to the NAc (Fig. 6k). This preferential decoding in NAc was clearest in sessions where the animal skipped at least 5 VTA DA stimulation rewards (Fig. 6l), which is what causes the CS+ value to decrease in the Conditioned Reinforcer model (while when DA stimulation is collected, it instead increases).

On the other hand, the value of the choice state (*V*(*S*_*choice*_) was not significantly encoded in any region (*P* > 0.05 vs. the 1% chance level by exact binomial test for all regions; Fig. 6m-o) and was encoded by significantly fewer striatal neurons when directly compared to (*V*(*S*_*cs+*_) < 1e-6 across all subregions by Mantel–Haenszel χ^2^ test). Similarly, decoding of(*V*(*S*_*choice*_) was weak or absent across the striatum (Fig. 6p,q).

Thus, while action values (and choice propensities) are preferentially encoded in CP, CS+ value (*V*(*S*_*cs+*_)) — the essential variable behind the conditioned reinforcement effects characterized in Fig. 5 — is preferentially represented in the NAc, the primary target of the stimulated VTA DA neurons.

## DISCUSSION

### Motivation for a new Conditioned Reinforcer model for VTA DA mediated reinforcement learning

Venerable and recent work has amply implicated DA in the NAc in conditioned reinforcement ^28,118–120^, the process through which a previously neutral cue becomes valuable and can reinforce actions. This is the basis of temporal difference reinforcement learning (RL) models of DA’s involvement in Pavlovian conditioning. Equally, dopamine in the dorsal striatum has long been associated with the process by which actions are reinforced. This is the basis of RL models of operant conditioning. However, there is a critical gap in our understanding of the link between these two phenomena, apart from general appeals to an ascending spiral architecture in the basal ganglia^121,122^.

In the most prominent RL models such as the Actor-Critic, a DA signal does double duty – training both conditioned reinforcers and actions. This means that we should expect to find the values of states in the NAc (enabling those states to act as conditioned reinforcers themselves) and that it should be possible to learn appropriate actions through the activation of DA neurons (bypassing the need for any stimuli). However, these expectations are inconsistent with two key findings in our data: i) state values (and indeed action and relative action values) from the traditional models are relatively weakly encoded in the NAc, the target site of the stimulated VTA DA neurons, when compared to the CP (Fig. 2,3), and ii) the CS+ is needed for VTA DA reinforcement to be effective (Fig. 5) ^118^.

We therefore built a new, Conditioned Reinforcer model (Fig. 6a), in which VTA DA neuron activity produces stimulus value representations in the NAc (Fig. 6h-l) (inspired by refs ^28,116,123,124^) rather than directly producing action value representations. These stimulus value representations generate a second error signal which in turn produces action value representations downstream, such as in the CP, where we observed the strongest action value correlates. This downstream influence could potentially be mediated by the ascending spiral architecture of the basal ganglia^121,122^ through a second error signal carried by SNc DA ^28,34^.

This new framework aligns with a classic view of the NAc as fundamental for Pavlovian associations^125–127^, while also clarifying how a brain structure dedicated to Pavlovian learning can indirectly impact operant behavior or action selection, such as in the present task, by endowing stimuli with a form of what has been called incentive salience^128,129^.

### Formulation of a new Conditioned Reinforcer model for VTA DA mediated reinforcement learning

We instantiated this conditioned reinforcer framework with a model composed of two interacting actor-critic modules. The first calculates the value of the CS state and controls the licking action, and the second calculates the value of the choice state and controls the decision to turn the wheel left or right. The CS module uses VTA stimulation as its error signal, while the choice module instead uses an error signal calculated by the CS critic. When fit to choices and licking behavior, this model produced a very similar choice policy to all of our earlier models (including Q-learning w/ CS+ as reward), and thus similar to our previous results (Fig. 2,3), this variable was preferentially encoded in CP. The substantial model mimicry in generating the various action values and propensities that directly determine choice arises because they are fit directly to the choices. The resolution of this mimicry and the formulation of the new model arises from the extra conditions such as removing the CS+ that allow us to interrogate the differences between the models.

One particularly salient difference lies in how they treat the value of the choice state. In the new model, it is not encoded significantly in any region (Fig. 6m-n), whereas in the model with a V-value RPE (Fig. 3a), we found it to be encoded preferentially in CP (Fig. 3e,g). The difference between the values (and hence their neural correlates in CP) stems from the fact that the learning rate of the choice critic in the new model was extremely low (alpha = 0.009 ± 0.001; mean ± s.e.m.); as indeed it also was in the traditional Actor-Critic model (alpha = 0.026 ± 0.002; mean ± s.e.m.). By contrast, in Fig. 3 we were examining state value based on the higher learning rate set by the policy of the Q-value agent (alpha = 0.245 ± 0.009; mean ± s.e.m.). This difference is consistent with what we found in our learning rate analysis of state value encoding in the striatum (Extended Data Fig. 8).

This apparently technical difference in the fitted learning rates of the two classes of model has wider implications: first that the choice critic might in fact not play its conventional role as a reinforcement comparison term in temporal difference learning (Sutton & Barto, 2018) and might not even be represented in the striatum at all; second, that we might entertain a potential modification to our formulation of the Conditioned Reinforcer model. In this, stimulus value may serve to update action values, but not the value of the choice state. In other words, the choice module might be more like Q-learning than Actor-Critic, but using the CS+ rather than DA stimulation as the RPE. This model actually fit best in our initial behavioral analysis: (Model 1;

Fig. 1). Our Conditioned Reinforcer model can be seen as an extension of that model: rather than assuming the value of the CS+ to be 1, the learning of the CS+ value from the VTA DA stimulation is explicitly modeled with the CS Actor-Critic. Thus, the fact that Q-learning w/ CS+ as reward best fit the data (and not Q-learning w/ DA as RPE or the standard Actor-Critic) provides additional support for our Conditioned Reinforcer model.

In addition, our new model allowed us to examine the CS+ value (i.e. the state value of the CS actor-critic) separately. In contrast to action values, which are preferentially encoded in CP, the CS+ value is best encoded in NAc. This implies that VTA DA stimulation directly produces stimulus values at its main target site. The encoding of this variable is sparse (∼4% of NAc neurons). This may represent a floor, because, as noted above, the CS+ value varies very little in our task design, decreasing on the (relatively rare) trials in which DA stimulation is not collected, and increasing slightly when it is collected.

### Differences between our Conditioned Reinforcer model and the traditional Actor-Critic model of the basal ganglia

Our new Conditioned Reinforcer model has several key differences from the traditional temporal difference Actor-Critic model of the basal ganglia (Extended Data Fig. 11a)^117^, which allows it to explain several behavioral and neural results that are unexplained by the traditional model.

First, in the traditional Actor-Critic model, DA is used to update the actor whether or not there is a CS, while in our model, only an error signal based on CS value (i.e. the state value of the CS) can update the choice actor-critic. Therefore, similar to the mice, our model (but not the traditional Actor-Critic) requires the CS+ to perform the task, even when VTA DA stimulation is provided immediately (Fig. 5g,h). Similarly, our model, but not the traditional Actor-Critic, explains why mice update actions similarly whether or not the VTA DA stimulation is provided (Fig. 5e).

Second, in the traditional Actor-Critic model, the critic is thought to reside in NAc and the actor in CP. Our results are consistent with the latter but not the former. Instead, when we calculate the state value of the choice state with a learning rate that matches the policy, we find state value correlates are preferentially located in CP, not NAc (Fig. 3; similar observation in ^52^). As noted above, when we fit the traditional Actor-Critic or this new Conditioned Reinforcer model to the data, and use the resulting lower learning rate for state values, we find the state value of the choice state is barely represented (Fig. 4,6). Neither finding lends support to the traditional Actor-Critic model in which state value is preferentially encoded in NAc. Instead, we find that the stimulus value (the state value specifically of the CS+) is encoded in NAc (Fig. 6h-l), consistent with our model in which VTA DA directly updates only the value of the stimulus state (and not the value of the action state). In our model, this stimulus value in NAc serves to update the choice policy in CP.

Third, the traditional Actor-Critic model is based on a single DA error signal, which trains both the actor and the critic. In order to produce conditioned reinforcement, our model instead utilizes two error signals. One error signal trains stimulus values (in NAc), which is used to produce a second error signal that trains action values (in CP). While the former error signal is provided by VTA DA, the latter could be provided by SNc DA. In fitting our model, we find a very slow learning rate for the CS value updating *αV*_*cs*_= 0.005 ± 0.002 mean ± s.e.m.), and a much faster update for the choice actor (*α*_*loss*_ = 0.17 ± 0.008,*β* ^*win*^= 3.98 ± 0.265; mean ± s.e.m.) - consistent with the relative rates of change of the quantities they are capturing^130–132^. A general question in the field is the extent DA error signals update value via synaptic plasticity as in the present models – which is thought to be a slow update ^133^ – or instead by faster changes in neural dynamics (as in working memory trained by “meta-learning”), which may instead be very fast ^109,134–138^ . The distinction in timescale that we find for the update by the two error signals suggests that the CS value updating via VTA DA could be mediated by synaptic weight changes while the action value updating may instead be mediated by neural dynamics changes rather than DA from the SNc.

### Purified experimental design enabled discovery of reinforcement learning mechanisms

Our discoveries depended on using VTA DA stimulation instead of a natural reward in our task. This is because a natural reward would have sensory properties (e.g., taste, sound, smell, sight) — in other words, a natural reward would have its own CS. Moreover, a natural reward would activate DA neurons not only in VTA but also in the SNc, which project directly to the CP and can likely reinforce actions directly^34^, without the intermediary of a CS.

In addition, the separation we discovered in stimulus vs action value coding in NAc vs CP likely benefited from head-fixation, as this eliminates contextual cues, which are likely valued in NAc ^139–142^. Given that contextual cues are correlated with actions in freely moving settings (e.g. left vs right levers are in different locations in the chamber), contextual cue values and action values would be confounded.

### Conclusions & future directions

Here, we focused on the longer-run, learning-based effects of VTA DA neuron activity. There are also shorter-term influences of VTA DA neurons over action, motivation, and forms of model-based valuation^83,143–146^, as well as affective heterogeneity within the VTA DA system^147^, which are beyond the scope of the present work. In sum, our study clarifies the power of VTA DA activity to mediate learning based on trial-and-error, along with limits to that power. Compelling prospects for future studies include dissecting the circuitry mediating the influence of VTA DA stimulation-induced conditioned reinforcers on action selection, as well as parallel experiments focusing on SNc (rather than VTA) DA stimulation.

## METHODS

### ANIMALS

11 male and 6 female adult heterozygous DAT::cre (Strain #006660, Jackson Laboratory) mice crossed with wild-type C57BL/6J (Strain #000664, Jackson Laboratory) were used in all experiments. Mice were maintained on a reversed 12 h light cycle and all experiments were conducted during the dark phase. Mice were between 3 and 8 months of age during surgery, training, and testing. Ambient temperature was maintained at 21-26 °C and humidity at 30-70% at all times. All experimental procedures were approved by the Princeton University Institutional Animal Care and Use Committee following the NIH Guide for the Care and Use of Laboratory Animals.

### STEREOTAXIC SURGERY

All surgeries were performed under isoflurane anaesthesia (5% for inductions, 1-2% for maintenance). All mice receive preoperative antibiotics (5 mg/kg, Baytril). Analgesia (10 mg/kg, Ketofen) was provided preoperatively, and postoperatively for 2 days.

Mice underwent the following surgical schedules, depending on the experiment:

A. Neuropixels recordings (Figs. 1–5; Extended Data Fig. 1-5,7-10; 6 male and 2 female): 1st surgery (Pre Training, 3 months old mice): Initial surgery where animals were headplated, viruses were injected, optical fibers and ground pins implanted, and the recording well prepared.
B. 2nd-4th Surgery (Post Training, 4-6 months old mice): Individual craniotomies for acute Neuropixels recordings. This surgery was repeated 1-2 more times to access extra recording locations.
C. Fiber photometry (Extended Data Fig. 6; 3 female): 1st surgery (Pre Training, 2 months old mice): Initial surgery where viruses were injected. 2nd Surgery (3-4 months old mice): Surgery where animals were implanted with a headplate and optical fibers implanted.
D. No Lick task (Fig. 5; 5 male and 1 female): 1st surgery (Pre Training, 3 months old mice): Surgery where animals were headplated, viruses were injected, optical fibers and ground pins implanted.

### Headplate implant

All mice had a headplate implanted as described in the IBL standardized protocol^148,149^. Briefly, the scalp and underlying periosteum was removed. Bregma and lambda were leveled, and a small steel headbar was centered at -6.9 mm anterior-posterior (A-P) relative to bregma and cemented to the skull with Metabond (Parkell).

### Virus injections

For animals used in Neuropixels recordings with optogenetic stimulation of the VTA, AAV2/5-EF1a-DIO-ChRmine-mScarlet-WPRE-hGHpA (titer: 9e12 genome copies/ml, Princeton Neuroscience Institute viral core) or AAV2/5-EF1a-DIO-hChR2-WPRE-hGHpA (titer: 2.65e12 genome copies/ml, Princeton Neuroscience Institute viral core) was injected bilaterally in the VTA (± 0.75 mm M-L, -3.1 mm A-P, -4.75 mm D-V). 500 nl were infused in each hemisphere at a speed of 100 nl/min. For the fiber photometry recordings, 500 nl of AAV2/5-EF1a-DIO-hChR2-WPRE-hGHpA (titer: 2.65e12 genome copies/ml, Princeton Neuroscience Institute viral core) was injected bilaterally in the VTA (± 0.75 mm M-L, -3.1 mm A-P, -4.75 mm D-V) and 250 nl of AAVdj-hSyn-GRAB-rDA3m-WPRE-hGHpA (titer: 5e12 genome copies/ml, Princeton Neuroscience Institute viral core) was injected in the NAc into each of 2 injection sites (1st: ± 0.75 mm M-L, -3.1 mm A-P, -4.75 mm D-V, 2nd: - 0.75 mm M-L, -3.1 mm A-P, -4.25 mm D-V) and CP (1st: - 2.5 mm M-L, 0 mm A-P, -3 mm D-V, 2nd: -2.5 mm M-L, 0 mm A-P, -2.5 mm D-V). All viruses were infused at a speed of 100 nl/min (Nanofil syringes with a UMP3 ultramicro pump, WPI). At the end of the injection, we waited a minimum of 5 minutes before retracting the syringe.

### Optical fiber implants

Optical implants used for optogenetic stimulation were made in house. The fiber (Ø300 mm core, 0.39 NA, 2.5 mm ferrule, Thorlabs) was polished and tested for an even transmission profile and a minimum transmission efficiency of 80%. For optogenetic stimulation of the VTA, optic fibers were implanted bilaterally at the following coordinates: ± 0.4 mm medial-lateral (M-L), -3.1 mm anterior-posterior (A-P), -4.15 mm dorsal-ventral (D-V). These locations were reached with a 10° M-L/D-V rotation (rotated coordinates: -3.1 mm A-P, ± 1.1 mm M-L, 4 mm D-V).

For fiber photometry recordings, low-autofluorescence optical fibers encased in a ferrule (Ø200 mm core, 0.37 NA, 1.25 mm ferrule, Neurophotometrics) were implanted unilaterally at each of the following locations (fiber tip location relative to bregma): DLS (-2.6 mm M-L, 0 mm A-P, -2.8 mm D-V) and NAc (-1 mm M-L, 1.45 mm A-P, -4.5 mm D-V).

### Recording well preparation

For animals used for Neuropixels recordings, at the time of headplate implantation, a well was built with cement around the perimeter of the dorsal skull and covered with UV transparent glue (Norland Optical Adhesives #81, Norland) and a small ground pin (82K7797, Newark electronics) was inserted on the right hemisphere. This UV glue protected the exposed skull and was removed on the first craniotomy surgery for recordings.

## BEHAVIOR

### Behavioral apparatus

We utilized the International Brain Laboratory’s standard behavioral setup, with comprehensive information on its parts and operation available in their documentation ^149,150^. Detailed component lists and setup instructions for the training and electrophysiology rigs can be found in ^151^ and ^152^, respectively.

The behavioral setups included “training” rigs that were not set up for electrophysiology, as well as an “electrophysiology” rig that was set up for electrophysiology. The training rigs were constructed with Thorlabs pieces inside a small soundproof cabinet (9U acoustic wall cabinet 600 3 600, Orion). The rigs featured an LCD screen (LP097Q, LG), a custom 3D-printed mouse holder to headfix the mouse in front of the screen, and a steering wheel (86652 and 32019, LEGO) positioned beneath its forepaws. A spout, connected to a water reservoir and controlled by a solenoid valve (225P011-21, NResearch), delivered water without obstructing the mouse’s view. The rigs also included a speaker (HPD-40N16PET00-32,Peerless by Tymphany) above the screen for generating sounds. A rotary encoder (05.2400.1122.1024, Kluber) tracked wheel movements. Visual stimulus timing was measured using a photodiode (Bpod Frame2TTL, Sanworks), in order to detect screen pixel flips. All task-related devices were controlled by a Sanworks Bpod State Machine, with task logic programmed in Python and visual stimulus/video handled by Bonsai and BonVision ^153^. The electrophysiology rig was a modified version of the training rig, situated on an air table (VH3048W-OPT Newport) within a larger custom soundproof cabinet to accommodate electrophysiology equipment. It differed by using a USB camera (CM3-U3-13Y3M-CS, Point Grey) for lick detection, a custom speaker (Champalimaud Foundation, V1.1), and 4x micromanipulators sitting in platforms above the mouse holder (uMp-4, Sensapex) outside the mouse and camera’s field of view. In the electrophysiology rig, behavior, video, audio and neural (electrophysiology or phometry) signals were synchronized through TTLs sent to a PXI system (National Instruments) ^150^.

All electrophysiological and photometry recordings, as well as pharmacological inactivation experiments, were conducted in the electrophysiology rig. Both rig types were used for training in the probabilistic reversal learning task.

### Probabilistic reversal learning task

Mice were trained to perform a probabilistic reversal learning task for VTA stimulation, which required them to track their history of rewards and choices to select the choice with higher reward probability (Fig. 1-4,5a-f). To initiate each trial, animals were required to hold the wheel still for a quiescent period that varied between 0.2 to 0.5 s. After this quiescent period, two identical Gabor patches (100% contrast, 0.1 cycles/degrees, vertical orientation, random phase) were presented on the left and right sides of a screen. Choices were made by rotating the wheel. Wheel movement was coupled in closed-loop fashion to the horizontal position of the Gabor patches with a gain of 4 visual degrees per mm of wheel displacement (2.17 visual degrees per degree of wheel rotation). A choice was registered when the mouse centered one of the two Gabor patches on the screen (35°). Importantly, the patches gave no information about reward probabilities. Clockwise and counterclockwise movements were classified as left and right choices respectively. When a choice was registered, the animal was either rewarded or unrewarded and the selected Gabor (left or right) froze in the center for 4 s regardless of the choice’s outcome. If a choice earned a potential VTA stimulation, a 100 ms duration, ∼ 65 dB narrowband sound (12-13 kHz) indicated VTA stimulation availability, and served as a positive conditioned stimulus (CS+).Upon the onset of this tone, VTA stimulation was contingent on the animal’s licking of a dry spout, with licking detected using a video camera ^154^. If the mouse licked the spout at any point during the 1 s period following this sound, stimulation was delivered. The VTA stimulation consisted of a 1 s train of laser stimulation (ChR2: 447 nm, ∼8 mW, 5 ms pulse width, 20 Hz; ChRmine: 532 nm, ∼0.5 mW, 5 ms pulse width, 20 Hz) generated by a DPSS laser (Shanghai Laser and Optics & Co) controlled with a Pulse Pal (Sanworks). Rewards were not baited. Unrewarded trials were signaled by 0.5 s of ∼ 65 dB white noise, which served as a negative conditioned stimulus (CS –). A 4-5 s inter-trial interval (ITI), following the CS+/CS– onset, followed each trial regardless of its outcome. If no choice was completed within 60 s from the go cue, the CS– was played and a new trial started after the ITI. Whether a choice was rewarded was dependent on a probabilistic block structure. Within each block, one choice (left or right) was designated as the high-reward probability choice (70% reward probability), while the other choice was designated as the low-reward probability choice (10% reward probability). Block lengths had a default random length between 20 and 40 trials, drawn from a uniform distribution. At each block transition, the high- and low-reward probability sides were switched. To prevent side biases, a debiasing mechanism was implemented such that blocks did not advance until the mouse made at least 7 choices toward the high-reward probability side within the preceding 10. This debiasing procedure could therefore extend block length beyond its default 40 trial upper limit.

All recordings were performed with mice that were already fully trained to perform the task. Sessions lasted between 60-90 minutes (546.5 ± 16.3 trials per session, mean ± s.e.m.). Sessions were only included for analysis when mice completed at least 9 blocks and showed win-staying behavior, that is when animals were more likely to repeat a choice when rewarded (after a CS+). For the “No CS+” condition in Fig. 5g-h, in which animals rarely progressed past the first block due to the lack of a CS+ we did not apply this performance criterion since that would have excluded the entire condition by design, rather than filtering for data quality. All sessions in the corresponding control condition met the standard criterion.

### Training in the probabilistic learning task

To train on the task, mice underwent a shaping procedure consisting of 4 stages. During shaping, the animals received 4ul of 10% sucrose solution as a reward, instead of optogenetic stimulation. While the same CS– as in the final task signaled unrewarded trials, the sucrose reward itself was accompanied by the sound of the solenoid valve but importantly was not paired with the 12-13 kHz narrowband CS+ used in the final task. In Stage 1, the high- and low-probability choices had a 100% and 0% probability of reward, respectively. Block lengths in this stage were 10 to 30 trials. Unrewarded choices had a 10s ITI. To facilitate the association of turning the wheel with the selection of the left or right Gabor patch, in this stage, the two patches had different spatial frequencies (0.1 cycles/degrees for the left Gabor and 0.5 cycles/degrees for the right Gabor). Once mice reached 70% correct choices in Stage 1 (14.3 ± 3 sessions on average), they advanced to Stage 2 . Stage 2 was identical to Stage 1, but the left and right Gabor patches had the same spatial frequency of 0.1 cycles/degrees (as in the final task). Upon reaching 70% correct choices in Stage 2 (2.83 ± 0.542 sessions on average), mice progressed to Stage 3, where the high- and low-reward probability choices had a reward probability of 70% and 10%, respectively, as in the final task. Equally, ITI for both rewarded and unrewarded trials were 4-5 s, as in the final task. Once mice could complete at least 9 blocks and showed win-staying behavior (17.9 ± 4.6 sessions on average), they moved to Stage 4 (as long as a minimum of 4 weeks from viral injection had passed to ensure ChR2/ChRmine expression). Stage 4 consisted of one to two 30-min conditioning sessions to learn to lick for the VTA DA stimulation reward. During these sessions, mice received a 1 s train of VTA DA optogenetic stimulation (1 s train, 5 ms pulse width, 20 Hz) in response to licking a dry spout within a 1s window following the sound of the CS+ from the final task (an unheard sound up to this point in training). After this conditioning stage, mice were moved to the final task (probabilistic learning task).

### Water restriction

Water access was restricted from 1 week prior to the start of training until the end of the experiment. Initial water restriction was necessary as mice worked for water rewards during early shaping. Later, when reward identity changed to optogenetic DA stimulation, mice were kept on water restriction to maintain a comparable internal state between testing and training. Weights never dropped more than 20% from their pre-water restriction weight, and mice consumed a daily minimum of 1 ml of water per 25 g of body weight. During the training phases of the probabilistic learning task (where mice received water rewards) most mice were able to obtain the majority of their daily allocation of water through the task alone. When this minimum was not achieved during the task, mice were supplemented at the end of the day. Once the mice moved to optogenetic rewards, they were provided with water at the end of every day.

### No Lick Task

In order to understand how the presence of a CS+ affected choice behavior, we performed an experiment (Fig. 5g-h) where mice worked for VTA DA optogenetic stimulation paired with an auditory CS+ (“No Lick Task with CS+”) or without it (“No Lick Task without CS+”). In our standard probabilistic reversal learning task, the CS+ indicated the presence of a potential optogenetic reward that could be obtained by licking an empty spout during the 1s following the CS+. This introduced a potential confound: Without removing this lick requirement any change in task performance in the no CS+ task could be due to a reduction in the collection of optogenetic stimulation rewards and not the absence of the CS+, since the animal might not know when to lick after making a rewarding choice. To prevent this, in this experiment optogenetic stimulation delivery was automatically provided after making a rewarding choice, removing the lick requirement. Additionally, the co-occurance of the CS+ and the VTA DA stimulation in this task variant ensured that the importance of the CS+ to performance cannot be attributed to a shorter or stronger temporal relationship between the action and the CS+ (compared to the action and VTA DA stimulation). Animals used in this task were first trained in the “No Lick Task with CS+” following the shaping steps described for the standard probabilistic reversal learning task. Once trained, mice first experienced the” No Lick Task with CS+” followed by the “No Lick Task without CS+” in the following session.

### BEHAVIORAL MODELS

To understand the decision-making processes of mice in our task, we built and tested three families of models: 1) Q-learning, 2) policy gradient (REINFORCE), and its extension 3) the Actor-Critic (Fig. 1.; Extended Data Fig. 2.; Extended Data Table 1). All models were designed to predict trial-by-trial choices based on slightly different assumptions about how mice learn from reward and experience. All models were designed to predict trial-by-trial choices based on different assumptions about how mice learn from reward and experience. Parameters were fit in a hierarchical Bayesian manner, allowing variability to be modelled at three distinct levels: the population, individual subjects, and experimental sessions.

We start by detailing the model families; we then describe the fitting procedure, describe the comparison and quality controls we used and finally how we derived point estimates of the decision variables that we then used for neural analyses.

### Models

#### Models 1-2: Extended Q-learning models

We tested two versions of Q-learning models ^155,156^ (Extended Data Table 1), one where the CS+ was treated as a reward regardless of whether the VTA DA stimulation reward was collected (Model 1), and a version where VTA DA stimulation was treated as an RPE (Model 2). Both models were extended to incorporate mechanisms to account for forgetting (an exponential decay of unchosen option values) and perseveration (the tendency to repeat previous choices irrespective of outcome) which are known factors to affect decisions ^157^.

#### Model 1: Extended Q-learning model with the CS+ as reward

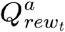 for each choice on trial *t* (*a*∈{0 if the choice was left, 1 if the choice was right }) were initialized at 0 at the start of each session. In every trial, *t*, the 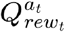 value for the chosen option (*a*_*t*_) was updated according to the following equations based on the outcome *o*_*t*_, where *o*_*t*_ =1 for a reward, and *o*_*t*_ =0 for no reward:

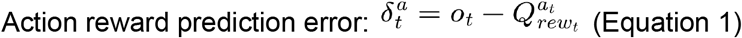

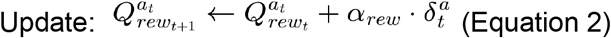

Here, *α*_*rew*_ was a free learning rate parameter. Trials where a CS+ was delivered, were treated as *rewarded* independently of whether the VTA stimulation was later consumed. *Unrewarded* trials only included the trials where a CS– was delivered.

To account for the possibility that the value of unchosen options might decay over time, we introduced a forgetting mechanism into the model ^36^. For the unchosen option, *ã*_*t*_=1− *a*_*t*_, the Q-value was updated according to:

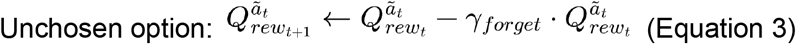

where *ϒ*_*forget*_ can be seen as a learning rate for the unchosen option’s value.

To capture the observed tendency of subjects to repeat choices regardless of their outcome, we incorporated a perseveration term ^157–160^. This was done by introducing 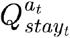, which represented a weighted average of the mouse’s past choices. This ‘stay’*Q*− value was updated as follows:

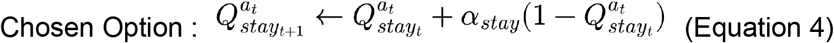

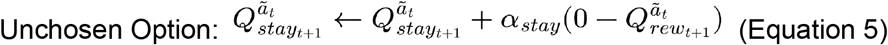

Here *α*_*stey*_ is a free learning rate specific to .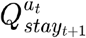.

Finally, the probability of choosing a given option (*a*_*t*_+1) on trial *t*+1, was then determined by passing the reward-based Q-values 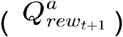 and the perseveration Q-values 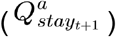 through a logistic sigmoid function *σ* (.):

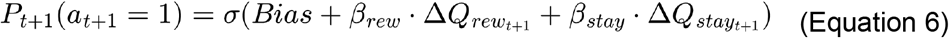

Where 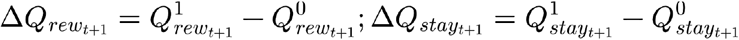 are the relative preferences for choosing R over L. Here *β*_*rew*_ and *β*_*stay*_ are free parameters acting as weights for the value and perseveration components of the decision, respectively. *Bias* is a free parameter accounting for the subject’s innate choice bias in favor of a right choice.

#### Model 2: Extended Q-learning model with VTA DA stimulation as an RPE

Given that DA is often thought to serve as a reward prediction error (RPE), not a reward, we also considered a version of Q-learning where a) the DA stimulation took the role of an RPE and b) trials where a CS+ was delivered but the VTA DA stimulation was not collected were treated as unrewarded ^83^. Therefore in this model we substituted the 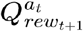 update rule for the chosen rewarded action described in Equation 2 with:

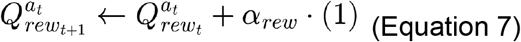

The unrewarded and unchosen options continued to be updated by the rules in Equation 2 (with 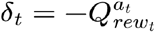 and Equation 3 respectively. The perseveration component was modelled as in Model 1 (Equation 4-5) and choice probabilities were calculated as described in Equation 6.

#### Model 3: Q-learning with a Dynamic (Surprise-Modulated) Learning Rate

This model extends Model 1 (Extended Q-learning model with the CS+ as reward), retaining Model 1’s forgetting and perseveration mechanisms and choice rule, but replacing the fixed learning rate *α*_*rew*_ governing the chosen option’s reward-based Q-value update with a learning rate that varies from trial to trial, scaling with the recent magnitude of prediction error. This is akin to Pearce–Hall associability ^84,85^, or surprise-modulated learning-rate models.

As in Model 1, trials in which a CS+ was delivered were treated as rewarded (*o*_*t*_ =1) independently of whether the VTA stimulation was subsequently collected, and CS– trials were treated as unrewarded (*o*_*t*_ = 0). For the chosen option (*a*_t_), an action reward prediction error was computed as:

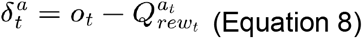

The chosen option’s reward Q value, 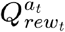, was then updated using a learning rate equal to the product of a fixed, freely-estimated base rate, *α*_*rew*_, and a dynamic scalar, *k*_*t*_, tracking the recent unsigned prediction error:

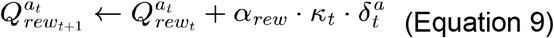

*k*_*t*_ was itself updated after every trial as an exponentially-weighted running average of the absolute prediction error:

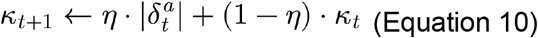

Where *η* is a free parameter, bounded between [0,1], governing how quickly *k*_*t*_ tracks recent surprise: values near 1 make the effective learning rate highly sensitive to the most recent trial’s prediction error, while values near 0 make *k*_*t*_ effectively constant across the session.*k*_*t*_ was initialized at a fixed value of 0.5 at the start of every session (not a free parameter).

The unchosen option’s Q-value continued to decay according to Model 1’s forgetting mechanism (Equation 3), and the perseveration (“stay”) Q-values were updated identically to Model 1 (Equations 4–5), with a learning rate *α* _*stay*_ fit separately for this model. Choice probability was computed exactly as in Model 1 (Equation 6), using free parameters *β*_*rew*_, *β*_*stay*_ and *Bias*, with the dynamically-scaled 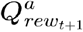 entering 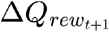 in place of Model 1’s fixed-learning-rate value

#### Model 4: REINFORCE

We tested a version of a policy gradient-based model, ‘REINFORCE’ ^161^, which directly updates the decision variable (or action propensity) based on the outcome of a trial, rather than computing Q-values for actions. This model had no perseveration component since it was non-beneficial in early testing.

For each trial *t*, the propensity for choosing R over L (*π* _*t*_) was updated based on the trial’s outcome:

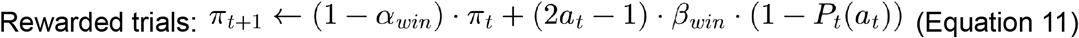

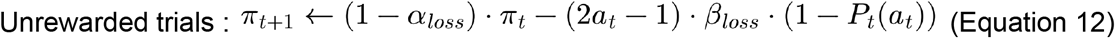

Here, 2_*at*_ − 1 represents the action taken in trial *t* (+1 for Right, -1 for Left). The parameters *α*_*win*_ and *α*_*loss*_ are free forgetting rates specific to rewarded and unrewarded trials, respectively. Similarly, *β*_*win*_ and *β*_*loss*_ are free parameters that modulate the influence of the action prediction error(1-*P*_*t*_(*a*_*t*_)) on the update for rewarded (CS+ delivered) and unrewarded outcomes (CS– delivered).

The probability of choosing a specific option *P*_*t*+1_(*a*_*t*+1_) was determined by passing *π* _*t*+1_through a sigmoid function:

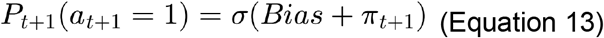

Where *Bias* is a free parameter accounting for any intrinsic preference for a right choice. *π* _*t*+1_ effectively accumulates evidence for right over left choices, with its magnitude and sign directly influencing the choice probability.

#### Model 5: Actor-Critic

We also implemented an Actor-Critic model extending from the REINFORCE model^82^. This architecture combines a ‘Critic’ component, which learns to predict the value of states, with an ‘Actor’ component, which uses these value predictions to update its policy (i.e., its decision-making strategy). This framework allows for learning through both reward prediction and direct policy adjustments based on observed outcomes.

The Critic component was responsible for learning the expected value of the current trial, denoted as *V*_*t*_. In each trial *t*, the Critic first computed astate reward prediction error 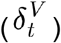, which quantifies the difference between the actual reward received (modeled using the CS+ as a fixed-size reward) and the value (*V*_*t*_) of the trial:

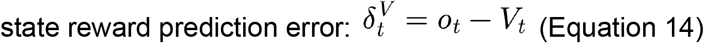

based on the outcome .*o*_*t* ._

The trial value *V*_*t*_ was then updated based on this prediction error and a learning rate, *α*_*rew*_ :

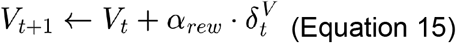

At the start of each session*V*_*t*_ was set to 0.

The Actor component was responsible for learning the propensity *π*_*t*_ that directly determined the probability of choosing an action, similar to the REINFORCE model. Critically, its updates were modulated by the *δ*_*t*_from the Critic, which acted as an adaptive baseline.

The value of *π*_*t*_ was updated depending on whether the trial was rewarded or unrewarded Rewarded trials:

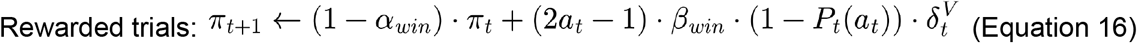

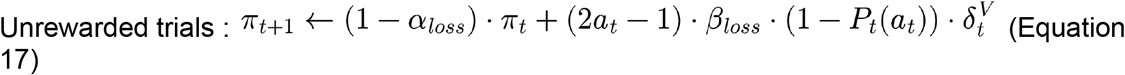

As for REINFORCE,2*a*_*t*_ − 1 signifies the action taken (+1 for Right, -1 for Left). The parameters *α*_*win*_ and *α*_*loss*_ are learning rates for the *π* update specific to rewarded and unrewarded outcomes, respectively. Similarly *β*_*win*_, and *β*_*loss*_ are free parameters modulating the influence of the prediction error (1 − *P*_*t*_(*a*_*t*_)) and the Critic’s 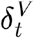 on the Actor’s update. The critical distinction here is the multiplication by 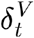, which scales the learning based on the magnitude and sign of the reward prediction error.

The probability of choosing a specific option (*P*_*t*_+1(*a*_*t*_+1)) was then calculated using a sigmoid function, incorporating *π*_*t*+1_ and a *Bias* term as previously described in Equation 13.

#### Model 6: Conditioned Reinforcer

Inspired by our neural and behavioral results (Fig. 2-5), we built a Conditioned Reinforcer model (schematized in Fig. 6a), extending the Actor-Critic architecture described in Model 5. Here, two interconnected Actor-Critic modules operate in sequence within a trial. The first calculates the value of the CS state and controls the licking decision, and the second calculates the value of the choice state and the decision to turn the wheel left or right. The CS module uses VTA stimulation as its error signal, while the choice module instead uses an error signal calculated by the CS critic.

Each trial is modelled as a sequence of states:

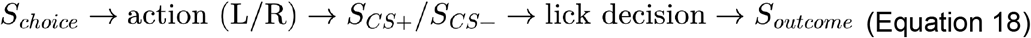

#### CS Actor-Critic

The CS Actor-Critic module learns the value of the CS+ and controls licking behavior (i.e. the collection of VTA Stimulation). The Critic tracks how much the CS+ is worth; the Actor uses that signal to set how readily the mouse licks.

On CS+ trials, the CS Critic tracks the ongoing value of the CS+,*V*_*t*_(*S*_*cs*+_), via a prediction error,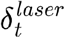, computed against the laser outcome (VTA DA stimulation):

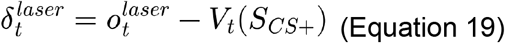

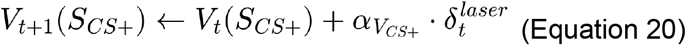

Where 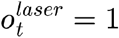 if the mouse licked (and thus triggered VTA stimulation) and 0 otherwise. This update is applied on every CS+ trial regardless of whether the mouse licked. Although VTA stimulation delivers a fixed dopaminergic signal, computing it as a prediction error (rather than treating it as a fixed reward) prevents *V*(*S*_*cs*_+) from growing without bound and allowing it to stabilize over time; without this, well-trained mice would never be predicted to forgo the reward, contrary to what we observe. This is consistent with recent work by Carter et al. ^83^.

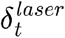 in turn updates the CS Actor, which governs the lick decision through its own propensity variable, 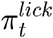 without any additional bias (see below):

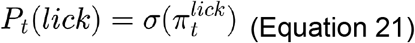

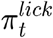 is updated according to the prediction error when the mouse licks; if the mouse does not lick (forgoing the available reward), it simply decays toward its unconditioned baseline:

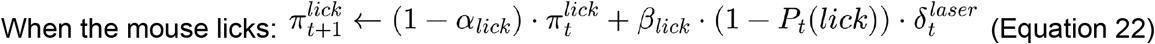

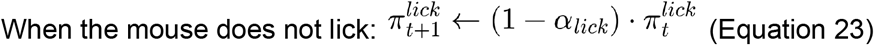

On CS-trials, the model assumes the CS-has no value (*V*(*S*_*cs*−_) = 0), since it has never been paired with reward and has no capacity to reinforce or trigger licking. The CS Actor-Critic module is therefore entirely inactive: there is no lick decision to make (the laser is not available), so 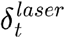 is not computed and neither *V*(*S*_*cs+*_) nor 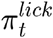 is updated. Both simply carry over unchanged from the previous trial. Note that 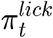 does not decay here, as Equation 23 applies only to a no-lick CS+ trial.

Beyond controlling licking, *V*(*S*_*cs*_) (*V*(*S*_*cs+*_) or *V*(*S*_*cs*−_)) feeds forward into the Choice Actor-Critic, which evaluates the choice state and controls the left/right wheel turn.

#### Choice Actor-Critic

At CS onset, the Choice Critic computes a prediction error, 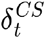, reflecting the change in value between the choice state *S*_*choice*_ to the CS state 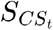.

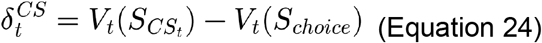

*V*_*t*_(*S*_*choice*_) is a slowly-moving baseline reflecting the value of engaging in the task, and 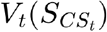 reflects the value of the CS state on trial *t*, with 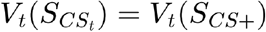 on CS+ trials, and *V*_*t*_(*S*_*cs*_) = *V*_*t*_(*S*_*cs*−_)on CS-trials (as above). Consequently, on CS+ trials 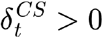 whenever *V*_*t*_(*S*_*choice*_) is below the current CS+ value, reflecting conditioned reinforcement, while on CS– trials 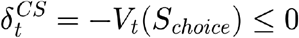.

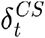 is used to update *V*_*t*_(*S*_*choice*_):

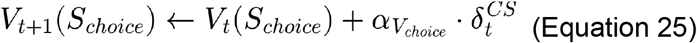

where *αV*_*choice*_ is a learning rate for *S*_*choice*_. The Choice Actor maintains a single propensity variable, 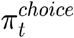, encoding the relative preference for choosing right over left, initialized at 0 at the start of each session and updated as follows:

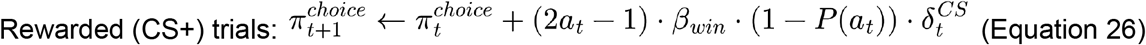

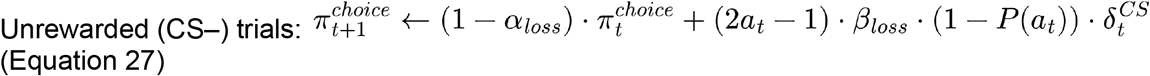

As in the previous Actor-Critic (Model 5),2_*at*_ -1 denotes the action taken (+1 Right, -1 Left), *α*_*loss*_ in an outcome-specific decay rates, and *β*_*win*_/*β*_*loss*_ scale the influence of the action prediction error (1 − *P*_*t*_(*a*_*t*_)) and 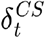 on the update.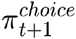 then determines choice probability through the sigmoid function together with a term, *Bias* as in Equation 13.

#### Initialization

*V*_*0*_(*S*_*cs*_+), *V*_*0*_(*S*_*choice*_), and 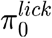 are not initialized at zero. We assume they enter each session with pre-formed estimates, which may vary from session to session. We therefore fit session-level initial values *V*_*0*_(*S*_*cs*_+), *V*_*0*_(*S*_*choice*_), and 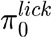 for the start of each session.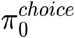, by contrast, is initialized at 0 at the start of every session, and choice probability includes no separate lick-specific bias term (any baseline lick tendency is effectively captured by 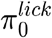.

### Fitting procedure

Each model was fitted using Hamiltonian Markov chain Monte Carlo (MCMC) sampling in Stan (2.36.0) ^162^. We employed the standard methodology of generating latent parameter values from Gaussian distributions, and then using non-linear transformations to map them into the appropriate spaces for each model (e.g., using log-odds transform for parameters confined to the interval [0,1], Extended Data Table 2).

We fitted the model hierarchically which allowed us to model variability simultaneously at three distinct levels: the population, individual subjects, and experimental sessions. By sharing information across these levels, we aimed to obtain more robust and precise estimates of the underlying parameters. The general hierarchical structure of the modelled parameters was as follows:

#### Population Level

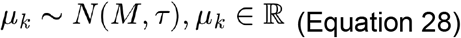

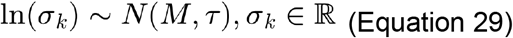

Where *μ* and *σ* represent the means and standard deviations for the *k* hyperparameters governing the distributions at the individual level.*M* is the mean and *τ* the standard deviation for the prior (see Extended Data Table 2 for the full list of priors for each parameter).

#### Individual Level

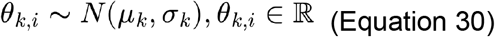

Here, *θ* represents the individual-level means for subject i (where *i* = 1, …,*n*, and *n* is the number of subjects) for each parameter *k*. These individual-level parameters were drawn from distributions parameterized by the population-level hyperparameters *μ* and *σ*.

#### Session Level

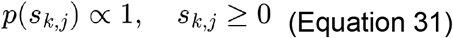

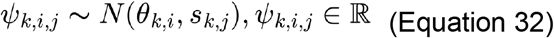

At the session level, *ψ* represents the session-specific parameter *k*, for subject *i* during session *j* (where *j* = 1,…*z*) where *z* represents the number of sessions for subject *i*. These session-level parameters were drawn from distributions parameterized by the individual-level means *θ*_*k,i*_ and variance *s*_*k,j*,_ *s*_*k,j*_. was sampled from an uniform positive prior (Equation 33).

Model 6 used the same three-level structure (Equations 29–33), with two departures. First, rather than sampling *θ*_*k,i*_ and *ψ*_*k,i,j*_ directly from *N (μ*_*k*_, *σ*_*k*_)and *N* (*θ*_*k,i*_,*s*_*k,j*_) we used a non-centered parameterization at both levels: standardized deviates *z*_*k,i*,_ *z*_*k,i,j*_ ∼ *N* (0,1) were drawn and the parameters reconstructed as *θ*_*k,i=*_ *μ*_*k*_ + *σ* _*k*_ · *z*_*k,i*_ and *ψ*_*k,i,j*_ = *θ*_*k,i*_+ *s*_*k,j*._ · _*zk,i,j*_This is mathematically equivalent to Equations 31 and 33 but decouples location and scale in the sampler’s geometry, which avoids the funnel-shaped posteriors that centered parameterizations are prone to under hierarchical models. Second, not every parameter in Model 6 was given session-level variability: *αV*_*cs*+_, *αV*_*choice*_, *α*_*lick*_, and β_*lick*_ were estimated at the population and individual levels only (i.e., *ψ*_k,i,j_= θ _*k,i*_ for these parameters, with no session term), as there was insufficient information within individual sessions to identify them accurately at that level. All remaining parameters, including *V*_*0*_(*S*_*cs*_+) and *V*_*0*_(*S*_*choice*_), which were built on the logit scale via the same non-centered hierarchy and mapped to [0,1] using the inverse-logit transform, followed the full three-level structure described above.

#### Prior Distributions

Finally, we employed weakly informative prior distributions for all parameters (see Extended Data Table 2 for details) to allow the data to largely drive the posterior inference. At the population and individual level we ensured positivity of all standard deviations by applying a exp(.) transformation during sampling (see Extended Data Table 2 for details).

#### Inference

For each model, we ran four independent chains, each with 8000 iterations, including a warm-up period of 2000 iterations and a parameter adapt_delta set to 0.99. Posterior inference was based on the samples obtained after the warm-up period. Samples from the posterior distribution were extracted using the Python package PyStan.

### Quality controls

#### Posterior analysis

Convergence of the MCMC chains was assessed by ensuring that the 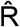 statistic for all parameters was below 1.01. We also assessed the shape of the posterior with histograms to inspect the shape, central tendency and variance of the parameter estimates^163^.

#### Recovery analysis

We performed recovery analysis to ensure that the fitted parameters were identifiable ^164^, that is to test whether the behavior can be best explained by a single set of parameters. To do this we generated a set of simulated data with the fitted parameters as the ground-truth and then fitted the data to these models. By comparing whether the fitted parameters closely followed the ground truth parameters we determined whether the parameter fits were identifiable (Extended Data Fig. 2d, Extended Data Fig. 11c,d).

#### Confusion analysis

While the recovery analysis detects whether the parameters within a model are meaningful, the confusion analysis tested whether different models are distinguishable ^164^. In other words it diagnoses whether the models are separable under our task design and parameter ranges. It is not a comparison of model fit to the observed data. To do this we generated simulated data from the best model of each family (Q-learning with the CS+ modeled as a reward, REINFORCE and the Actor-Critic). We then fit each model to each simulated datasets. Finally, we tested whether the ground-truth model was significantly better at capturing the behavior than the other models (Extended Data Fig. 2b,c).

### Model comparison

To compare the predictive performance of the candidate models (Models 1–6), we used Pareto-smoothed importance sampling leave-one-out cross-validation (PSIS-LOO; ^165^), implemented in the Python package ArviZ (Extended Data Fig. 2a, Extended Data Fig. 11f).

Models were compared on 22,406 trials from 41 sessions in 8 mice. For each model, we supplied the full posterior (all chains and draws, excluding sampler diagnostic columns) together with the per-trial log-likelihood, so that the relative effective sample size was estimated from the posterior draws rather than assuming independent MCMC samples.

For each model we obtained the expected log pointwise predictive density (*ELPD*_*loo*_) and its standard error, the effective number of parameters (*p*^*loo*^), and the Pareto- *k* diagnostic for each trial, which flags observations for which the importance-sampling approximation may be unreliable. With 24,000 post-warmup draws the diagnostic threshold is 0.7 (Vehtari et al., 2024); all values fell below this except five of 22,406 observations (0.02%) in Model 5 (Actor-Critic). Models were then ranked by *ELPD*_*loo*_, and pairwise differences in predictive performance (*ΔELPD*), along with the standard error of each difference (dSE), were computed using ArviZ’s compare function. Standard errors are computed across trials. Trial ordering was verified across models. For interpretability we also report per-trial predictive probability, exp(*ELPD*_*loo*_*/N*), the geometric mean probability assigned to the observed choice (chance = 0.5) in the figure caption.

Because Model 6 (Conditioned Reinforcer) outputs separate log-likelihoods for the choice (log_lik_c) and lick (log_lik_l) decisions, whereas Models 1–5 predict choice only, model comparison across all six models was restricted to log_lik_c for Model 6, so that all models were evaluated on their ability to predict the animals’ choices.

### Decision Variable Estimation for Neural Analysis

Finally, to correlate with neural activity, we estimated decision variables from two models: 1) In Fig. 2,3, our best-performing model overall, as determined by model comparison (Model 1: Extended Q-learning model with the CS+ as reward, Fig. 1c,d and Extended Data Table 1), and In Fig. 6, Model 6 (Conditioned Reinforcer), which captures the conditioned reinforcement effect described in Fig. 5. These estimates were derived by running each fitted model “off-policy.” This involved feeding the mice’s actual choices, outcomes, and (for Model 5) licks into the model to update its decision variables. For each set of post-warm-up parameter draws (6000) from the posterior, we calculated the value of each decision variable on every trial. This process generated a posterior distribution for each decision variable at each trial, and our final point estimate for neural analyses was the median of this posterior distribution.

We extracted four primary decision variables from the Q-learning model: Contralateral (*Q*_*Contra*_) and Ipsilateral (*Q*_*Ipsi*_) Action Values, Relative Value (*ΔQ*), and State Value (*V*). From the Conditioned Reinforcer model, we extracted three additional variables that capture the conditioned-reinforcement and task-engagement signals unique to that architecture, and that have no equivalent in the Q-learning model: the choice propensity (*π*^*Choice*^), the choice-state value (*V*(*S*_*Choice*_), and the learned CS+ value (*V*(*S*_*cs+*_)).

*Action values (Figs. 2,3,4; Extended Data Figs. 4, 5, 8):*

Contra- and Ipsilateral values were calculated using the following equation:

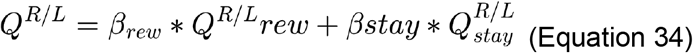

Where each value component (*Q*^*R/L*^*rew* and *Q*^*R/L*^ *stay*) were calculated using Equations 1-5. R or L was labelled Contralateral or Ipsilateral depending on the recording location (e.g. *Q*^*R*^ was *Q*^*Contro*^ for recording in the left hemisphere).

*Relative and State Value (Fig. 3; Extended Data Fig. 7):*

The Relative Value (Δ*Q*) represents the model’s subjective preference for one choice over the other, incorporating both reward-based value and perseverative tendencies. It was calculated as:

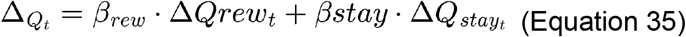

The components Δ *Qrew*_*t*_ (difference in reward-based Q-values) and Δ *Qstay*_*t*_ (difference in perseverative Q-values) were updated on every trial according to the rules described in the Q-learning model section (Equations 1-5).

The State Value (*V*) represents the model’s expectation of future reward from the current trial. This variable reflects the overall reward expectation in trial-by-trial bases. It was calculated as:

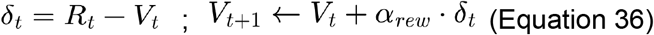

Where*R*_*t*_ indicates if a reward was given a trial *t* and *Vt* is initialized at 0.

We also analyzed a variant of state value that was parametrized as the sum or weighted sum of action values (Extended Data Fig. 7). They were calculated as:

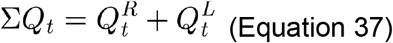

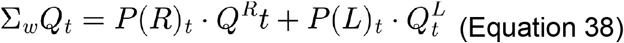

Here 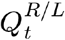 is calculated as described in Equation 34 and *P(R/L)*as in Equation 6.

*Conditioned Reinforcer Decision Variables (Fig. 6; Extended Data Fig. 11, 12):*

*π*^*choice*^ reflects the model’s evolving preference for right over left, incorporating both the animal’s chosen action and the conditioned-reinforcement signal, *δ*_CS_, delivered by the CS+. It was updated on every trial according to Equations 22–23. Unlike the Contra/Ipsi action values *π*^*choice*^ above, is a single relative-preference term, since the Choice Actor in this model maintains one propensity variable rather than separate per-action values.

*V(S*_*choice*_) reflects a slow-moving estimate of the value of the choice state, independent of the action ultimately taken, and was updated on every trial according to Equation 20.

*V* (*S*_*CS*_+) reflects the model’s evolving estimate of the value of the CS+ itself. It was updated only on CS+ trials, according to Equations 24–25, and held at its previous value on CS– trials, when the CS Actor-Critic module is inactive.

### Comparing action value updating by the CS+, with and without the VTA DA stimulation (Extended Data Fig. 10)

To compare action value updating by the CS+ with vs without VTA DA stimulation, we fitted a new version of Model 1 (Equations 1-6). In this version, every update rule was maintained from the original model (Equation 1), except we updated the Q-value of the chosen option differently after a CS+ trial, depending on whether or not the VTA DA stimulation was collected :

- CS+ with VTA DA stimulation

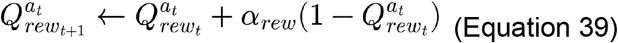
- CS+ without VTA DA stimulation

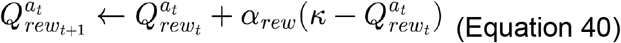

Here was a fitted free parameter representing the value of the CS+ in the absence of VTA DA stimulation. The prior for this parameter was set to *k* ∼ *N* (0,1)*k* ∈ ℝ . Since not all sessions had trials with uncollected VTA DA stimulation rewards, for this analysis we only fitted sessions with at least 5 trials resulting in a CS+ without VTA DA stimulation. We then compared how close the posterior estimate for *k* in each session was to 1 (preset value of CS+ with VTA DA stimulation) and away from 0 (preset value for trials resulting in a CS–, Equation 2) .

## NEUROPIXELS RECORDINGS

### Recording strategy

We targeted all major striatal subregions and their associated cortical inputs and pallidal outputs. Recordings were done bilaterally, maximizing the number of distinct simultaneously recorded regions. To maximize the number of simultaneously inserted probes, all insertions were made with a combination of 10°, 15° and 30° zenith angles from probes positioned at 0°, 45°, 135° and 180° azimuth. Recording craniotomies (0.5 mm radius) across animals spanned from -1 to 3 mm A-P, and ± 0.2 - 4 mm M-L. Up to three recording sessions were performed per craniotomy; for each new session, the probe insertion site was moved by approximately 300 µm in the M-L and A-P axes, allowing us to easily associate probe histological reconstruction to recording/sessions. See Extended Data Table 4 for a summary of all craniotomies and recording trajectories tried and Extended Data Table 5 for a summary of regions targeted in each subject.

### Craniotomies

On the day prior to the first recording, animals were anesthetized, head-fixed and the UV glue covering the skull was carefully removed with a scalpel. The skull surface was cleaned with saline and dried with surgical cotton tipped applicators. Small craniotomies were made over the desired recording locations with a dental drill (¼ drill bit). No durotomy was performed. Craniotomies were covered with a small layer of Neomycin antibiotic ointment (Thermofisher) and the skull surface was then covered with a silicone elastomer (Kwikcast or Kwikseal, WPI). Up to 4 craniotomies were performed in each animal. Craniotomies could be penetrated with Neuropixels probes for up to 4 days. After this point, the dura hardened, preventing new insertions. Therefore, depending on the desired recording locations, craniotomy surgeries were separated in time (4-5 days apart) to maximize the amount of recordings possible per animal. A maximum of three craniotomy surgeries were made in each mouse^150,166,167^.

### Neuropixels insertion and recordings

All probes were grounded to the same ground pin that had been fixed during the headplating surgery into the cerebellum. Probes were connected to a Neuropixels PXI board (IMEC). The probes generated a 1 Hz square pulse sent to a I/O PXI module (National Instruments) to synchronize to behavioral and video streams. Recordings were acquired with the SpikeGLX software at 30 kHz. If noise levels were high (RMS >20 µV), grounding was set to internal referencing from the tip of the probes. All recordings were from the bottom 384 channels of each probe.

On each recording day, the tips of Neuropixels 1.0 probes (IMEC) were dipped in a drop of CM-DiI (1 µg/µl in ethanol) as previously described ^168^. Animals were head-fixed, the silicone elastomer was removed from the skull, and the craniotomies were rinsed with saline. Up to 4 probes targeting both hemispheres were inserted in each session. To avoid obstructing the visual field of adjacent craniotomies, probes were inserted in a two-stage process. First, all probes were lowered sequentially to a depth of ∼1 mm. Subsequently, each probe was advanced individually to its final target depth. Each probe was lowered into the brain at a speed of 2-5 µm/s using a micromanipulator system (uMp, Sensapex). Once all probes reached their final depth, to prevent the craniotomies from drying out and to further stabilize the probe, the skull was covered with Dow-Sil to fill the craniotomies and engulf the probe shanks. To minimize drift during the recording session, the probes were left in position at their final depth for 5-10 min before the start of the recording. After this waiting period, recording and the behavioral task (probabilistic reversal learning task) started, which lasted 60-90 minutes. At the end of the recording session, the probes were carefully and sequentially removed at a speed of ∼ 100 µm/s, the skull was cleaned, neomycin gel reapplied to the craniotomies and the skull was resealed with silicone elastomer.

### Spike sorting and manual curation

We used the standardized IBL pipeline for pre-processing and spike sorting ^150,169^. Briefly, raw electrophysiological recordings underwent three main pre-processing steps:

1. *Correction for ‘sample shift’ along the length of the probe by aligning the samples with a frequency domain approach*: This correction accounts for the small lag in between the sampling of each channel.
2. *Automatic detection, rejection and interpolation of failing channels on the probe*: Due to channel or tissue damage some probes channels are ‘dead’ or excessively noisy producing data that could corrupt the spike sorting procedure. This correction removes data from these channels.
3. *Application of a spatial ‘de-striping’ filter*: This correction removes noise common to several channels by applying a high-pass spatial DC filter prior to spike sorting.

After these preprocessing steps, spike sorting was performed using ibl-sorter^170^, a Python port of the Kilosort 2.5 algorithm^171^ that includes modifications to preprocessing. At this step, we applied registration, clustering, and spike deconvolution. Finally, data was manually curated with the Phy2 software (https://github.com/cortex-lab/phy). Manual curation focused on the removal of units that contained electrical noise or missing spikes (inferred based on a truncated distribution of spike amplitudes), and the splitting of clusters with clearly separable spike sources. Manual curation labels were then combined with automated metrics that differed for single units and multi-unit activity (MUAs).

### Inclusion criteria and pre-processing

For single units, we required the following values of our automated metrics: firing rate ≥0.01 sp/s, amplitude >30 µV, refractory period violations <10%, and spiking spanning at least 90% of trials in the behavioral session (defined as the fraction of total trials covered by the time from a neuron’s first to last spike). Single units were used for all analyses.

For MUAs, we required the following values: firing rate ≥0.01 sp/s, amplitude >30 µV, spiking in >75% of 10 s bins across the entire recording. All firing rates were estimated using only the time bins covered by the event kernels in the encoding model (described below). MUAs were only included in decoding analyses.

For all neural analyses, we binned and normalized each single unit’s spiking activity. First, we discretized each neuron’s spiking into 10 ms time-bins for each trial of the behavioral session with the bin edges aligned to the onset of the go cue. We then identified all of the time-bins that are covered by the event kernels in the encoding model (described below) and calculated the mean, *μ*, and standard deviation, *σ*, across those time-bins. We then *z*-scored the spiking activity for each neuron as:

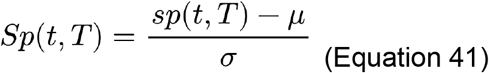

where *sp*(*t,T*) is the raw binned spiking activity at time *t* on trial *T* and *Sp*(*t,T*) is the normalized binned spiking activity.

For all neural analyses, we excluded the first and last ten trials of each behavioral session as well as any trials on which the animal did not make a choice or for which the time of wheel movement onset could not be identified post-hoc (described below).

For all neural analyses we used the raw firing rates from each neuron without applying any smoothing. For the PSTHs in Fig. 2.b,d and Fig. 3b,c, we applied a causal half-Gaussian filter with a 50 ms standard deviation to the neurons’ firing rates binned at 10 ms.

## NEURAL ANALYSES

### Neural encoding model

#### Basic encoding model

To determine how activity of single units was modulated by task events (Figs. 2–4 and Extended Data Figs. 3–5,7–9), we built linear encoding models to relate neural activity to each event^172,173^. We used multiple linear regression analyses of each neuron with the -scored 10 ms binned spiking activity as the dependent variable and behavioral events as the independent variables. To account for temporal offsets in the relationship between changes in spiking and behavioral events, independent variables were generated by convolving linearly spaced cosine basis sets (1 cosine per 25 ms of kernel duration) with binary vectors of event times (1 at the time of the behavioral event and 0 otherwise). To account for spike history effects^174^, we also included each neuron’s spike history as a set of additional independent variables that were generated by convolving a cosine basis set with log-scaling of the time axis (10 cosines spanning from 10 ms to 1 s in the past) with each neuron’s *z*-scored spiking activity. The cosine basis sets were generated with the MATLAB *raisedCosineBasis* package (https://github.com/pillowlab/raisedCosineBasis). The full model is:

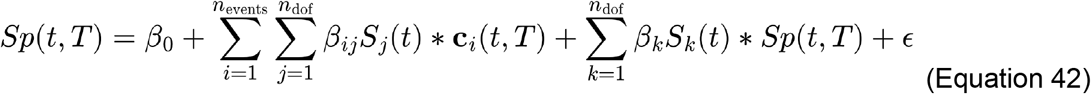

where *Sp*(*t,T*) is the *z*-scored spiking activity at time *t* on trial *T, β*_0_ is the intercept, *ϵ* is the noise, *n*_*events*_ is the number of behavioral events, *β*_*ij*_ is the regression coefficient for the *j*^th^ cosine basis function and *i*^th^ event, *S*_*j*_ is the *j*^th^ cosine basis function, *n*_*dof*_ is the number of cosines basis functions (i.e., degrees of freedom) for the *i*^th^ event, ***c***_*i*_(*t,T*) is a binary vector the same length as *Sp*(*t,T*) that is 1 at the time of event *i* and 0 otherwise, *β*_*k*_ is the regression coefficient for the *k*^th^ cosine basis function of the spike history kernel, *S*_*k*_ is the *k*^th^ cosine basis function of the spike history kernel, and * represents the convolution operation. Thus, spiking activity is modeled as the convolution of the event time vector ***c***_*i*_(*t,T*), with a temporal kernel, *K*_*i*_(*t*), summed across events:

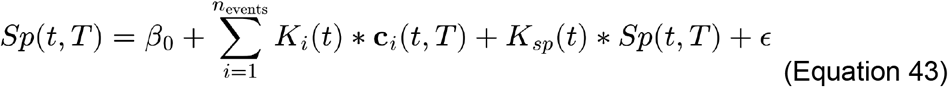

where the temporal kernel for event *i* is defined as:

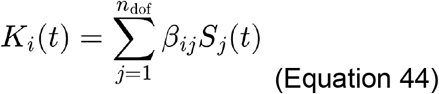

and the spike history kernel,*K*_*sp*_(*t*), is defined analogously.

We included temporal kernels for the following behavioral events: go cue (all trials), left movement onset (only left choice trials), right movement onset (only right choice trials), CS+ (only trials with CS+), laser stimulation onset (only trials with CS+ and laser stimulation), and CS– (only trials with CS–). The kernel for the go cue spanned from 0 to +1 sec. The kernels for left and right movement onset spanned from -0.5 s to +0.5 s. The kernels for CS+, laser stimulation onset, and CS– spanned from 0 to +2 sec. We estimated these temporal kernels with linear regression and lasso (*ℓ*_1_) regularization using the MATLAB *glmnet* package^175^ (https://github.com/junyangq/glmnet-matlab). The regularization parameter, *λ*, was chosen from a logarithmically spaced vector of 251 values from 10^-5^ to 10^5^ based on the minimum mean squared error using five-fold cross validation. For all encoding model analyses, trials were randomly assigned to cross validation folds with the constraint of approximately balancing block probability (left vs right high probability block), choice (left vs right), and reward (CS+ vs CS–) across folds.

We identified the onset of wheel movement using a strategy similar to ref. ^166^. First, we coarsely detected periods of movement as those with an average wheel speed (200 ms moving average) of ≥1 cm/sec. We next merged adjacent periods of movement that were separated by <100 ms and then removed periods of movement that were <50 ms duration. To precisely define movement onset times, for each movement detected above we identified the first moment at which the instantaneous wheel speed (5 ms moving average) was ≥0.2 cm/sec. The movement onset time for each trial was defined as the first movement onset time that occurred from 0.2 s before the time of the go cue to the time of the CS+ or CS–.

We classified choices and their associated Q-values as contralateral or ipsilateral with respect to each hemisphere of the brain based on the direction of wheel movement. For neurons on the left hemisphere, contralateral choices were defined as counterclockwise wheel movements (i.e., right choices) by the mouse and ipsilateral choices were defined as clockwise wheel movements (i.e., left choices). This is consistent with the definitions used for the IBL brain-wide map dataset^169^, which was collected in an identical recording and behavioral set up. Conversely, for recordings in the right hemisphere, contralateral choices were defined as clockwise wheel movements and ipsilateral choices were defined as counterclockwise wheel movements.

To identify significant modulation of neural activity by a particular event or set of events (Extended Data Fig. 3g–k; details for specific models below), we calculated an *F*-statistic comparing the fit of the full model to that of a reduced model refit without the predictors for that event. The *F*-statistic was calculated by comparing the predicted spiking activity, from five-fold cross validation, to the real spiking activity of each model:

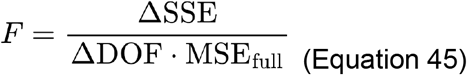

where ΔSSE is the difference in sum squared error between the full and reduced models, ΔDOF is the difference in degrees of freedom (i.e., number of predictors) between the full and reduced models, and MSE_full_ is the mean squared error of the full model. Because the binned spiking data could be autocorrelated and therefore not i.i.d., we calculated *P*-values by comparing these *F*-statistics to those from a null distribution that preserved much of the same autocorrelation^89^. To do this, 200 instances of null spiking data were generated by circularly shifting the binned spiking activity from each session by a random integer (noting the problems with this described in ref. ^89^). ^89^e did not adopt a “pseudosession” approach^89,90^ here because the time-course of the neural responses to task events should be less susceptible to drift across a session (that may lead to spurious significance) than the trial-by-trial-modulation in amplitude or gain of the neural response (described below). *P*-values were calculated as the fraction of *F*-statistics from the null dataset that were greater than the real *F*-statistic. Neurons with *P* ≤were considered to significantly encode that event.

We first used this strategy to identify all neurons that were significantly modulated by any task event. We compared the full model to a reduced model with all behavioral kernels removed and only the spike history kernel and intercept remaining. For all downstream encoding analyses, we only included neurons for which the full model was significantly better than this reduced model (Extended Data Fig. 3d; referred to as “task-modulated” in the main text; 4,036 out of 5,109 (79.0%) striatal single units).

We then used this strategy to classify neurons as significantly encoding (1) go cue (Extended Data Fig. 3g), by removing only the go cue kernel in the reduced model; (2) movement (Extended Data Fig. 3h), by removing both the left and right movement onset kernels; (3) outcome (Extended Data Fig. 3i), by removing the CS+, laser stimulation, and CS– kernels; (4) choice (Extended Data Fig. 3j), by replacing the left and right movement onset kernels with a single movement onset kernel that was not choice-specific; and (5) reward (Extended Data Fig. 3k), by replacing the CS+ and CS– kernels with a single CS kernel that was not reward-specific and also removing the laser stimulation onset kernel. We also used this strategy to specifically examine encoding of the CS+ by comparing the full model to a reduced model with only the CS+ kernel removed.

#### Bilinear encoding model

To determine how the transient, event-related responses of striatal neurons were modulated by trial-by-trial fluctuations of the decision variables described above, we fit bilinear models that include multiplicative gains that scale the temporal kernels for each event on a trial-by-trial basis by the value (or decision propensity) of that trial ^42,80^. The trial-by-trial gain for event *i* is:

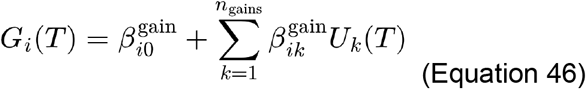

where 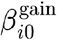 is the gain offset for event *i,n*_*gains*_ is the number of trial-by-trial variables, 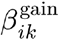 is the gain coefficient for trial-by-trial variable *k* and event *i* $, and *U*_*k*_(*T*)is the value of trial-by-trial variable *k* (for example, Δ*Q* or *V*) on trial *T*. Thus, the spiking activity at time *t* on trial is modeled as.

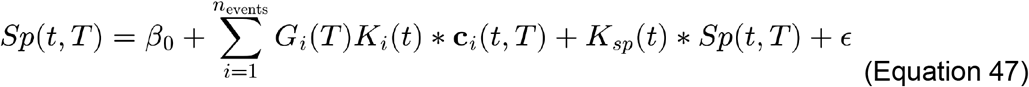

Where *c*_*i*_(*t,T*)is a binary indicator of event times for event *i* and *K*_*i*_ (*t*) and *K*_*sp*_(*t*) are the behavioral event and spike history kernels described above.

To fit the bilinear model defined in Equation 47, we iteratively estimated the temporal kernels and intercept – *K*_*i*_ (*t*), *K*_*sp*_(*t*), and β_0_ – and the value-dependent gains – *G*_*i*_.(*T*) On each iteration, we first kept the value-dependent gains fixed and estimated the temporal kernels and intercept with linear regression and lasso (ℓ_1_) regularization using the MATLAB *glmnet* package. After fitting, the temporal kernels were normalized by dividing the coefficients for the cosine basis set, β_*ij*_, by the Euclidean norm of the kernel. On the first iteration, *G*_*i*_.(*T*) were all set to 1 and on subsequent iterations were based on the gain coefficients estimated in the previous iteration. We next kept the temporal kernels and intercept fixed and estimated the value-dependent gains with linear regression and ridge (ℓ_2_) regularization using the same MATLAB package. When fitting the value-dependent gains, the intercept, β_0_, and spike history term, *K*_*sp*_(*t*) ∗*Sp*(*t*), fitted in the previous step were provided to the model as offset terms. The regularization parameters, λ, for both steps were chosen from a logarithmically spaced vector of 251 values from 10^-5^ to 10^5^ based on the minimum mean squared error using five-fold cross validation on the first iteration and then held constant at the same values for the remaining iterations. This process was repeated until the model converged (defined as three consecutive iterations for which every *β* changed by ≤0.001 compared to the previous iteration) or for 100 iterations.

We fit three separate bilinear models to evaluate the influence of:

1. Contralateral and ipsilateral choice value from the Extended Q-learning model with the CS+ as reward (Model 1;Figs. 2,4; Extended Data Fig. 4),
2. Relative and state value from the Extended Q-learning model with the CS+ as reward (Model 1; Fig. 3; Extended Data Fig. 7,8).
3. Choice propensity, CS+ value, and choice-state value from the Conditioned Reinforcer model (Model 6; Fig. 6; Extended Data Fig. 12).

In the contra/ipsi value encoding model (Fig. 2), the trial-by-trial variables were:

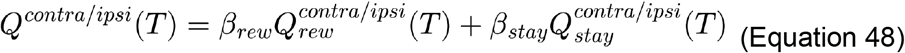

In the relative/state value encoding model (Fig. 3), the trial-by-trial variables were:

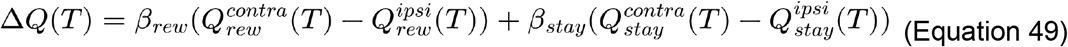

and *V*(*T*)from Equation 36.

In the Conditioned Reinforcer encoding model, the trial-by-trial variables were:

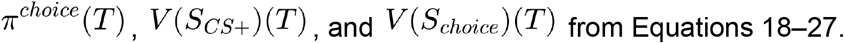

Each gain was permitted to multiply every kernel, and so there were a total of 18 predictors (2 trial-by-trial variable gains and 1 gain offset each for 6 kernels) in the contra/ipsi value and relative/state value encoding models and 24 predictors (3 trial-by-trial variable gains and 1 gain offset each for 6 kernels) in the Conditioned Reinforcer encoding model.

We next compared the full model with all trial-by-trial variable gains present to reduced models with some or all of these gains removed. For both the full and reduced models, the gains were refit using the kernels from the final iteration of the bilinear encoding model. The regularization parameter,λ, for ridge (ℓ_2_), was chosen independently for each model from a logarithmically spaced vector of 251 values from 10^-5^ to 10^5^ based on the minimum mean squared error using five-fold cross validation.

For the bilinear model with *Q*^*contra*^ and *Q*^*ipsi*^, we fit reduced models with either or both of these trial-by-trial variables removed from every kernel (three reduced models total). For the bilinear model with Δ*Q* and *V*, we fit reduced models with either of these trial-by-trial variables removed from every kernel (three reduced models total). For the Conditioned Reinforcer model (π^*choice*^,*V*,(*S*_*cs+*_),and *V*(*S*_*choice*_), we fit reduced models with each of these three trial-by-trial variables removed from every kernel. To quantify how strongly each neuron encoded a particular trial-by-trial variable or combination of variables, we calculated the difference in cross-validated variance explained, ΔR^2^, by the full model compared to the reduced models. We also used the residuals (i.e., the difference between the real and predicted firing rates) from these reduced models as the basis for the decoding analyses in Fig. 6k,l,p,q (see Neural Decoders section of the Methods).

To identify significant modulation of neural activity by a particular trial-by-trial variable or combination of variables (Figs. 2–4; Extended Data Fig. 4,7,8), we calculated an *F*-statistic comparing the fit of the full and reduced models as in Equation 45 and then compared this *F*-statistic to an appropriate null distribution. To generate the null distribution, we then calculated the same *F*-statistic for the same comparison after refitting the full bilinear model and reduced models for 200 instances of pseudosession data. Our goal was to preserve the temporal auto-correlations within the neural data as well as within the trial-by-trial value estimates — while breaking the temporal relationship between these two variables in the pseudosessions — to control for spurious correlations and significance in our encoding model that could potentially arise due to slow but unrelated drift in spiking and/or value^89,90^. These pseudosessions were created by concatenating the trial-by-trial variables from all behavioral sessions and then randomly selecting chunks of consecutive trials the same length as the real behavioral session; the event times from the real behavioral session were used when fitting the models. The same pseudosessions were used for every neuron. The regularization parameter, *λ*, for ridge (ℓ_2_) was chosen independently for each full or reduced model and for each instance of pseudosession data. *P*-values were calculated as the fraction of *F*-statistics from the null dataset that were greater than the real *F*-statistic. Neurons with *P* ≤ 0.01 were considered to significantly encode that trial-by-trial variable or combination of variables. Similarly, we report the change in variance explained for each model comparison as corrected ΔR^2^, in which the mean ΔR^2^ across all pseudosessions in the null distribution is subtracted from the ΔR^2^ fit to the real behavioral data.

For the Conditioned Reinforcer encoding model specifically, we included an additional correction to generate valid pseudosessions. Because *V*(*S*_*cs+*_) and *V*(*S*_*choice*_) are initialized to different values at the start of each session, concatenating them end-to-end as described above would introduce an artificial jump or discontinuity at each session boundary. To prevent this, before concatenating sessions we shifted each session’s *V*(*S*_*cs+*_) or *V*(*S*_*choice*_) trace by a constant offset equal to the difference between the last trial’s value in the preceding session and the first trial’s value in the current session.

#### Learning rate analysis

In Fig. 4, we used a variant of the bilinear encoding model to test how the learning rates in the behavioral model (α_*rew*_ and γ^*forget*^) impact neural encoding of estimated trial-by-trial 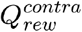 and 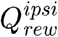 values from the Extended Q-learning model with the CS+ as reward (Model 1). These learning rates only impact the reward and not stay component of the Q-values, so we considered only the contribution of reward to action value (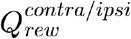) rather than the full estimate (*Q*^*contra/ipsi*^ from Equation 48) in this encoding analysis.

First, we calculated 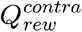 and 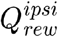, values for every trial of every session using Model 1 but with the α_*rew*_ learning rate parameter set to a new value (γ^*forget*^ remains fixed at the original value). We then refit trial-by-trial gains for each neuron using the kernels from the final iteration of the bilinear encoding model described above but replacing the trial-by-trial 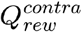 and 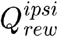 values with those refit using the new α_*rew*_ value. The regularization parameter, λ, for ridge (ℓ_2_) was chosen independently for each value of the learning rate α_*rew*_ from a logarithmically spaced vector of 251 values from 10^-5^ to 10^5^ based on the minimum mean squared error using five-fold cross validation. We also refit reduced models with either or both of the and Δ_*rew*_ *V* gains removed for every kernel, refitting the regularization parameter independently for each reduced model. We then calculated the difference in cross-validated variance explained, ΔR^2^, for the model with either the 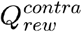 or the 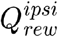 gains present compared to the reduced model with no trial-by-trial gains. We then repeated this analysis for 40 linearly spaced values of α_*rew*_ ranging from 0.025 to 1 (Fig. 4c,d, Extended Data Fig. 8a,b).

We used an analogous approach to test how γ^*forget*^ varying impacts neural encoding of the estimated trial-by-trial 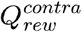 and 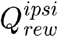 values. In this case, we repeated the analysis above for 40 linearly spaced values of γ^*forget*^ ranging from 0.025 to 1 with Δ_*rew*_ remaining fixed at the original value (Fig. 4e,f, Extended Data Fig. 8c,d).

We repeated this entire process for the relative/state value encoding model using the estimated trial-by-trial 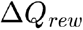 and *V* values as gains (Extended Data Fig. 8e–g). Here we once again considered only the contribution of reward, and not the full estimate including the stay parameter.

We found that estimates of *V*,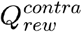, and 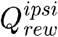 each monotonically increased throughout a behavioral session for low learning and forgetting rates, which could result in spurious correlations with slow but unrelated drift in spiking across a session. Therefore, for all encoding analyses above we detrended the raw estimates of *V*,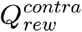, and 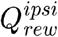 before fitting the encoding models. Specifically, we fit a linear regression of trial number (independent variable) and *V*,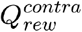, or 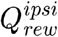 (dependent variable) using the MATLAB *fitlm* function and then used the residuals from this model as the trial-by-trial variable in the encoding models described above. We performed this operation separately for *V*,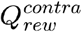,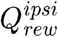 and derived from each value of α_*rew*_ or γ *forget*.

We used a pseudosession approach, analogous to that described above for the bilinear encoding model, to control for spurious correlations and significance in our encoding model that could potentially arise due to slow but unrelated drift in spiking and/or value^89,90^. These pseudosessions were created by concatenating the trial-by-trial variables from all behavioral sessions and then randomly selecting chunks of consecutive trials the same length as the real behavioral session. The same trial order was used for the pseudosession for each value of α_*rew*_ and γ*forget*. We then report the change in variance explained for each model comparison as corrected ΔR^2^, in which the mean ΔR^2^ across all pseudosessions in the null distribution is subtracted from the ΔR^2^ fit to the real behavioral data.

#### Statistical comparisons across brain regions

We used the matlab *chi2cont* function (https://www.mathworks.com/matlabcentral/fileexchange/45203-chi2cont) to compare the proportion of significantly value-modulated neurons across pairs of brain regions (Fig. 2g–i, Fig. 3d,e, Fig. 6i,n, Extended Data Fig. 12b) or within M/L or D/V bins of intermediate CP (Fig. 3j) with χ^2^ tests. We used the built-in matlab *signrank* function to compare the strength (ΔR^2^) of value-modulation across intermediate and caudal CP (Fig. 6j,o, Extended Data Fig. 4a, Extended Data Fig. 7e and Extended Data Fig. 12c) or within M/L or D/V bins of intermediate CP (Extended Data Fig. 7m) with Mann–Whitney U tests. Detailed results of all statistical comparisons are shown in Extended Data Table 7.

### Neural decoders

To investigate the neural representation of the modeled behavioral variables, we employed a decoding approach. We trained linear decoders to predict four key variables derived from our computational models: Contralateral choice value (*Q*^*contra*^) and Ipsilateral choice value (*Q*^*ipsi*^) (Fig. 2j-n; Extended Data Fig. 4f-i,5), Relative Value (Δ*Q*) and State Value (*V*) (Fig. 3f,g; Extended Data Fig. 7i,j)), and, from the Conditioned Reinforcer model, the learned CS+ value (*V*(*S*_*cs+*_), the choice state value (*V*(*S*_*choice*_)) and the choice propensity ((π^*choice*^,) (Fig. 6k,l,p,q; Extended Data Fig. 12d).

Decoders were trained using varying numbers of neurons (curated units and multiunits) to assess the robustness of the neural encoding, with combinations of 10, 20, 30, and 40 neurons. For each *n neurons* decoder, the decoding procedure was run 100 times on random combinations of neurons and summarized by its mean performance. All neurons selected for a given decoder were drawn from simultaneously recorded neurons from the same striatal or cortical subregion (for subregion designation for all decoding analyses please see Extended Data Table 3). Neural activity was binned into 100 ms intervals for decoding, with a separate decoder trained for each time bin. The temporal windows for decoding were chosen to align with relevant task events: for State Value (*V*), neural activity was analyzed from 0.5 s *before* the outcome event to 2 s *after* the outcome. Critically, we predicted the pre-update value (i.e., the value before the reward was obtained on that trial). For Relative Value (Δ*Q*), activity was analyzed from 0.5 s *before* the Go cue to 0.5 s *after* the Go cue. To mitigate potential confounds arising from the inherent correlation of Relative Value with choice and State Value with outcome, only trials where the mouse made a contralateral choice were included for Relative Value decoding and only rewarded trials were used for State Value decoding (Fig. 3f,g; Extended Data Fig. 7i,j,p,q).

In Fig. 6k,l,p,q (Conditioned Reinforcer model), because *V*(*S*_*cs+*_) and *V*(*S*_*choice*_) are correlated, decoding either variable directly from raw activity would be confounded by neuronal firing attributable to the remaining variables. Therefore, to decode *V*(*S*_*cs+*_) or *V*(*S*_*choice*_)specific neural variance, we instead used the reduced model described in the encoding model section, in which the variable of interest’s gain was removed but the remaining two gains were retained. We computed the residual spiking activity for each neuron by subtracting this reduced model’s prediction from the observed activity, and used these residuals as input to that variable’s population decoder. Because the CS+ kernel in the linear encoding model is only defined over a 0 to 2 s window following CS onset, residual activity and consequently the decoders for these variables were restricted to this window. Additionally, since the encoding model was only fit for well isolated single units, no multiunit activity was used for this family of decoders.

Our decoders predicted trial by trial value at each time point based on the population activity using linear regression with Lasso (L1) regularization and an intercept. We performed 5-fold nested stratified cross-validation ^176^ to evaluate model performance and optimize regularization strength (λ) for each decoder (i.e., for each time bin and neuron combination). The procedure consists of an inner and outer loop both having 5 stratified folds. The outer folds are used for model evaluation and the inner folds for λ selection (λ= {10^−6^,10^−5^,10^−4^,10^−3^,10^−2^,10^−1^,10^0^,10^1^}). Briefly, (i) trials were first discretized into bins of uniform width using a Freedman-Diaconis rule based on the trials value (e.g. *Q*^*contra*^). Once trials were divided in bins, (ii) trials within each bin were shuffled and (iii) divided into 5 outer folds with a similar number of trials from each bin. One fold serves as a test set for evaluation and the other four make up the training set. (iv) For each outer training set, an inner cross-validation procedure is used to select an optimal λ. This inner cross-validation procedure also consists of 5 folds. A decoder was trained with a specific λ on 4 of the inner folds and validated on the remaining inner fold. This was repeated 5 times, and the average validation score across the 5 inner folds is stored. This was repeated for every tested λ. From this, a winner λ (best R^2^) is selected. (v) Finally, the model is trained with the winner λ in the entire outer training set and evaluated on the held-out outer test set. Steps iv-v are repeated 4 more times for the remaining outer fold sets. The final reported performance is the average score across the 5 outer test sets.

To account for nonsense correlations ^89^ we corrected the R^2^ values reported in all time-resolved decoders (Fig. 2k-n, Fig. 3f,g, Extended Data Fig. 5, Fig. 6k,l,p,q, and Extended Data Fig. 12d) by subtracting the mean R^2^ obtained from decoding a null distribution consisting of 200 “pseudosessions” from the R^2^ on the real session^169,177^. These pseudosessions for the decoders in Fig. 2-3 and Extended Data Fig. 7 were created by concatenating all actual Qcontra, Qipsi or State or Relative values from all experimental sessions and then randomly selecting chunks of the same length as the original session data. For the decoders in Fig. 6 and Extended Data Fig. 12 where we analyzed the behavioral variables from the Conditioned Reinforcer model, we included an additional correction to generate valid pseudosessions. Since *V*(*S*_*cs+*_)and *V*(*S*_*choice*_) are initialized to different values at the start of each session, concatenating sessions end-to-end would create discontinuities at the session boundaries. To avoid this, we offset each session’s *V*(*S*_*cs+*_) or *V*(*S*_*choice*_) trace by a constant before concatenation, equal to the difference between the last trial’s value in the preceding session and the first trial’s value in the current session. To compare performance across regions (Fig. 2k-n, Fig. 3f,g, Fig. 6k,l,p,q, and Extended Data Fig. 12d) we statistically compared the corrected R^2^s obtained from running the decoders in NAc vs all other CP subregions.

## ALLEN ATLAS PROJECTION INTENSITY ANALYSIS

For Fig. 3 k-m, we analyzed cortical viral injection experiments and projection images from the Allen Institute’s Mouse Connectivity Atlas (http://connectivity.brain-map.org/) ^97^ to determine which cortical areas projected to different striatal subregions and domains, and to correlate those patterns with the encoding of state and action value.

We queried the API for experiments from wild-type mice (n = 58) and several Cre lines targeting specific cortical layers. In selecting Cre lines, we focused on lines that target layer 5 and layers 2/3. Lines that instead target layer 4 and layer 6 were excluded because these layers do not project to striatum ^97,178^. The Cre lines analyzed were: A930038C07Rik-Tg1-Cre (targets layer 5b neurons, n = 30) ^179^, Rbp4-Cre_KL100 (targets layer 5 pyramidal neurons, n = 43 experiments) ^97,180^, Tlx3-Cre_PL56 (targets layer 5a IT neurons, n = 27 experiments) ^97,180^, Cux2-IRES-Cre (targets layers 2/3 and 4, n = 39 experiments) ^97,181^, Emx1-IRES-Cre (targets cortical excitatory neurons, n = 8 experiments) ^97,182^, and Grp-Cre_KH288 (targets layers 2/3 and 4, n = 14 experiments) ^97,180^. The full list and details of used experiments selected for analysis can be found in Extended data table 6.

For each experiment, we first normalized projection intensity by dividing by the total injection intensity in the source cortical structure. Specifically, we used data provided by the Allen Institute to normalize: the value ‘sum_pixel_intensity’ given by structure_unionizes for the injection region (https://allensdk.readthedocs.io/en/latest/unionizes.html). We grouped injections together by region (PFC, ACC, OFC, Insula, SS and MO), and averaged across experiments.

For Fig. 3k, we then scaled the mean projection intensity from 0 to 1 at the voxel level on this average data for each source region and subdivided the striatum into the 31 different domains. For each striatal domain, we averaged the normalized mean projection intensity in that subregion’s voxels. For each cortical area, we then quantified the correlation (Pearson *r*) between the mean projection intensity in each domain versus the percentage of neurons significantly encoding state or relative value in each domain for which we had recorded at least 20 neurons (n = 25 domains).

For Fig. 3l,m visualizations of projections from source data in MO and OFC, we averaged across allen atlas slices at the AP locations 48, 49 and 50 ARA (spanning 200 µm), corresponding to the subregion CP-int.

## FIBER PHOTOMETRY RECORDINGS

We simultaneously recorded rGRAB-DA^183^ signals from dorsolateral striatum (DLS) and nucleus accumbens core (NAcc) with a multi-fiber photometry system (FP3002, Neurophotometrics) at the same time we stimulated VTA DA cell bodies expressing ChR2 in naive mice not performing any task (Extended Data Fig. 6). We waited a minimum of 1 month from viral injection for expression before starting photometry recordings. Stimulation and imaging were controlled with the Bonsai Neurophotometrics module. The photometry system consisted of an LED for excitation (560 nm light of 10 ms width pulses at 20 Hz) and a CMOS camera acquiring fluorescence emissions. The camera acquisition epochs were synchronous with the LED flashes. We used a low autofluorescence patch cord with 3 bundled branches (Ø200/220 mm fiber core/cladding diameter, 0.37 NA, 1.25 mm ferrule diameter, Doric) to be able to image DLS and NAcc simultaneously (one branch was unused). On the recording day and prior to the recording, we passed 0.5 mW 470 nm light through the patch cord for 1 hour in order to photobleach autofluorescence within the patch cord, and improve recording quality. 1s light trains of optogenetic stimulation (20 Hz, 8 mW, 447 nm, 5ms pulse width) was delivered bilaterally with a separate patch cord (Ø200/220 mm fiber core/cladding diameter, 0.39 NA, 2.5 mm ferrule diameter, Doric) connected to optical fibers located over the VTA (0.39 NA, Ø300 mm core, 2.5 mm ferrule, Thorlabs). To prevent optical bleedthrough from the stimulation light to the recording system, imaging and stimulation were interleaved such that optogenetic stimulation occurred in the periods that the CMOS was not acquiring

Fluorescence signals recorded during each session from each location were transformed to ΔF/F_0_ using the following formula:

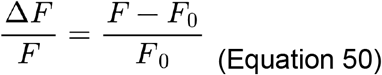

F_0_ signal was the ± 30 s rolling average of the raw fluorescence signal. Finally, ΔF/F_0_ signals were z-scored per-session, using a mean and standard deviation calculated based on all the data from each session.

## HISTOLOGY

Mice were anesthetized with pentobarbital sodium (2 mg/kg, Euthasol) and transcardially perfused first with 10 ml of ice-cold phosphate buffered saline (PBS) followed by 30 ml of 4% paraformaldehyde (PFA) in PBS. Brains were then dissected and post-fixed in 4% PFA overnight at 4°C. After fixation, brains were either: a) shipped to the Sainsbury Wellcome Centre for whole brain serial sectioning 2-photon microscope (for Neuropixels experiments in Figs. 1–5, Extended Data Figs. 1-5,7-10) or b) imaged with an automated slide scanner (NanoZoomer S60, Hamamatsu) (for photometry experiments in Extended Data Fig. 6).

### Whole-brain serial two photon imaging

Brains were embedded in agarose and then serially sectioned and imaged using a 4 kHz resonant scanning two-photon system immersed in 50 mM PBS ^184,185^. The microscope was operated with ScanImage Basic (Vidrio Technologies, USA), and imaging parameters were configured through BakingTray, a custom-developed software (https://github.com/SWC-Advanced-Microscopy/BakingTray). Individual image segments were stitched together into 2D planes with StitchIt (https://github.com/SWC-Advanced-Microscopy/StitchIt). Complete coronal brain image stacks were imaged at a resolution of 4.4 x 4.4 x 25.0 µm (XYZ) using a Nikon 16x objective (NA 0.8) and a 920 nm laser with approximately 150 mW power at the tissue. A vibrating blade microtome (VT1000 vibratome, Leica) created 50 µm slices. Optical imaging within each slice captured two focal planes at depths around 30 µm and 55 µm from the top surface with a piezo objective scanner (PIFOC, PI). Two fluorescence channels were simultaneously recorded using Hamamatsu R10699 multialkali PMTs: ‘Green’ (525 nm center, 50 nm bandwidth; Chroma ET525/50m) and ‘Red’ (570 nm long-pass filter; Chroma ET570lp).

### Registration and localization of Neuropixels probes and optical fibers from whole-brain stacks

The obtained whole brain images from serial two-photon imaging were first downsampled to 25 µm isotropic voxels. Then, they were spatially registered to the adult mouse Allen Common Coordinate Framework ^186^ via BrainRegister (https://github.com/stevenjwest/brainregister), a registration procedure based on elastix ^187^ with optimized settings for mouse brain data. Samples were registered to the CCF template image and the CCF template was also registered to the sample.

Neuropixels probe tracks were then manually traced on image stacks registered to the Allen CCF using the python package Lasagna (https://github.com/SainsburyWellcomeCentre/lasagna). Tracing was performed on merged stacks of the autofluorescence and CM-DiI channel using coronal and horizontal views. Neuropixels channels were mapped to the traced tracks (now in Allen CCF space) and further aligned based on electrophysiological features using a custom alignment tool ^188^.

While the Allen CCF groups all caudoputamen regions into a single entity, we required finer anatomical resolution for precise localization of our recording sites. To achieve this, we utilized a striatum atlas ^92^, which is based on the cortico-striatal projectome ^91^. This specialized atlas, also defined within the AllenCCF space, subdivides the caudoputamen into 27 distinct CP “domains”. These domains are further organized into “subregions” (Rostral, Intermediate, and Caudal) based on their anterior-posterior positions (for details see Fig. 2e, Extended Data Fig. 3.). To assign our Neuropixels recording channels to these domains, we located each channel’s closest 100 µm coronal slice in the specialized striatal atlas and assigned the corresponding anatomical label for that M-L x D-V location. As both atlases are in AllenCCF space, no further alignment was necessary.

For locating optical fibers in mice doing the probabilistic learning task (Extended Data Fig. 1), optical fiber location was determined as the coordinates of the center of the lesion tip in the whole-brain stacks registered to the AllenCCF.

### Cryostat sectioning and Slide scanning (Nanozoomer)

Brains were dehydrated in 30% sucrose for cryoprotection. The brain was mounted within OCT and cryosectioned in 50 µm slices with a cryostat. Sections were mounted with DAPI Fluoromount-G (Southern Biotech) and imaged with an automated slide scanner (NanoZoomer S60, Hamamatsu).

### Localization of optical fibers from Nanozoomer stacks

Coronal cryosections sections containing the most ventral lesion made by the optical fiber were manually aligned to the striatum atlas (see previous section). Alignment was based on anatomical landmarks such as the thickness and shape of the corpus callosum and size of the lateral ventricles. The lesion tip was used as the reference coordinates for fiber or injection location.

## STATISTICS

No statistical methods were used to predetermine sample sizes. Sample sizes were chosen based on previous studies^53,80,166^ and on the availability of animals. Replication: All attempts at replication were successful. Most experiments were replicated in multiple independent cohorts of animals with all experimental groups present in each cohort. We used multiple independent experimental approaches, and multiple independent analyses within each experiment, to confirm our findings whenever possible. Randomization: For experiments with multiple groups, individual animals or entire cages were randomly assigned to a group either at the time of surgery or at the beginning of behavioral testing Blinding: Automated analyses and experimental hardware and software, without manual intervention, were used whenever possible. Experimenters were not blinded to the group assignments of the animals. Statistical tests and data analysis were performed in MATLAB (R2021a), Stan (2.36) and Python (3.8.10) as described above.. Individual data points are shown when practical (and always for n ≤ 10), and box plots show the data distribution for larger sample sizes. Sample sizes, statistical tests, exact P-values, error bars, shading, and box plots are defined in the figure captions, Methods, and Extended Data Table 7.

### Single neuron statistics

To determine if neurons are significantly modulated by *any* task event, we first compared the full linear encoding model for each neuron to a reduced model with all behavioral kernels removed. We then used a similar strategy to classify neurons as significantly encoding *individual* task events, by comparing the full linear encoding model for each neuron to reduced models with the relevant behavioral kernels (e.g., the go cue kernel or the action kernels) removed. For each statistical test, we compared the *F*-statistic from modeling the real spiking and behavioral data to a null distribution of *F*-statistics that was created for each neuron using circularly shifted spiking data. We considered neurons with *P* ≤ 0.01 to be significantly modulated by a given task event. See the “Basic encoding model” section of the neural encoding model Methods for details.

To determine if neurons were significantly modulated (*P* ≤ 0.01) by trial-by-trial estimates of value, we compared the full bilinear encoding model for each neuron to reduced models with the relevant trial-by-trial gains (e.g., and/or) removed. For each statistical test, we compared the *F*-statistic from modeling the real spiking and behavioral data to a null distribution of *F*-statistics that was created for each neuron using pseudosessions of behavioral data. We considered neurons with *P* ≤ 0.01 to significantly encode value. See the “Bilinear encoding model” section of the neural encoding model Methods for details.

All other statistical comparisons are described in the “Statistical comparisons across brain regions” section of the neural encoding model Methods.

### Decoding model statistics

To statistically compare the decoding performance for the different studied value variables (e.g.*Q* ^*contra*^) between CP regions and the NAc, we compared the null corrected R^2^ (see Methods: Neural Decoders for details on the null correction) for every *Subregion x Session x Time point* using a Mann-Whitney U rank sum test. For comparing decoding performance for different value variables (*Q*^*contra*^ vs.*Q* ^*contra*^) within a region, we used the same approach but using a paired Wilcoxon signed rank test instead.

## DATA AND CODE

All data is available through the International Brain Laboratory interface and data repository (https://www.internationalbrainlab.com/data). Instructions on how to download the data can be found at: https://github.com/alejandropan/panzimmerman_VTA_DA. Final code for every visualization and analysis will be released upon publication on GitHub.

## ACKNOWLEDGEMENTS

We thank E. Engel, O. Huang, A. Chan and staff at the PNI Viral Core Facility for AAV production; A. Sirko and staff at the Princeton Laboratory Animal Resources for help with animal husbandry; members of the Witten lab, Dayan lab for help and feedback; and the International Brain Laboratory for support, particularly the project board (Matteo Carandini and Kenneth Harris). Funding was provided by the Wellcome Trust (216324 to IBL, P.D., 221657/Z/20/Z to A.P.V.), the Helen Hay Whitney Foundation (to C.A.Z.), the Max Planck Society (to P.D.), the Alexander von Humboldt Foundation (to. P.D.), the Simons Collaboration on the Global Brain (to IBL, I.B.W. and P.D.), the Howard Hughes Medical Institute (to I.B.W.) and the National Institutes of Health (K99-DA059957 and R00-DA059957 to C.A.Z.; U19-NS123716 and DP1-MH136573 to I.B.W.).

## AUTHOR CONTRIBUTIONS

Conceptualization, A.P.-V., C.A.Z., P.D., and I.B.W.; data collection, A.P.-V., B.M., M.L., T.J. and S.J.W.; methodology, A.P.-V., C.A.Z., P.D., and I.B.W.; data curation, A.P.-V., C.A.Z. and M.F.; formal analysis, A.P.-V., C.A.Z. and J.M.J.F.; software, A.P.-V., C.A.Z., M.F. and J.M.J.F.; visualization, A.P.-V., C.A.Z. and J.M.J.F.; writing, A.P.-V., C.A.Z., P.D., and I.B.W.; supervision, A.P.-V., C.A.Z., P.D., and I.B.W.; funding acquisition, A.P.-V., C.A.Z., P.D., and I.B.W.

---

**Extended Data Figure 1.**
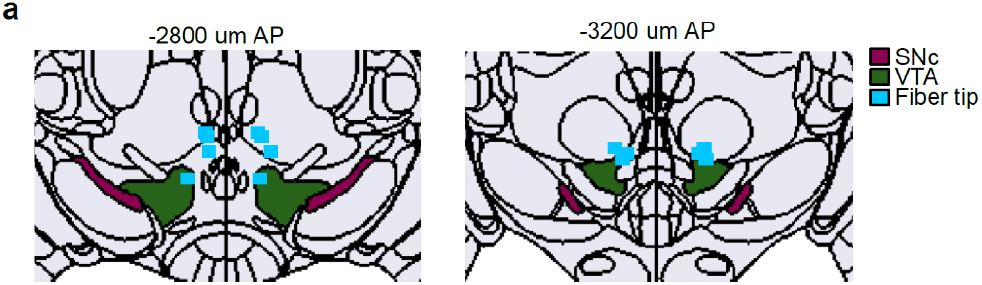
VTA stimulation histology. **a**, Histological reconstruction of optogenetic fiber tip locations (blue rectangles) for VTA DA stimulation projected to the nearest 400-µm slice of the Allen CCF (all mice in Figs. 1–4).

**Extended Data Figure 2.**
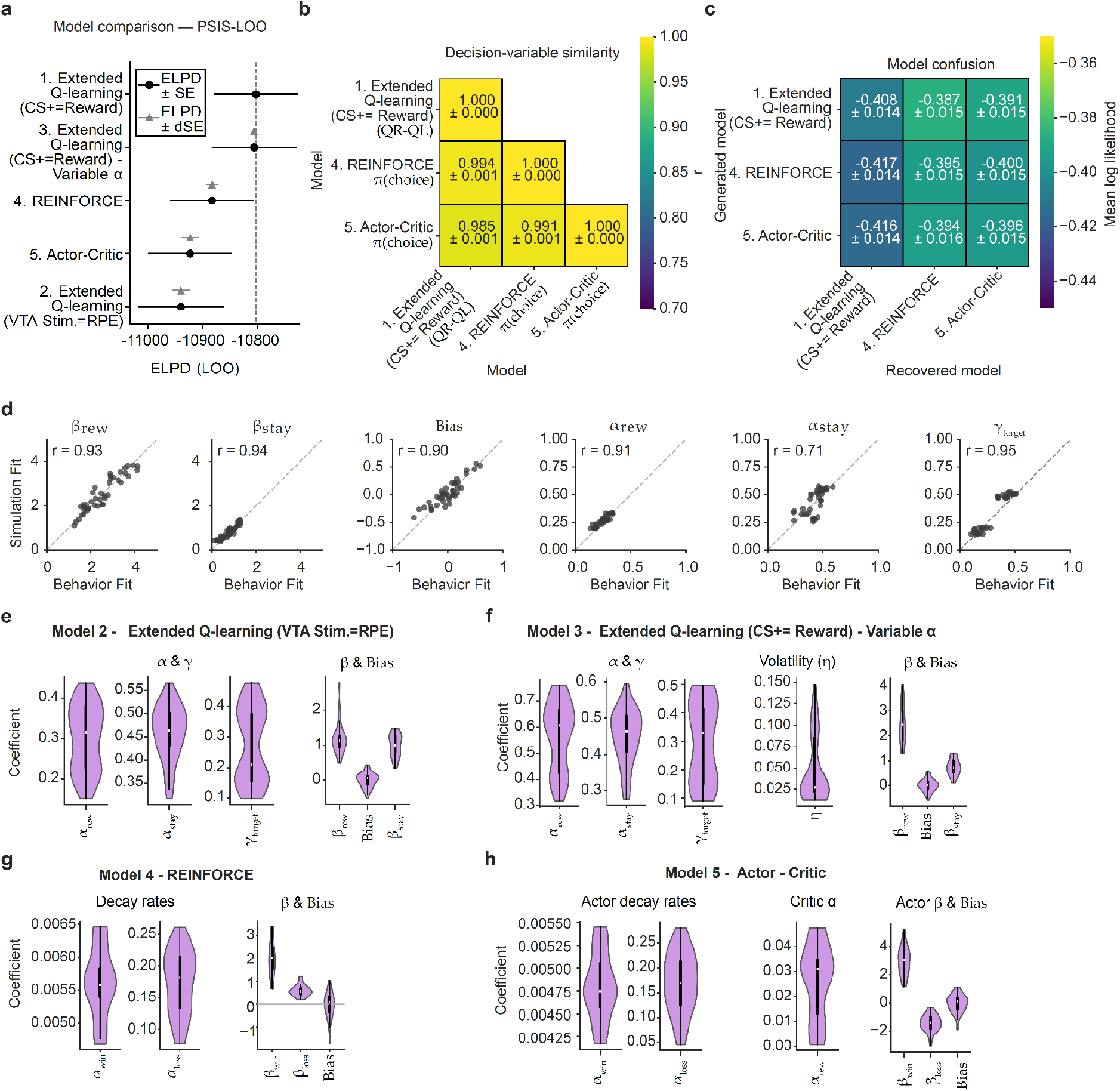
The Q-learning model parameters are recoverable, but the model’s behavior is indistinguishable from that of REINFORCE and Actor-Critic models. **a**, Model comparison by PSIS-LOO, ranked by expected log pointwise predictive density (ELPD), for which greater values indicate superior out-of-sample predictive performance. Both markers show each model’s ELPD; only the error bars differ. Circles: ± s.e. of the ELPD. Triangles: ± s.e. of the trial-wise predictive difference from the best model (dSE). n = 22,406 trials, n = 8 mice, n = 41 sessions.difference from the best model (dSE). All models predicted held-out choices within 0.5% of one another (cross-validated mean predicted probability of the observed choice, 67.8–68.3%). n = 22,406 trials, n = 8 mice and n = 41 sessions. **b**, Similarity between the decision variables from every model: Relative value (*Q*^R^ −*Q*^*L*^) for Extended Q-learning models, and propensities *(π*) for REINFORCE and the Actor-Critic models (see Behavioral Models section in Methods for details on how these variables are computed). The cell values and color represent the mean ± s.e.m. pearson correlation coefficient (*r*) across sessions. **c**, Confusion matrix from model recovery analysis characterizing the distinguishability of the Q-learning, REINFORCE, and Actor-Critic models on simulated data. The y-axis indicates which model was used to simulate the choice data, and the x-axis indicates which model was then fit to recover it. Cell values and color represent the mean ± s.e.m. log-likelihood per choice of the fitting model across simulated sessions. This analysis diagnoses whether the models are separable under our task design and parameter ranges. It is not a comparison of model fit to the observed data.Confusion matrix characterizing the distinguishability of the Q-learning, REINFORCE, and Actor-Critic models. The cell values and color represent the mean ± s.e.m. log-likelihood per choice of the recovered fit across sessions. The y-axis indicates which model was used to simulate the choice data, and the x-axis indicates which model was used to recover that simulated choice data. **d**, Parameter recovery for the best performing model (Q-learning (CS+ as a reward)). The x-axis represents the Q-learning model’s fitted parameters from the animals’ choice data, which was then used to generate new simulated choice data on-policy. The y-axis represents the estimates of those same parameters from fitting the Q-learning model to the simulated choice data. Each point represents one simulated session. e-h, Distributions of the best-fit parameters for models 2-5. Violin plots illustrate the distribution of the mean of each parameter across sessions; the limits of the violin represent the extreme of the fit parameters, and the boxplot within the violin marking the median (white) and interquartile range (IQR; thick black line). n = 41 sessions from 8 animals for all panels.

**Extended Data Figure 3.**
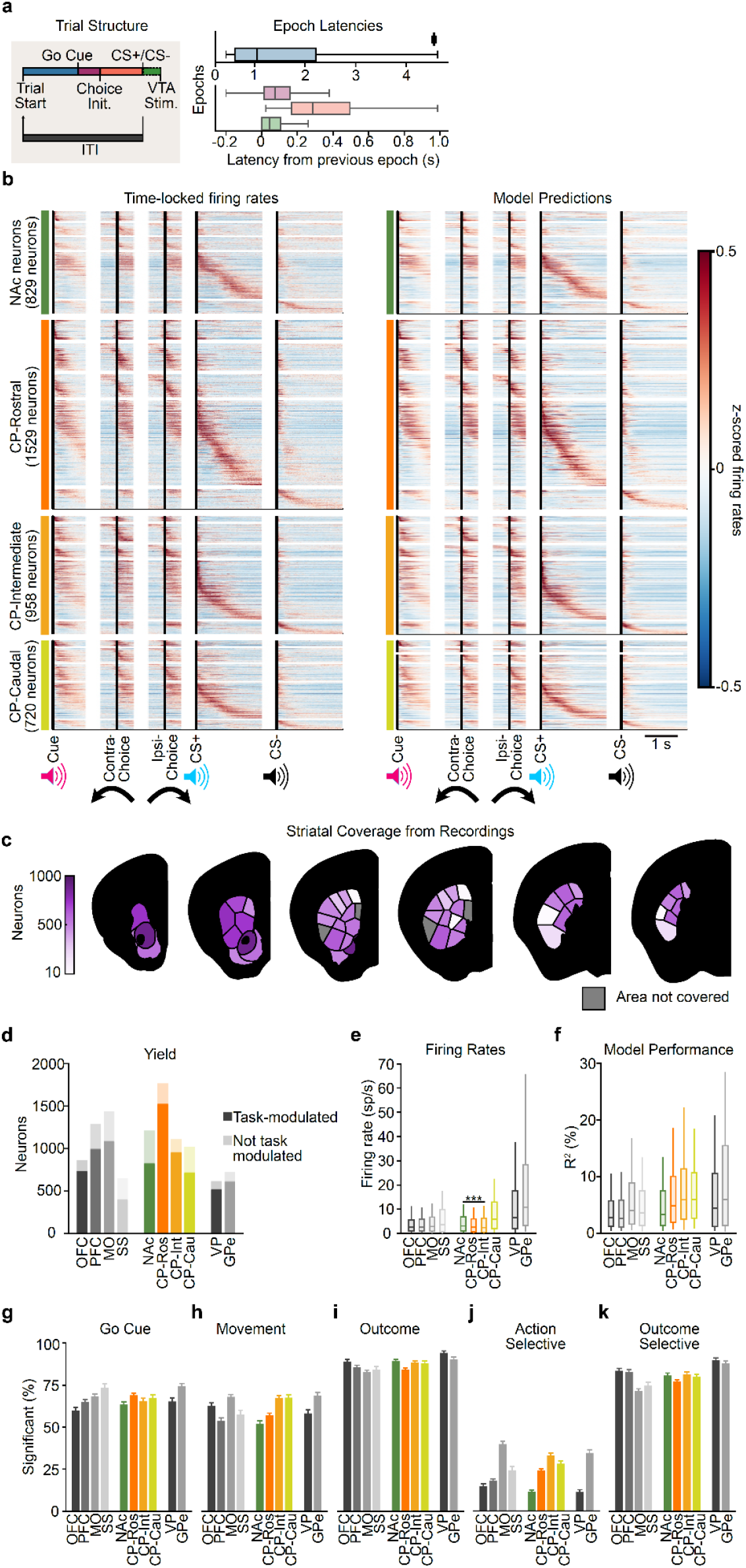
Recording statistics and modulation by task events. **a**, Box plot showing the distribution of latencies for between the main task events. Inter-trial-interval (ITI) from CS+/CS– onset to next trial start (Black), Trial Start to Go Cue (Blue), Go Cue to Choice Initiation / First Movement (Purple), Choice Initiation to CS+/CS– (Red), CS+ to Lick/First VTA stim pulse (Green). The boxplots show the median, inter-quartile range (IQR), and whiskers extending to the furthest point within 1.5x IQR (n = 22,365 trials from 41 sessions, 8 animals). **b**, *Left:* Time-locked, z-scored firing rate peri-event time histograms (PETHs) for all task-modulated single units from each striatal subregion. *Right:* Corresponding encoding model predictions for each neuron. In both plots, activity is aligned to task events: go cue, action (contra- or ipsilateral movement onset), and outcome (CS+ or CS– onset). Neurons within each subregion were first sorted based on their peak response timing in the concatenated PETHs using only odd trials, and then only even trials were used to generate the PETHs shown in the figure. The model predictions are shown in the same order as the neurons. **c**, Yield of task-modulated single units for striatal domains in the ^91,92^ atlas. Grey areas were not covered by the recordings. **d**, Yield of single-units recorded in cortical, striatal and pallidal subregions, categorized by whether each neuron was task-modulated (dark bars) or not (light extension to the barsbars). For all encoding model analyses, only task-modulated single units were included. **e**, Box plot showing firing rates across regions. **f**, Box plot showing the bilinear encoding model’s model performance, quantified as the cross-validated percentage of variance explained (R^2^) for neurons in each subregion. Box plots represent the 10th, 25th, 50th, 75th, and 90th percentiles. **g–k**, We classified neurons as significantly encoding task events by comparing the performance of a “full” encoding model with kernels for all events present to reduced models with a subset of event kernels removed (see Methods for details). We used this strategy to assess significance for (**g**) the go cue, by removing only the go cue kernel in the reduced model; (**h**) action, by removing both the left and right movement onset kernels; (**i**) outcome, by removing the CS+, laser stimulation, and CS– kernels; (**j**) choice-selectivity, by replacing the left and right movement onset kernels with a single movement onset kernel that was not choice-specific; and (**k**) outcome-selectivity, by replacing the CS+ and CS– kernels with a single CS kernel that was not reward-specific and also removing the laser stimulation onset kernel. All values were computed relative to a null distribution (see Methods for details). Error bars in bar plots represent the mean ± s.e.m.

**Extended Data Figure 4.**
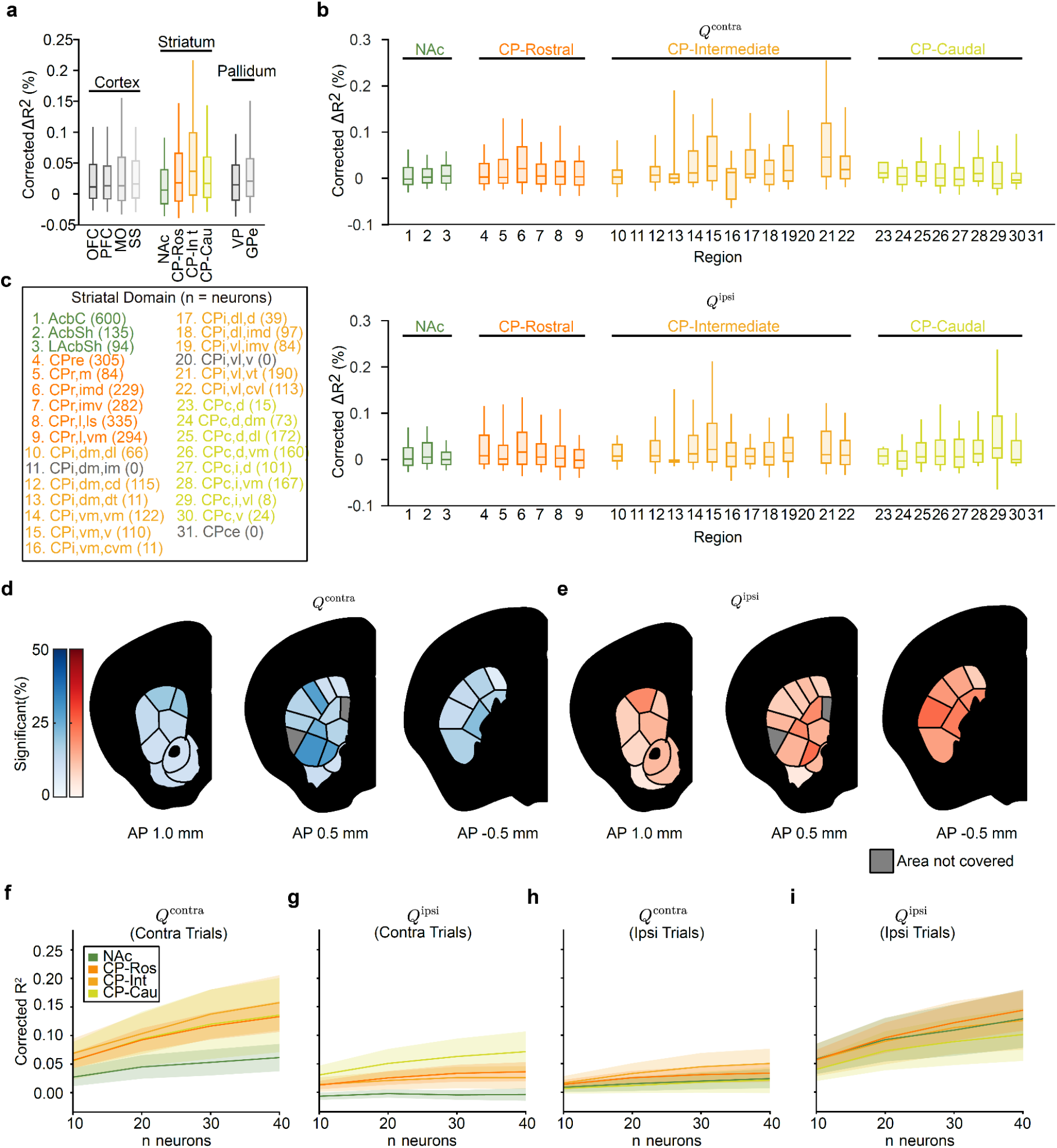
Supplementary information related to encoding and decoding analyses of *Q*^*contra*^ and *Q*^*ipsi*^. Box-plot showing the increase in null corrected variance explained (ΔR^2^) for adding the action value gains (*Q* ^*contra*^ or *Q* ^*ipsi*^) to the encoding model across cortical, striatal and pallidal regions. Corrected ΔR^2^ values are computed by subtracting the mean ΔR^2^ of a session-specific null distribution from the ΔR^2^ of the encoding model applied to the observed behavioral data (see Methods for details). **b,c**, Distribution of corrected ΔR^2^ values for (**b**)*Q* ^*contra*^ or (**c**) *Q* ^*ipsi*^ action value encoding in each striatal domain, grouped by subregion. Box plots represent the 10th, 25th, 50th, 75th, and 90th percentiles. Legend displays relation between numbers in x-axis and striatal domain names. **d,e** Coronal sections with color saturation indicating the percentage of neurons significantly encoding (**d**) *Q* ^*contra*^ or (**e**) *Q* ^*ipsi*^ Qlpsi action values in each striatal domain (*n* = 829 NAc, 1529 DS-Ros, 958 DS-Int, 720 DS-Cau). Grey areas were not covered by our recordings. **f–i**, Population decoding performance (corrected R^2^) 200 ms after the go cue,as a function of neuron count and region for (**f,g**) *Q* ^*contra*^ or (**h,i**) *Q* ^*ipsi*^ action values in trials with contralateral (**f,h**) or ipsilateral (**g,i**) choices. Lines and shading represent the mean ± s.e.m. across sessions with at least 40 neurons in that region (*n* = 10 NAc, 17 CP-Ros, 13 CP-Int, and 9 CP-Cau sessions).

**Extended Data Figure 5:**
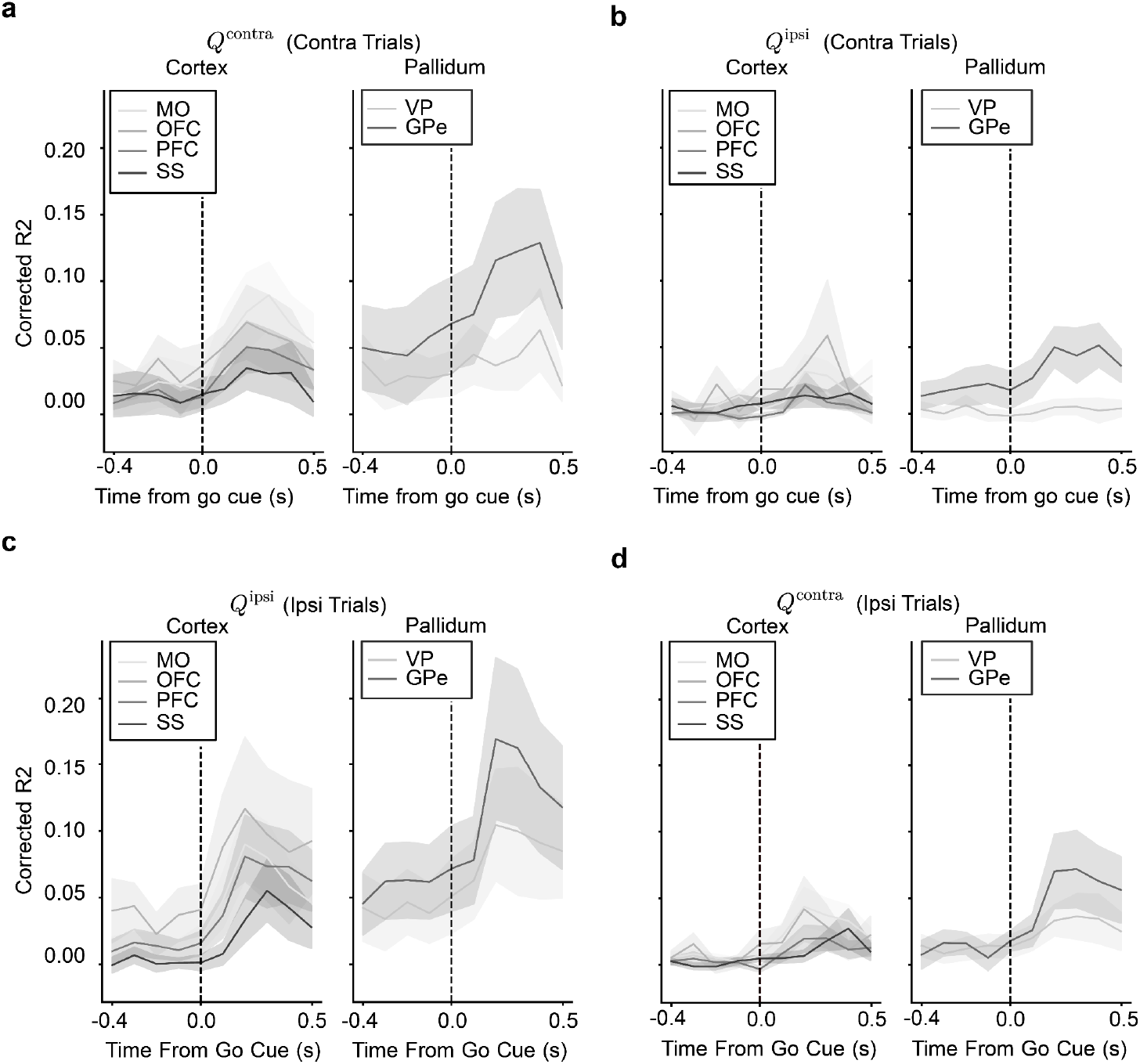
Action value decoding from striatal inputs (cortex) and outputs (pallidum). Time-resolved population decoding (groups of 20 randomly picked groups combos of 20 neurons) performance (corrected R^2^) for (**a,d**) *Q* ^*contra*^ or (**b,c**) *Q* ^*ipsi*^ action values in trials with (**a,b**) contralateral or (**c,d**) ipsilateral trials choices. The corrected R^2^ is computed by subtracting the mean R^2^ of a session-specific null distribution from the mean R^2^ of decoders applied to the observed behavioral data (see Methods for details). Lines and shading represent the mean ± s.e.m. across sessions (*n* = 19 MO, 10 OFC, 11 PFC, 12 SS, 11 GPe, and 13 VP sessions).

**Extended Data Figure 6.**
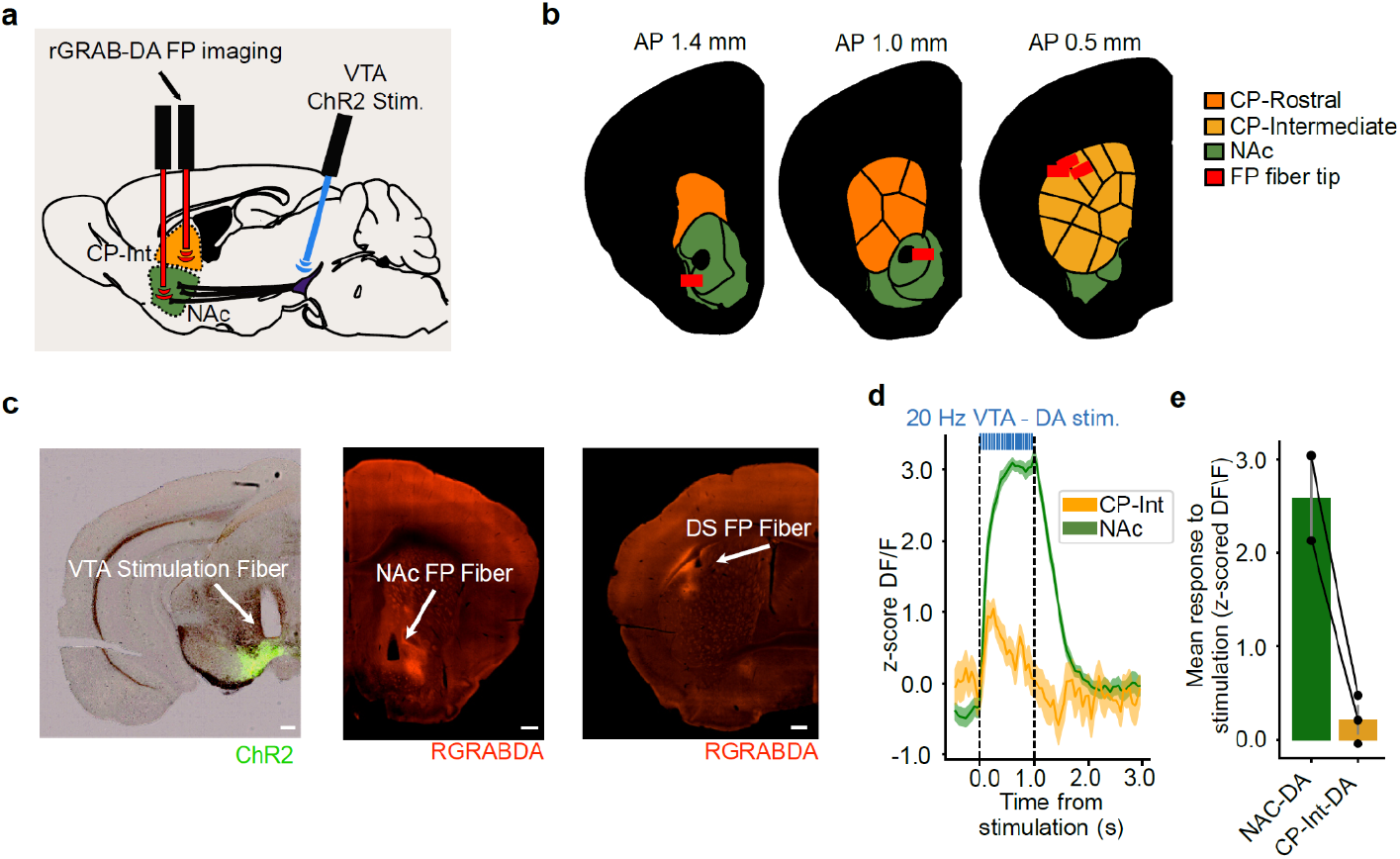
VTA stimulation primarily drives DA release in the NAc. **a**, Schematic of the fiber photometry experiment for the validation of the VTA DA stimulation approach. **b**, Histological reconstruction of photometry fiber tip locations (red rectangles) projected to the nearest 400-µm slice on the Allen CCF with striatal domains segmented ^91,92^. **c**, Representative images confirming optical fiber placement and virus expression. *Left:* Optogenetics fiber track for VTA DA stimulation and ChR2 expression (green) in the VTA. *Center:* Photometry fiber track and rGRAB-DA expression in the NAc. *Right:* Photometry fiber track and rGRAB-DA expression in the intermediate CP. Scale bars, 200 µm. **d**, Example peri-stimulus time histogram, from one animal, of DA transients in the NAc (green) and intermediate CP (orange) evoked by VTA DA stimulation (447 nm; 20 Hz; 5 ms pulse width; 8 mW). The signal represents the z-scored change in the ΔF/F fluorescence trace. Lines and shading represent the mean ± s.e.m. across 20 trials. The dashed line represents the VTA DA stimulation period. **e**, Quantification of the mean stimulation-evoked DA response in the NAc and intermediate CP during the 1 s VTA DA stimulation period (epoch represented with vertical dashed lines in (**d**)). Points indicate individual animals (n = 2 NAc animals, 3 intermediate CP animals), and lines connect data from the same animal. Error bars represent the mean ± s.e.m. across animals.

**Extended Data Figure 7:**
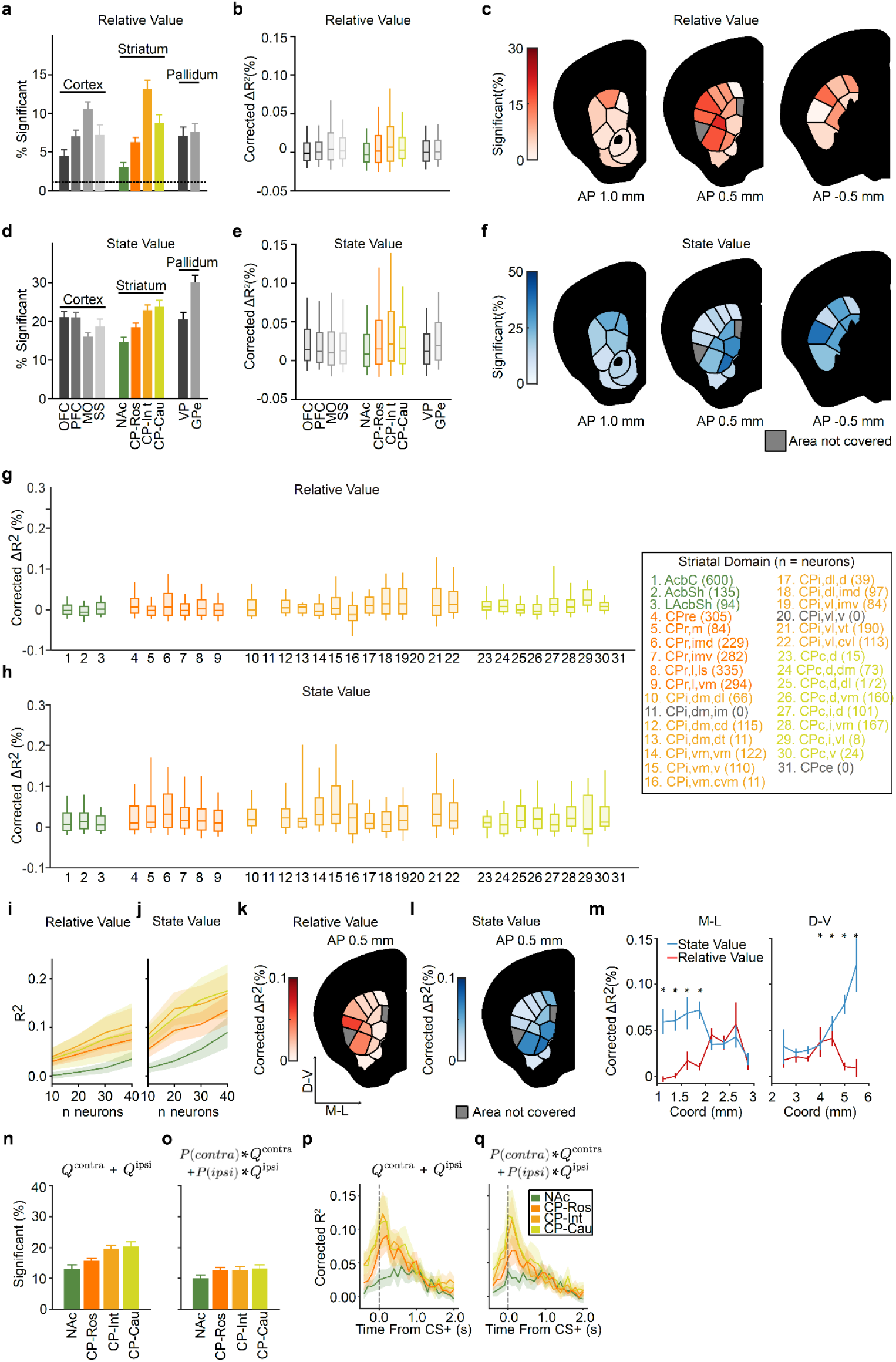
Supplementary information on state and relative value representations in striatum. **a**, Percentage of neurons significantly encoding relative value, **b**, and mean null corrected explained variance increase (ΔR^2^) for adding the relative value gain to the encoding model across cortical, striatal and pallidal regions. Bar plots show mean ± s.e.m. Box plots represent the 10th, 25th, 50th, 75th, and 90th percentiles (*n* = 829 NAc, 1529 DS-Ros, 958 DS-Int, 720 DS-Cau). Corrected ΔR^2^ are computed by subtracting the mean ΔR^2^ of a session-specific null distribution from the ΔR^2^ of the encoding-model applied to the observed behavioral data (see Methods for details). **c**, Coronal sections with color saturation indicating the percentage of neurons significantly encoding relative value in each striatal domain. Grey areas were not sufficiently covered by the recordings. **d,e,f**, Same as (**a-c**) but for state value. **g,h**, Distribution of ΔR^2^ values for (**g**) relative or (**h**) state value encoding in each striatal domain, grouped by striatal subregion. Box plots represent the 10th, 25th, 50th, 75th, and 90th percentiles. Legend displays relation between numbers in x-axis and striatal domain names. **i, j**, Population (20 neurons) decoding performance (R^2^) for (**i**) relative and (**j**) state value as a function of neuron count and striatal subregion. Line and shading represent mean ± s.e.m. across sessions with at least 40 neurons in that region (*n* = 10 NAc, 17 CP-Ros, 13 CP-Int, and 9 CP-Cau sessions). **k, l**, Coronal sections illustrating analyzed striatal domains within CP-Int, the subregion where we found the strongest value encoding. Saturation indicates the local corrected ΔR^2^ from adding (**k**) relative or (**l**) state value gains. Grey areas were not sufficiently covered by the recordings. **m**, Average corrected ΔR^2^ from adding relative value (red) and state value (blue) gains to the encoding model along the mediolateral (ML; left), and dorsoventral (DV; right) axes within intermediate CP. Line and error bar represent mean ± binomial s.e.m (*P < 0.05, Mann–Whitney U test). **n,o**, Percentage of neurons in each striatal subregion significantly encoding state value formulated as the (**n**) unweighted or (**o**) weighted sum of q-values. Bar plots represent mean ± s.e.m. (n = 829 NAc, 1529 DS-Ros, 958 DS-Int, 720 DS-Cau). **p,q**, Time-resolved population decoding performance (corrected R^2^) for state value (formulated as the (**p**) unweighted or (**q**) weighted sum of q-values) in trials with contralateral choices (based on groups of 20 randomly chosen neurons). The corrected R^2^ is computed by subtracting the mean R^2^ of a session-specific null distribution from the mean R^2^ of decoders applied to the observed behavioral data (see Methods for details). Lines and shading represent the mean ± s.e.m. across sessions (n = 18 NAc, 20 CP-Ros, 15 CP-Int, and 13 CP-Cau sessions).

**Extended Data Figure 8.**
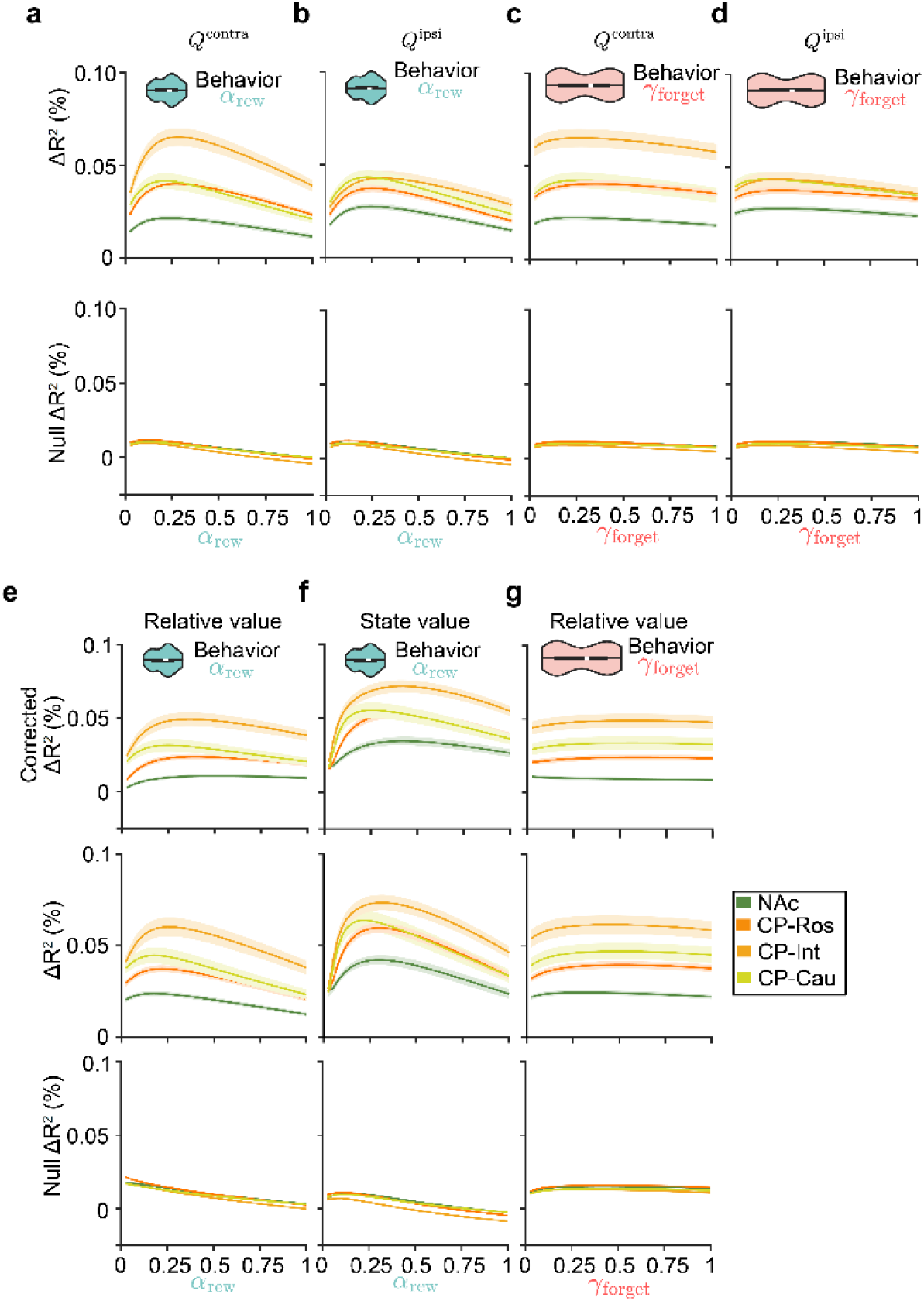
Across learning and forgetting rates, relative and state value coding is strongest in intermediate CP. **a,b**, Improvement in uncorrected explained variance (ΔR^2^) in the encoding model as a function of the learning rate parameter (*α* _*rew*_) used to generate the (**a**) *Q* ^*contra*^ and *Q* ^*ipsi*^ (**b**) values inputted into the model for the real session (top) or null distribution of pseudosessions (bottom). Colored lines indicate different striatal subregions.**c,d**, Same as (**a,b**) but when varying the forgetting parameter (*γ* _*forget*_) used to generate the (**c**) *Q* ^*contra*^ and *Q* ^*ipsi*^ (**d**) values instead of *α* _*rew*_. **e-g**, Top: Improvement in corrected explained variance (ΔR^2^) as a function of the learning rate parameter (**e,f**, αRew) or forgetting parameter (g, *γ* _*forget*_) used to generate the (**e,g**) relative, and (**f**) state values inputted into the model. No *γ* _*forget*_ equivalent of (**f**) is plotted since the state value update rule does not contain a forgetting term. Middle: Same as top row but for uncorrected ΔR^2^. Bottom: Same as middle row but for the pseudosession-based null distribution. Corrected ΔR^2^ (top row) is computed by subtracting the mean ΔR^2^ of the pseudosession-based null distribution (bottom row) from the ΔR^2^ of the encoding model applied to the observed behavioral data (middle row; see Methods for details). For all plots lines and shading represent mean ± s.e.m. across sessions. Violin plots illustrate the distribution of *α* _*rew*_ or *γ* _*forget*_ obtained from fitting the behavior. Plot limits are the limits of the observed data, the boxplot within the violin marks the median (white) and IQR (thick black line).

**Extended Data Figure 9.**
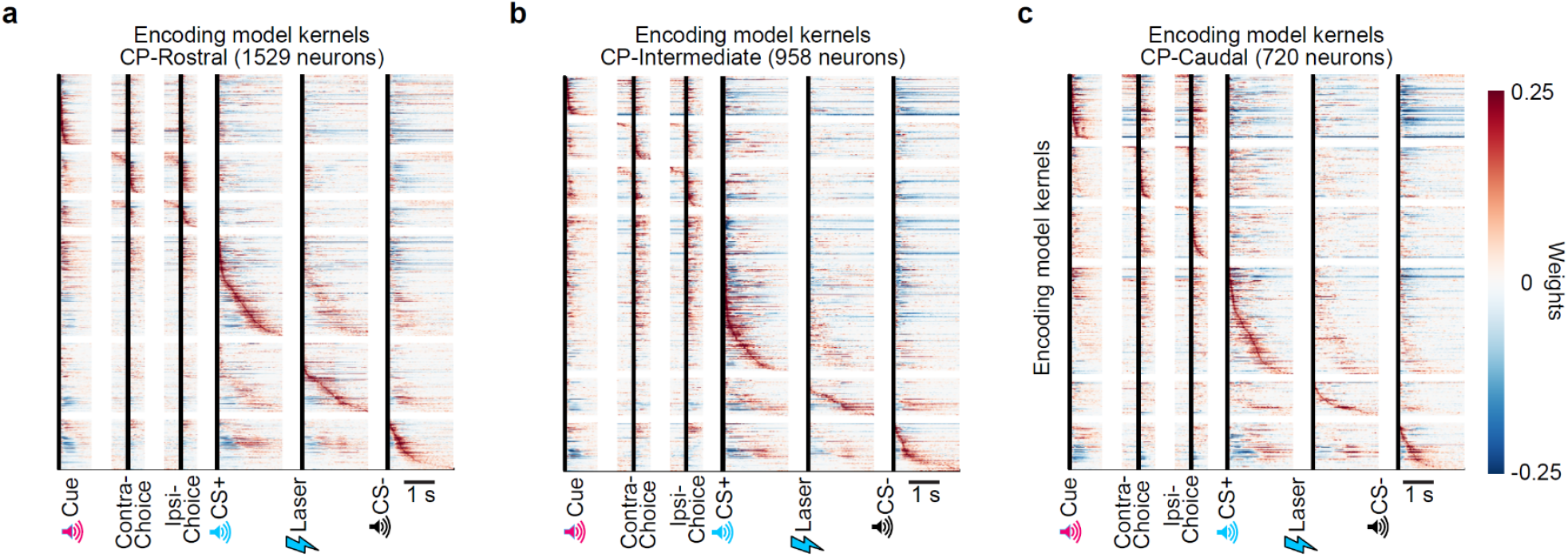
Outcome coding ending is prevalent across striatal subregions. **a-c**, Time-locked normalized kernels from the encoding model for individual neurons in (**a**) CP-Rostral, (**b**) CP-Intermediate, (**c**) CP-Caudal. Neurons are sorted by the peak across their kernels.

**Extended Data Figure 10.**
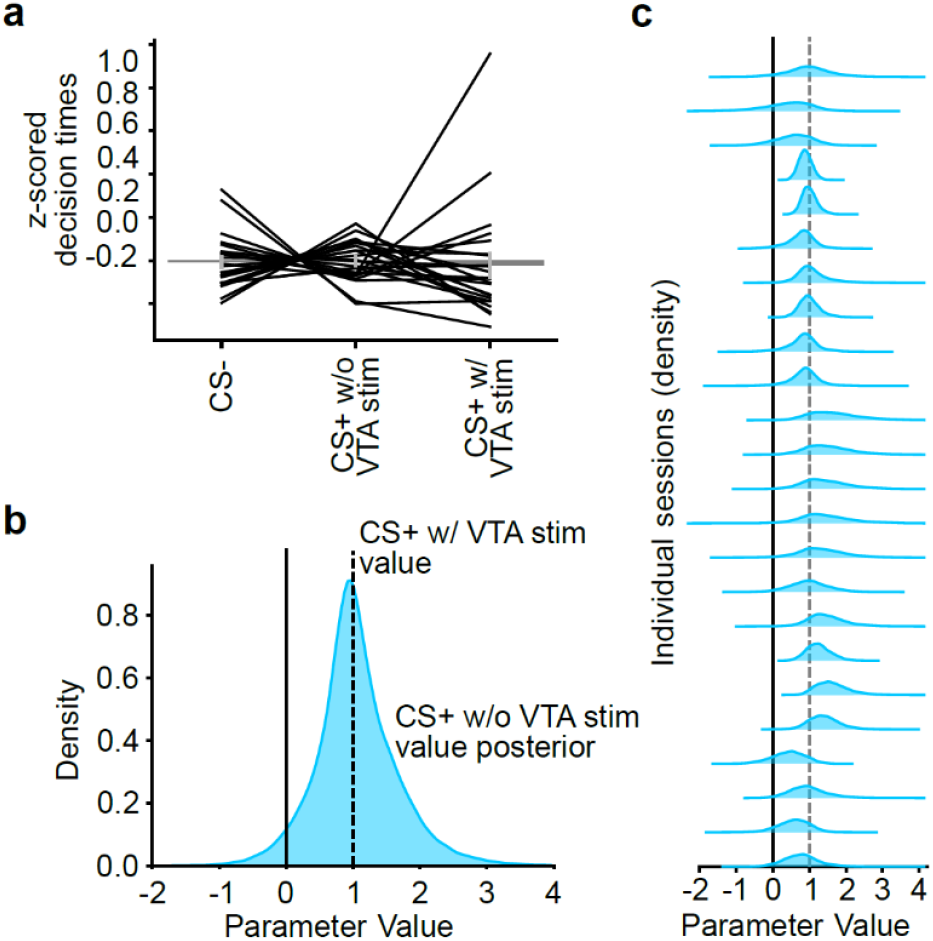
CS+ fitted value is significantly greater than 0, and similar to CS+ w/ VTA stimulation value. **a**, Average normalized decision times (Time from Go Cue to decision) after trials with different outcomes. Individual lines represent sessions. Bar plots show mean ± s.e.m. **b**, Posterior distribution of the CS+ w/o laser parameter (comparison to 1 which is the pre-set value for the CS+ with laser that is then scaled by *β _rew_*, See Equations 24 and 25 in Methods for details) across sessions. **c**, Same as (**b**) but for individual sessions. For all panels we included all sessions with at least 5 trials in each outcome condition (*n* = 24 sessions from 7 mice), so that we were able to estimate the value of the CS+ w/o VTA stim.

**Extended Data Figure 11.**
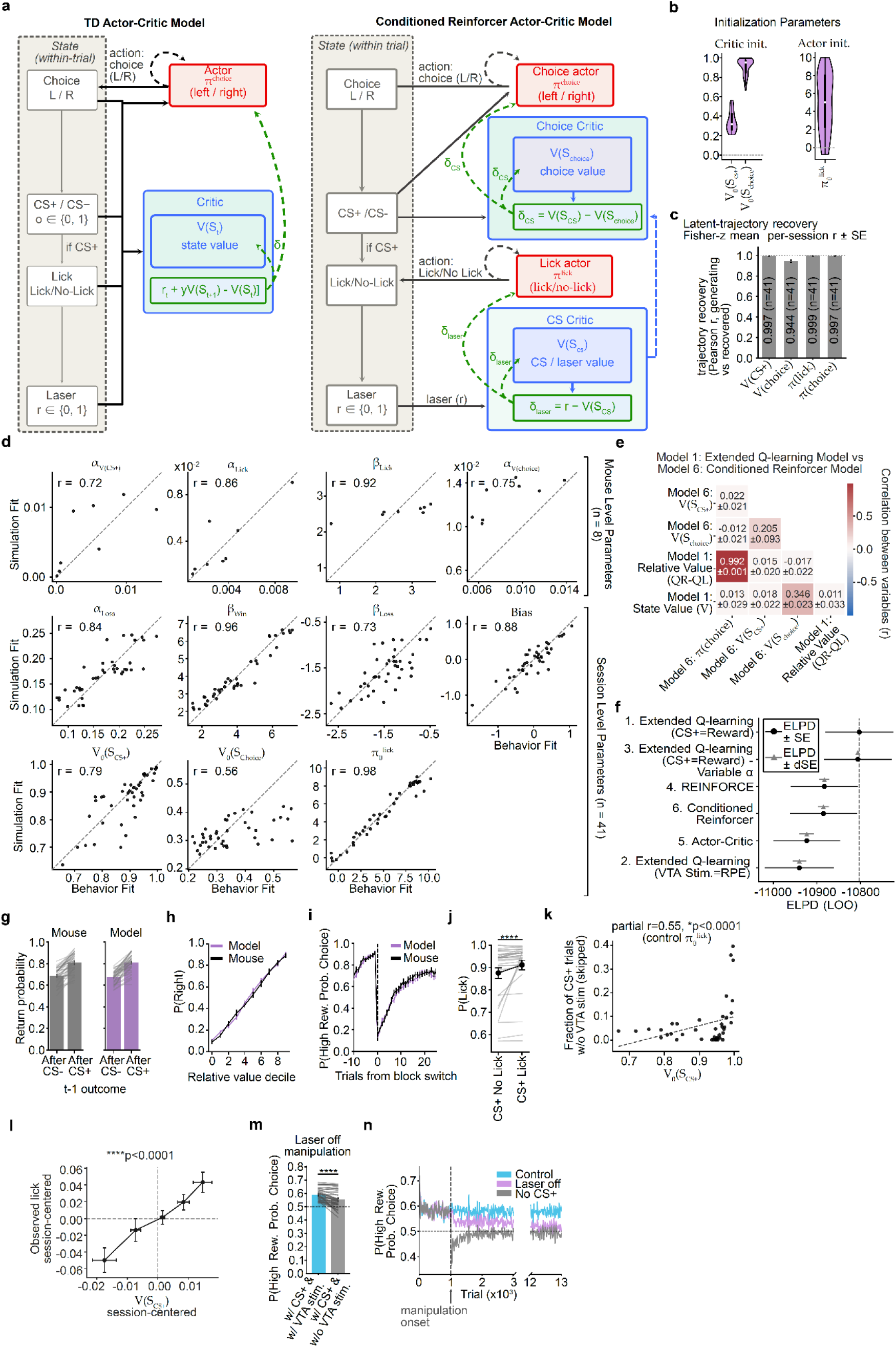
The Conditioned Reinforcer model is recoverable and reproduces the mice’s behavior. **a**, Left, Schematic of the temporal-difference (TD) Actor-Critic model. Within a trial the animal transitions through the choice state, the outcome state signalled by the CS+ or CS– (*o* ∈ {0,1}), the lick / no-lick state on CS+ trials, and the laser state (*r* ∈{0,1}). A single critic (blue) learns the state value *V*(*S*_t_) and the TD error *r*_*t*_*+ϒ V*(*S*_t+1_) − *V*(*S*_*t*_) trains the choice actor *π* ^*choice*^(red). Right, Conditioned Reinforcer Model from Fig. 6a. **b**, Distributions of the fitted initialization parameters for the two critics (*V*_0_(*S*_*cs*+_),V_0_(*S*_*choice*_) and the lick actor (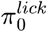). The boxplot within the violin marks the median (white) and IQR (thick black line). **c**, Recovery of the model’s latent variable trajectories. Bars show the per-session Pearson correlation between the generating and the recovered trajectory of each latent variable, averaged in Fisher-z space (mean ± s.e.). **d**, Parameter recovery for the Conditioned Reinforcer model. The x-axis represents the model’s fitted parameters from the animals’ choice data, which was then used to generate new simulated choice data on-policy. The y-axis represents the estimates of those same parameters from fitting the model to the simulated choice data. Top, parameters fit per mouse; middle and bottom, parameters fit per session.. **e**, Similarity between the different latent decision variables of the extended Q-learning model (Model 1) and of the Conditioned Reinforcer model (Model 6). The cell values and color represent the mean ± s.e.m. pearson correlation coefficient (r) across sessions. **f**, Model comparison by PSIS-LOO with the Conditioned Reinforcer model added to the comparisons in Extended Data Fig. 2a. Models are ranked by expected log pointwise predictive density (ELPD), for which greater values indicate superior out-of-sample predictive performance. Both markers show each model’s ELPD; only the error bars differ. Circles: ± s.e. of the ELPD. Triangles: ± s.e. of the trial-wise predictive difference from the best model (dSE). All models predicted held-out choices within 0.5% of one another (cross-validated mean predicted probability of the observed choice, 67.8–68.3%). N = 22,406 trials, n = 8 mice and n = 41 sessions. **g**, Calibration of the Conditioned Reinforcer model’s win-stay behavior. Bar plot showing the probability of repeating a choice following an CS+ or CS– trial for the mice (grey) and for the model’s on-policy simulations (purple). Error bars represent the mean ± s.e.m. across sessions. **h**, Calibration of the Conditioned Reinforcer model’s action selection policy. The probability of choosing the right action is plotted as a function of the percentile of the model derived choice propensities (*π* ^*choice*^) for the mice (grey) and for the model’s on-policy simulations (purple). Lines and shading represent the mean ± s.e.m. across sessions. **i**, Calibration of the Conditioned Reinforcer model’s reward-contingency reversal (block switch) behavior. The probability of choosing the high reward probability action is plotted as a function of trials from block switches for the mice (grey) and for the Conditioned Reinforcer model’s on-policy simulations (purple). Lines and error bars represent the mean ± s.e.m. across sessions. **j**, Probability of licking on CS+ trials with and without a lick, in the model’s on-policy simulations. Individual lines represent sessions (***P < 0.001). **k**, Correlation across sessions between the fitted *V*_0_(*S*_*cs*+_), (V (*S*_*CS+*_) session initialization) and the fraction of CS+ trials without VTA DA stimulation, controlling for 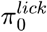(π ^*lick*^ session initialization). Dashed line represents linear regression. **l**, Relationship between the model’s *V*(*S*_*cs*+_)and the observed probability of licking both centered within session, for sessions where there were at least 5 CS+ trials without VTA DA stimulation *n* = 24 sessions from 7 mice. Points and error bars represent mean ± s.e.m. across sessions. ****p<0.0001 from fixed effect in logistic mixed effects model. **m**, Model performance when the VTA stim. is removed *in silico*. Individual lines represent sessions and the dashed line indicates chance level (0.5) (\*\*\**P* < 0.001). **n**, Model performance as a function of trial number for the control (cyan), laser-off (pink) and no-CS+ (grey) simulated manipulations. The dashed line indicates the onset of the manipulation. For simulations in (**g,h,i,m,n**) the parameters were obtained from model fits to the behavior of animals and sessions in Fig. 1-5f. For every panel except (l), *n* = 41 sessions from 8 animals. See Extended Data Table 7 for statistical details.

**Extended Data Figure 12.**
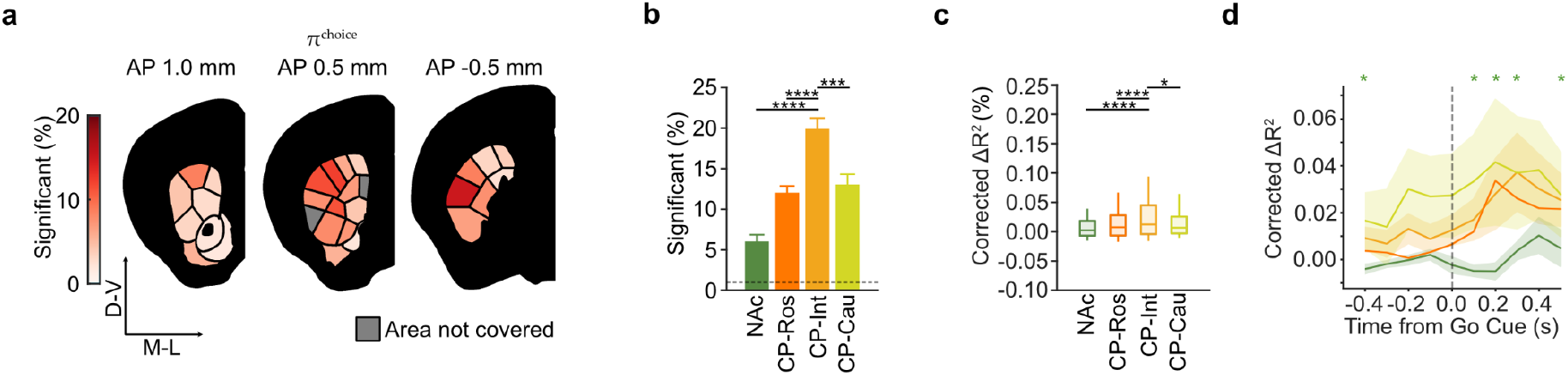
The Conditioned Reinforcer model’s choice policy is encoded most strongly in the intermediate CP. **a**, Coronal sections with color saturation indicating the percentage of neurons significantly encoding the model’s choice policy, *π* ^*choice*^, in each striatal domain. Grey areas were not covered by the recordings. **b**, Percentage of neurons significantly encoding in each striatal subregion. Bars denote the mean ± bernoulli s.e.m. across all recorded neurons and the dashed line indicates the chance level for significance (1%; see Methods for details on significance testing). **c**, Distribution of corrected ΔR^2^ values from removing the *π* ^*choice*^ gain from the encoding model, computed by subtracting the mean ΔR^2^ of a pseudosession-based null distribution from the ΔR^2^ of the encoding model applied to the observed behavioral data (see Methods for details).Box plots represent the 10th, 25th, 50th, 75th, and 90th percentiles. n = 829 NAc, 1529 CP-Ros, 958 CP-Int, 720 CP-Cau task-modulated single units. *P < 0.05, **P < 0.01, ***P < 0.001, ****P < 0.0001 from χ^2^ tests highlighted in the results, for all comparison see Extended Data Table 7. **d**, Time-resolved population decoding performance (corrected R2) for *π* ^*choice*^ in contralateral choice trials (based on groups of 40 randomly chosen neurons). The corrected R^2^ is computed by subtracting the mean R^2^ of a session-specific null distribution from the mean R^2^ of decoders applied to the observed behavioral data (see Methods for details). Lines and shading represent the mean ± s.e.m. across sessions (*n* = 16 NAc, 19 CP-Ros, 15 CP-Int, and 12 CP-Cau sessions). * indicates time points where decoding performance for CP was significantly different from NAc (*P* ≤ 0.05, from two-tailed Mann–Whitney U test at each timepoint).

## Extended Data Tables

**Extended Data Table 1.**
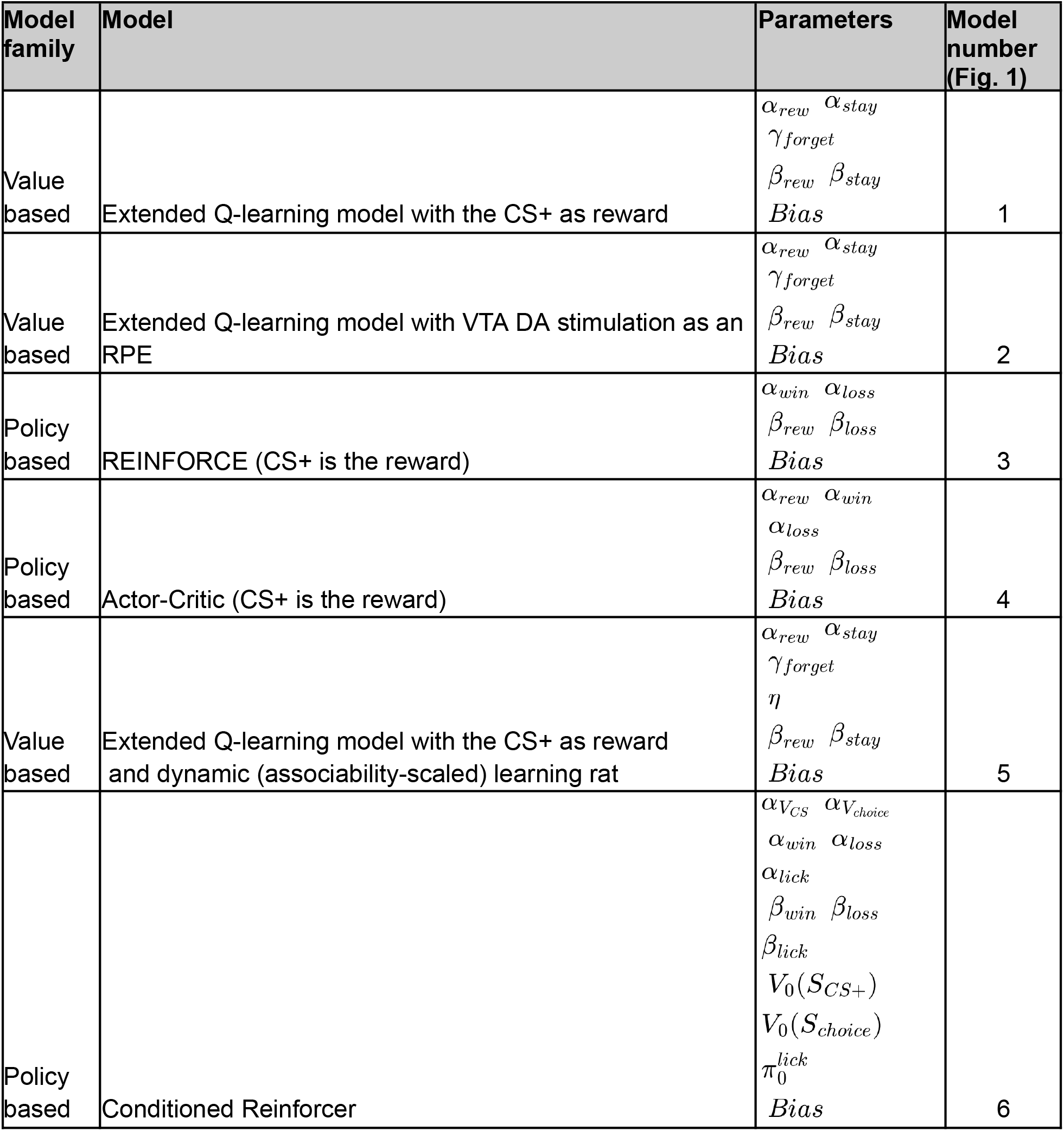
Summary of behavioral models tested.

| Model family | Model | Parameters | Model number (Fig. 1) |
| --- | --- | --- | --- |
| Value based | Extended Q-learning model with the CS+ as reward | $\alpha_{rew}$ $\alpha_{stay}$<br>$\gamma_{forget}$<br>$\beta_{rew}$ $\beta_{stay}$<br>$Bias$ | 1 |
| Value based | Extended Q-learning model with VTA DA stimulation as an RPE | $\alpha_{rew}$ $\alpha_{stay}$<br>$\gamma_{forget}$<br>$\beta_{rew}$ $\beta_{stay}$<br>$Bias$ | 2 |
| Policy based | REINFORCE (CS+ is the reward) | $\alpha_{win}$ $\alpha_{loss}$<br>$\beta_{rew}$ $\beta_{loss}$<br>$Bias$ | 3 |
| Policy based | Actor-Critic (CS+ is the reward) | $\alpha_{rew}$ $\alpha_{win}$<br>$\alpha_{loss}$<br>$\beta_{rew}$ $\beta_{loss}$<br>$Bias$ | 4 |
| Value based | Extended Q-learning model with the CS+ as reward and dynamic (associability-scaled) learning rat | $\alpha_{rew}$ $\alpha_{stay}$<br>$\gamma_{forget}$<br>$\eta$<br>$\beta_{rew}$ $\beta_{stay}$<br>$Bias$ | 5 |
| Policy based | Conditioned Reinforcer | $\alpha_{V_{CS}}$ $\alpha_{V_{choice}}$<br>$\alpha_{win}$ $\alpha_{loss}$<br>$\alpha_{lick}$<br>$\beta_{win}$ $\beta_{loss}$<br>$\beta_{lick}$<br>$V_0(S_{CS+})$<br>$V_0(S_{choice})$<br>$\pi_0^{lick}$<br>$Bias$ | 6 |

**Extended Data Table 2.** Summary of priors used for each parameter.

| Parameter | Model(s) | Constraints | Population-level prior | $M, \tau$ | Transformation |
| --- | --- | --- | --- | --- | --- |
| $\alpha_{rew}, \alpha_{stay}, \gamma_{forget}, \alpha_{win}, \alpha_{loss}$ | 1-5 | $\alpha, \gamma \in (0, 1)$ | $\alpha \sim N(M, \tau)$<br>$\gamma \sim N(M, \tau)$ | $M = 0.5$<br>$\tau = 1$ | $\alpha = \text{logit}^{-1}(\tilde{\alpha})$<br>$\gamma = \text{logit}^{-1}(\tilde{\gamma})$ |
| $\eta$ | 5 | $\eta \in (0, 1)$ | $\tilde{\eta} \sim N(M, \tau)$ | $M = -2$<br>$\tau = 1.5$ | $\eta = \text{logit}^{-1}(\tilde{\eta})$ |
| $\alpha_{VCS}, \alpha_{Vchoice}, \alpha_{win}, \alpha_{lick}$ | 6 | $\alpha \in (0, 1)$ | $\tilde{\alpha} \sim N(M, \tau)$ | $M = -3.0$<br>$\tau = 0.5$ | $\alpha = \text{logit}^{-1}(\tilde{\alpha})$ |
| $\alpha_{loss}$ | 6 | $\alpha \in (0, 1)$ | $\tilde{\alpha} \sim N(M, \tau)$ | $M = -2.0$<br>$\tau = 0.5$ | $\alpha = \text{logit}^{-1}(\tilde{\alpha})$ |
| $\beta_{rew}, \beta_{stay}, \beta_{win}, \beta_{loss}$ | 1-5 | $\beta \in \mathbb{R}$ | $\beta \sim N(M, \tau)$ | $M = 0$<br>$\tau = 1$ | –NA |
| $\beta_{win}, \beta_{loss}, \beta_{lick}$ | 6 | $\beta \in \mathbb{R}$ | $\beta \sim N(M, \tau)$ | $M = 0$<br>$\tau = 2$ | –NA |
| $V_0(S_{CS+}), V_0(S_{choice})$ | 6 | $V \in (0, 1)$ | $\tilde{V} \sim N(M, \tau)$ | $M = 0$<br>$\tau = 1.5$ | $V = \text{logit}^{-1}(\tilde{V})$ |
| $\pi_0^{lick}$ | 6 | $\pi \in \mathbb{R}$ | $\pi \sim N(M, \tau)$ | $M = 0$<br>$\tau = 1$ | –NA |
| $Bias$ | 1-6 | $Bias \in \mathbb{R}$ | $Bias \sim N(M, \tau)$ | $M = 0$<br>$\tau = 1$ | –NA |
| All standard deviations | 1-4 | $\sigma \in (0, \infty)$ | $\sigma \sim N(M, \tau)$ | $M = 0$<br>$\tau = 1$ | $\sigma = \exp(\tilde{\sigma})$ |
| All standard deviations | 5 | $\sigma \in (0, \infty)$ | $\sigma \sim N(M, \tau)$ | $M = 0$<br>$\tau = 0.5 \text{ to } 1$ | $\sigma = \exp(\tilde{\sigma})$ |
| All standard deviations | 6 | $\sigma \in (0, \infty)$ | $\sigma \sim N(M, \tau)$ | $M = 0$<br>$\tau = 0.3 \text{ to } 1$ | $\sigma = \exp(\tilde{\sigma})$ |

∼ denotes the unconstrained variable Stan samples; the Transformation column maps it to the modeled parameter. A prior on the plain parameter ( $\alpha \sim N$ ) is specified on the natural scale ( $M$ is a rate); a prior on the tilde variable ( $\tilde{\alpha} \sim N$ ) is on the unconstrained scale ( $M$ is a log-odds).

**Extended Data Table 3.** Group labels for recorded brain subregions.

| Allen Brain Atlas | Chon et al. 2019 Labels | Decoders Groupings | Projection Analysis Groupings |
| --- | --- | --- | --- |
| SSp-bfd5 |  | SS | SS |
| SSp-bfd6a |  | SS | SS |
| SSp-bfd6b |  | SS | SS |
| SSp-n6a |  | SS | SS |
| SSp-n6b |  | SS | SS |
| SSp-un2/3 |  | SS | SS |
| SSp-un4 |  | SS | SS |
| SSp-un5 |  | SS | SS |
| SSp-un6a |  | SS | SS |
| SSp-un6b |  | SS | SS |
| SSp-bfd1 |  | SS | SS |
| SSp-bfd2/3 |  | SS | SS |
| SSp-bfd4 |  | SS | SS |
| SSp-ll5 |  | SS | SS |
| SSp-ll6a |  | SS | SS |
| SSp-ll6b |  | SS | SS |
| SSp-m4 |  | SS | SS |
| SSp-m5 |  | SS | SS |
| SSp-m6a |  | SS | SS |
| SSp-m6b |  | SS | SS |
| SSp-n4 |  | SS | SS |
| SSp-n5 |  | SS | SS |
| SSp-ul2/3 |  | SS | SS |
| SSp-ul4 |  | SS | SS |
| SSp-ul5 |  | SS | SS |
| SSp-ul6a |  | SS | SS |
| SSp-un6b |  | SS | SS |
| SSp-ul6b |  | SS | SS |
| SSp-n2/3 |  | SS | SS |
| MOp1 |  | MO | MO |
| MOp2/3 |  | MO | MO |
| MOp5 |  | MO | MO |
| MOp6a |  | MO | MO |
| MOp6b |  | MO | MO |
| MOs2/3 |  | MO | MO |
| MOs5 |  | MO | MO |
| MOs6a |  | MO | MO |
| MOs6b |  | MO | MO |
| MOs1 |  | MO | MO |
| DP |  | PFC | PFC |
| ILA1 |  | PFC | PFC |
| ILA2/3 |  | PFC | PFC |
| ILA5 |  | PFC | PFC |
| ILA6a |  | PFC | PFC |
| PL1 |  | PFC | PFC |
| PL2/3 |  | PFC | PFC |
| PL5 |  | PFC | PFC |
| PL6a |  | PFC | PFC |
| PL6b |  | PFC | PFC |
| ACAd5 |  | PFC | ACC |
| ACAd6a |  | PFC | ACC |
| ACAv2/3 |  | PFC | ACC |
| ACAv5 |  | PFC | ACC |
| ACAv6a |  | PFC | ACC |
| ACAd6b |  | PFC | ACC |
| ACAd2/3 |  | PFC | ACC |
| ORBI1 |  | OFC | OFC |
| ORBI2/3 |  | OFC | OFC |
| ORBI5 |  | OFC | OFC |
| ORBI6a |  | OFC | OFC |
| ORBI6b |  | OFC | OFC |
| ORBm2/3 |  | OFC | OFC |
| ORBm5 |  | OFC | OFC |
| ORBvl1 |  | OFC | OFC |
| ORBvl2/3 |  | OFC | OFC |
| ORBvl5 |  | OFC | OFC |
| ORBm6a |  | OFC | OFC |
| ORBvl6a |  | OFC | OFC |
| FRP5 |  | OFC | OFC |
| FRP6a |  | OFC | OFC |
| CP | Caudoputamen- rostral extreme | Rostral |  |
| CP | Caudoputamen- rostral, medial | Rostral |  |
| CP | Caudoputamen- rostral, intermediate, dorsal | Rostral |  |
| CP | Caudoputamen- rostral, intermediate, ventral | Rostral |  |
| CP | Caudoputamen- rostral, lateral, lateral strip | Rostral |  |
| CP | Caudoputamen- rostral, lateral, ventromedial | Rostral |  |
| CP | Caudoputamen- intermediate, dorsomedial, dorsolateral | Intermediate |  |
| CP | Caudoputamen- intermediate, dorsomedial, intermedial | Intermediate |  |
| CP | Caudoputamen- intermediate, dorsomedial, central dorsal | Intermediate |  |
| CP | Caudoputamen- intermediate, dorsomedial, dorsal tip | Intermediate |  |
| CP | Caudoputamen- intermediate, ventromedial, ventromedial | Intermediate |  |
| CP | Caudoputamen- intermediate, ventromedial, ventral | Intermediate |  |
| CP | Caudoputamen- intermediate, ventromedial, central ventromedial | Intermediate |  |
| CP | Caudoputamen- intermediate, dorsolateral, dorsal | Intermediate |  |
| CP | Caudoputamen- intermediate, dorsolateral, intermedial dorsal | Intermediate |  |
| CP | Caudoputamen- intermediate, ventrolateral, intermedial ventral | Intermediate |  |
| CP | Caudoputamen- intermediate, ventrolateral, ventral | Intermediate |  |
| CP | Caudoputamen- intermediate, ventrolateral, ventral tip | Intermediate |  |
| CP | Caudoputamen- intermediate, ventrolateral, central ventrolateral | Intermediate |  |
| CP | Caudoputamen- caudal, dorsal | Caudal |  |
| CP | Caudoputamen- caudal, dorsal, dorsomedial | Caudal |  |
| CP | Caudoputamen- caudal, dorsal, dorsolateral | Caudal |  |
| CP | Caudoputamen- caudal, dorsal, ventromedial | Caudal |  |
| CP | Caudoputamen- caudal, intermediate, dorsal | Caudal |  |
| CP | Caudoputamen- caudal, intermediate, ventromedial | Caudal |  |
| CP | Caudoputamen- caudal, intermediate, ventrolateral | Caudal |  |
| CP | Caudoputamen- caudal, ventral | Caudal |  |
| CP | Caudoputamen- caudal extreme | Caudal |  |
| ACB | Accumbens nucleus, core region | NAc |  |
| ACB | Accumbens nucleus, shell region | NAc |  |
| ACB | Lateral accumbens, shell region | NAc |  |
| GPe |  | GPe |  |
| MA |  | VP |  |
| SI |  | VP |  |

**Extended Data Table 4.** Summary of coordinates and locations of recording targets.

| Center of craniotomy coordinates | Probe Depth | Probe Angle | Targeted regions |
| --- | --- | --- | --- |
| 1.94 mm AP, -0.76 mm ML | -4 mm DV | 325/35° azimuth, 15° zenith | MO, PFC |
| 1.84 mm AP, -2.06 mm ML | -5.12 mm DV | 325/35° azimuth, 20° zenith | MO, CP, NAc, VP |
| 2.43 mm AP, 1.72 mm ML | -4.82 mm DV | 325/35° azimuth, 20° zenith | MO, OFC, NAc |
| 0.16 mm AP, 2.58 mm ML | -4.04 mm DV | 325/35° azimuth, 15° zenith | SS, CP, GPe |
| 0.6 mm AP, 3.87 mm ML | -5.27 mm DV | 90°/180° azimuth, 30° zenith | SS, CP, NAc, VP |
| 1.6 mm AP, 0.3 mm ML | -5.37 mm DV | 90°/180° azimuth, 20° zenith | PFC, NAc, VP |
| 0.7 mm AP, 3.0 mm ML | -5.48 mm DV | 90°/180° azimuth, 20° zenith | CP, NAc, VP |
| 0.8 mm AP, -1.78 mm ML | -4.00 mm DV | 90°/180° azimuth, 10° zenith | MO, CP |
| 2.7 mm AP, -1.19 mm ML | -4.00 mm DV | 90°/180° azimuth, 10° zenith | MO, OFC |
| 1.3 mm AP, 0.3 mm ML | 4.44 mm DV | 90°/180° azimuth, 10° zenith | PFC, CP, NAc |

**Extended Data Table 5.** Summary of Neuropixels recordings.

| Mouse Number | Mouse Name | Brain Regions Recorded | Number of recording sessions |
| --- | --- | --- | --- |
| 1 | dop_47 | SS, MOp, CP, NAc, GPe, VP | 4 |
| 2 | dop_48 | OFC, PFC, SS, MOs, CP, NAc, VP | 4 |
| 3 | dop_49 | OFC, PFC, SS, MOs, CP, NAc, VP, GPe | 8 |
| 4 | dop_50 | OFC, PFC, SS, MOs, CP, NAc, VP, GPe | 6 |
| 5 | dop_53 | SS, MOs, NAc, CP, VP, GPe | 5 |
| 6 | dop_56 | OFC, PFC, MO, CP, NAc, VP | 5 |
| 7 | dop_57 | MO, CP, NAc | 3 |
| 8 | dop_59 | PFC, MO, CP, NAc | 6 |

**Extended Data Table 6.** Summary of Allen Projection Data injections analyzed in Fig. 3l–m.

| Experiment ID | Mouse line | Injection region |
| --- | --- | --- |
| 175372863 | Wild-type | AAA |
| 146593590 | Wild-type | ACAd |
| 112458114 | Wild-type | ACAd |
| 139426984 | Wild-type | ACAd |
| 166323896 | A930038C07Rik-Tg1-Cre | ACAd |
| 286774770 | A930038C07Rik-Tg1-Cre | ACAd |
| 171275617 | A930038C07Rik-Tg1-Cre | ACAd |
| 125833030 | Rbp4-Cre_KL100 | ACAd |
| 496576666 | Rbp4-Cre_KL100 | ACAd |
| 526783054 | Rbp4-Cre_KL100 | ACAd |
| 293432575 | Tlx3-Cre_PL56 | ACAd |
| 293431869 | Tlx3-Cre_PL56 | ACAd |
| 539521918 | Cux2-IRES-Cre | ACAd |
| 522774029 | Cux2-IRES-Cre | ACAd |
| 583747816 | Emx1-IRES-Cre | ACAd |
| 478491090 | Emx1-IRES-Cre | ACAd |
| 478376911 | Emx1-IRES-Cre | ACAd |
| 584194481 | Emx1-IRES-Cre | ACAd |
| 501483880 | Emx1-IRES-Cre | ACAd |
| 183472596 | Grp-Cre_KH288 | ACAd |
| 112514202 | Wild-type | ACAv |
| 139520203 | Wild-type | ACAv |
| 159319654 | A930038C07Rik-Tg1-Cre | ACAv |
| 298797288 | A930038C07Rik-Tg1-Cre | ACAv |
| 286482701 | Rbp4-Cre_KL100 | ACAv |
| 161458737 | Rbp4-Cre_KL100 | ACAv |
| 126117554 | Rbp4-Cre_KL100 | ACAv |
| 294434161 | Cux2-IRES-Cre | ACAv |
| 571823196 | Cux2-IRES-Cre | ACAv |
| 183470468 | Cux2-IRES-Cre | ACAv |
| 183461297 | Cux2-IRES-Cre | ACAv |
| 265288825 | Grp-Cre_KH288 | ACAv |
| 265291552 | Grp-Cre_KH288 | ACAv |
| 184213701 | Grp-Cre_KH288 | ACAv |
| 266583498 | Grp-Cre_KH288 | ACAv |
| 112596790 | Wild-type | AId |
| 272737914 | Wild-type | AId |
| 267397941 | A930038C07Rik-Tg1-Cre | AId |
| 296048512 | Rbp4-Cre_KL100 | AId |
| 294525229 | Tlx3-Cre_PL56 | AId |
| 299783689 | Cux2-IRES-Cre | AId |
| 296047806 | Cux2-IRES-Cre | AId |
| 264076854 | Cux2-IRES-Cre | AId |
| 184168899 | Grp-Cre_KH288 | AId |
| 286775476 | A930038C07Rik-Tg1-Cre | Alp |
| 183330908 | Rbp4-Cre_KL100 | Alp |
| 272827141 | Cux2-IRES-Cre | Alp |
| 166153483 | Rbp4-Cre_KL100 | Alv |
| 113144533 | Wild-type | BLA |
| 120282354 | Wild-type | BLA |
| 277710753 | Wild-type | BLA |
| 113935990 | Wild-type | BMA |
| 115958115 | Wild-type | BMA |
| 125832322 | Wild-type | BMA |
| 146012934 | Wild-type | BMA |
| 273028004 | Wild-type | BMA |
| 127761449 | Wild-type | CEA |
| 146795148 | Wild-type | CEA |
| 112459547 | Wild-type | CEA |
| 180404418 | Wild-type | GU |
| 292375491 | A930038C07Rik-Tg1-Cre | GU |
| 249402048 | Rbp4-Cre KL100 | GU |
| 516846757 | Cux2-IRES-Cre | LA |
| 100141780 | Wild-type | MOp |
| 180720175 | Wild-type | MOp |
| 127084296 | Wild-type | MOp |
| 584903636 | Wild-type | MOp |
| 126909424 | Wild-type | MOp |
| 100141273 | Wild-type | MOp |
| 100141563 | Wild-type | MOp |
| 272697944 | Wild-type | MOp |
| 166461193 | A930038C07Rik-Tg1-Cre | MOp |
| 287461719 | A930038C07Rik-Tg1-Cre | MOp |
| 166082842 | Rbp4-Cre KL100 | MOp |
| 591612976 | Tlx3-Cre PL56 | MOp |
| 182616478 | Cux2-IRES-Cre | MOp |
| 180709230 | Wild-type | ORBI |
| 112306316 | Wild-type | ORBI |
| 170721670 | Wild-type | ORBI |
| 180673746 | Wild-type | ORBI |
| 168164972 | A930038C07Rik-Tg1-Cre | ORBI |
| 156741826 | Rbp4-Cre KL100 | ORBI |
| 183471174 | Cux2-IRES-Cre | ORBI |
| 288324211 | Grp-Cre KH288 | ORBI |
| 112423392 | Wild-type | ORBvl |
| 158435116 | Wild-type | ORBvl |
| 167902586 | Rbp4-Cre KL100 | ORBvl |
| 287769286 | Rbp4-Cre KL100 | ORBvl |
| 480994108 | Rbp4-Cre KL100 | ORBvl |
| 293433283 | Tlx3-Cre PL56 | ORBvl |
| 522773270 | Cux2-IRES-Cre | ORBvl |
| 266564027 | Grp-Cre KH288 | ORBvl |
| 157711748 | Wild-type | PL |
| 292374777 | A930038C07Rik-Tg1-Cre | PL |
| 263106036 | Rbp4-Cre KL100 | PL |
| 545434921 | Tlx3-Cre PL56 | PL |
| 278433737 | Cux2-IRES-Cre | PL |
| 287494320 | Cux2-IRES-Cre | PL |
| 294396492 | Cux2-IRES-Cre | PL |
| 478376197 | Emx1-IRES-Cre | PL |
| 266582782 | Grp-Cre KH288 | PL |
| 112951804 | Wild-type | SSp-bfd |
| 126907302 | Wild-type | SSp-bfd |
| 100142655 | Wild-type | SSp-bfd |
| 127866392 | Wild-type | SSp-bfd |
| 112882565 | Wild-type | SSp-bfd |
| 100141473 | Wild-type | SSp-bfd |
| 182467026 | A930038C07Rik-Tg1-Cre | SSp-bfd |
| 177907797 | A930038C07Rik-Tg1-Cre | SSp-bfd |
| 643749624 | A930038C07Rik-Tg1-Cre | SSp-bfd |
| 644250774 | A930038C07Rik-Tg1-Cre | SSp-bfd |
| 647806688 | Rbp4-Cre_KL100 | SSp-bfd |
| 648253235 | Rbp4-Cre_KL100 | SSp-bfd |
| 511550172 | Rbp4-Cre_KL100 | SSp-bfd |
| 272735030 | Rbp4-Cre_KL100 | SSp-bfd |
| 183171679 | Rbp4-Cre_KL100 | SSp-bfd |
| 297654263 | Tlx3-Cre_PL56 | SSp-bfd |
| 288170549 | Tlx3-Cre_PL56 | SSp-bfd |
| 670228985 | Tlx3-Cre_PL56 | SSp-bfd |
| 560736273 | Cux2-IRES-Cre | SSp-bfd |
| 168003640 | Cux2-IRES-Cre | SSp-bfd |
| 266486371 | Cux2-IRES-Cre | SSp-bfd |
| 294435580 | Cux2-IRES-Cre | SSp-bfd |
| 583748537 | Emx1-IRES-Cre | SSp-bfd |
| 657041814 | Emx1-IRES-Cre | SSp-bfd |
| 112791318 | Wild-type | SSp-II |
| 112935169 | Wild-type | SSp-II |
| 114292355 | Wild-type | SSp-II |
| 166323186 | A930038C07Rik-Tg1-Cre | SSp-II |
| 264629246 | A930038C07Rik-Tg1-Cre | SSp-II |
| 272698650 | Rbp4-Cre_KL100 | SSp-II |
| 297652799 | Rbp4-Cre_KL100 | SSp-II |
| 182803137 | Rbp4-Cre_KL100 | SSp-II |
| 591622344 | Tlx3-Cre_PL56 | SSp-II |
| 296052839 | Tlx3-Cre_PL56 | SSp-II |
| 584902900 | Tlx3-Cre_PL56 | SSp-II |
| 298830161 | Cux2-IRES-Cre | SSp-II |
| 266644610 | Cux2-IRES-Cre | SSp-II |
| 178284661 | Cux2-IRES-Cre | SSp-II |
| 180717881 | Wild-type | SSp-m |
| 157654817 | Wild-type | SSp-m |
| 112936582 | Wild-type | SSp-m |
| 114290938 | Wild-type | SSp-m |
| 100149969 | Wild-type | SSp-m |
| 298796577 | A930038C07Rik-Tg1-Cre | SSp-m |
| 272928602 | Rbp4-Cre_KL100 | SSp-m |
| 303785454 | Rbp4-Cre_KL100 | SSp-m |
| 303614706 | Tlx3-Cre_PL56 | SSp-m |
| 294397199 | Cux2-IRES-Cre | SSp-m |
| 278317945 | Cux2-IRES-Cre | SSp-m |
| 126908007 | Wild-type | SSp-n |
| 112162251 | Wild-type | SSp-n |
| 112373124 | Wild-type | SSp-n |
| 298326521 | A930038C07Rik-Tg1-Cre | SSp-n |
| 171276330 | A930038C07Rik-Tg1-Cre | SSp-n |
| 120875816 | Rbp4-Cre_KL100 | SSp-n |
| 249327301 | Rbp4-Cre_KL100 | SSp-n |
| 126852363 | Wild-type | SSp-tr |
| 100141495 | Wild-type | SSp-tr |
| 281459203 | A930038C07Rik-Tg1-Cre | SSp-tr |
| 298182842 | Tlx3-Cre PL56 | SSp-tr |
| 180718587 | Wild-type | SSp-ul |
| 112229814 | Wild-type | SSp-ul |
| 148964212 | Wild-type | SSp-ul |
| 292792016 | A930038C07Rik-Tg1-Cre | SSp-ul |
| 266249483 | Rbp4-Cre KL100 | SSp-ul |
| 286312782 | Rbp4-Cre_KL100 | SSp-ul |
| 303615412 | Tlx3-Cre PL56 | SSp-ul |
| 298106713 | Cux2-IRES-Cre | SSp-ul |
| 286300594 | Cux2-IRES-Cre | SSp-ul |
| 180601025 | Wild-type | SSp-un |
| 112514915 | Wild-type | SSs |
| 117298988 | Wild-type | SSs |
| 174360333 | Wild-type | SSs |
| 156395997 | A930038C07Rik-Tg1-Cre | SSs |
| 286834976 | A930038C07Rik-Tg1-Cre | SSs |
| 168163498 | Rbp4-Cre_KL100 | SSs |
| 298179622 | Rbp4-Cre_KL100 | SSs |
| 120916102 | Rbp4-Cre_KL100 | SSs |
| 286314623 | Rbp4-Cre_KL100 | SSs |
| 303616833 | Tlx3-Cre PL56 | SSs |
| 288171256 | Tlx3-Cre PL56 | SSs |
| 297855879 | Cux2-IRES-Cre | SSs |
| 287951098 | Cux2-IRES-Cre | SSs |
| 577773267 | Cux2-IRES-Cre | SSs |

**Extended Data Table 7.**
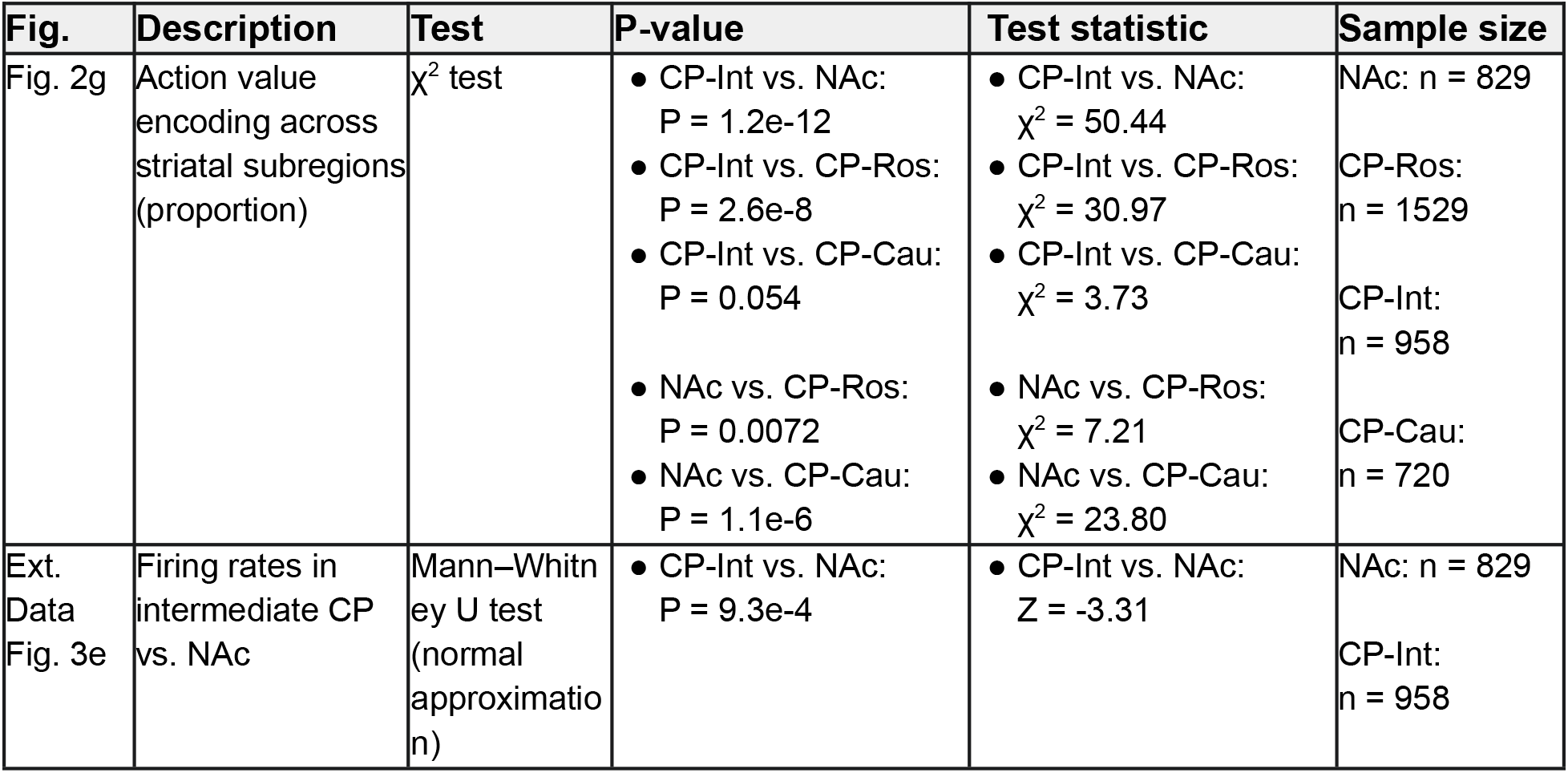
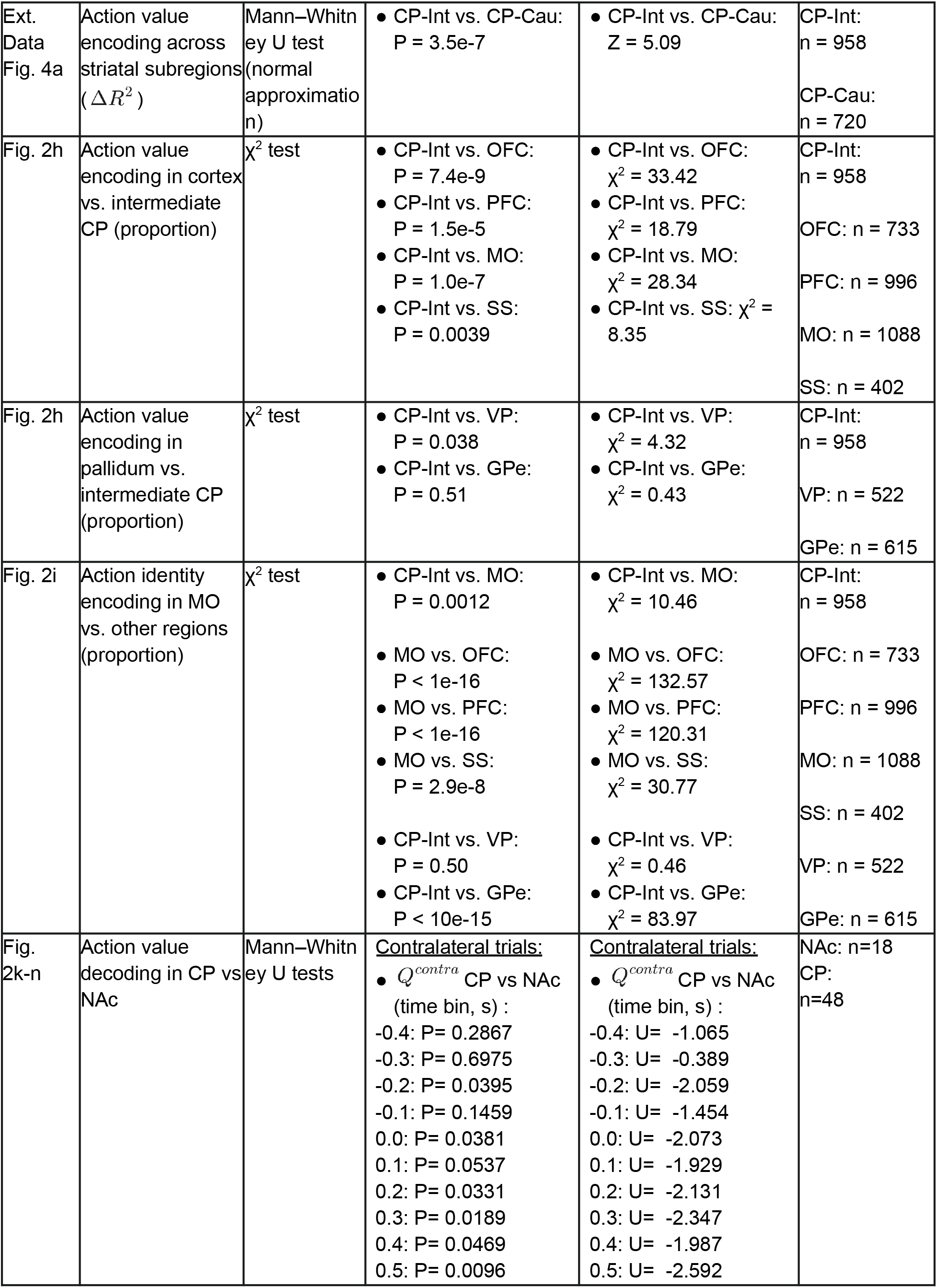
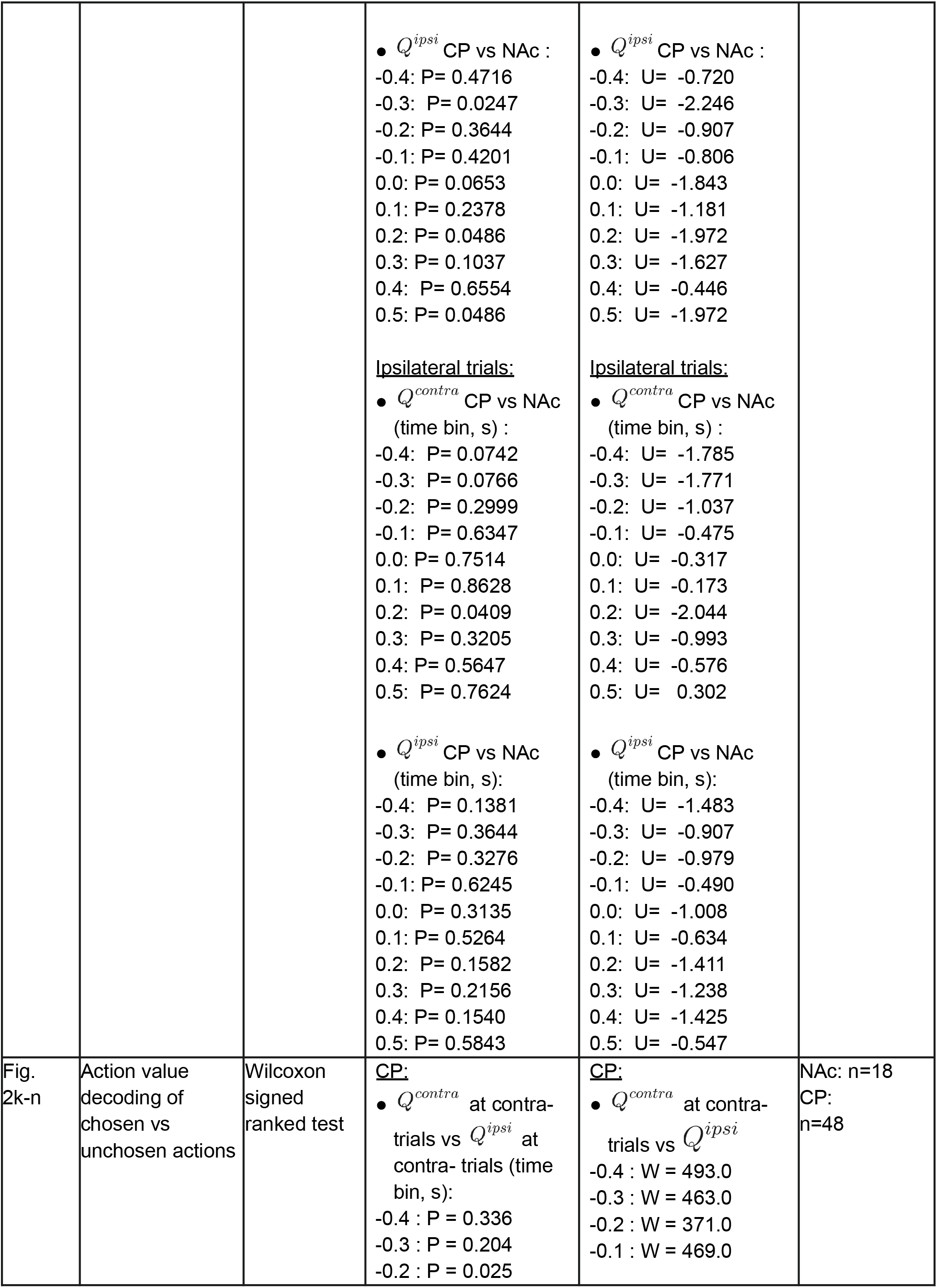
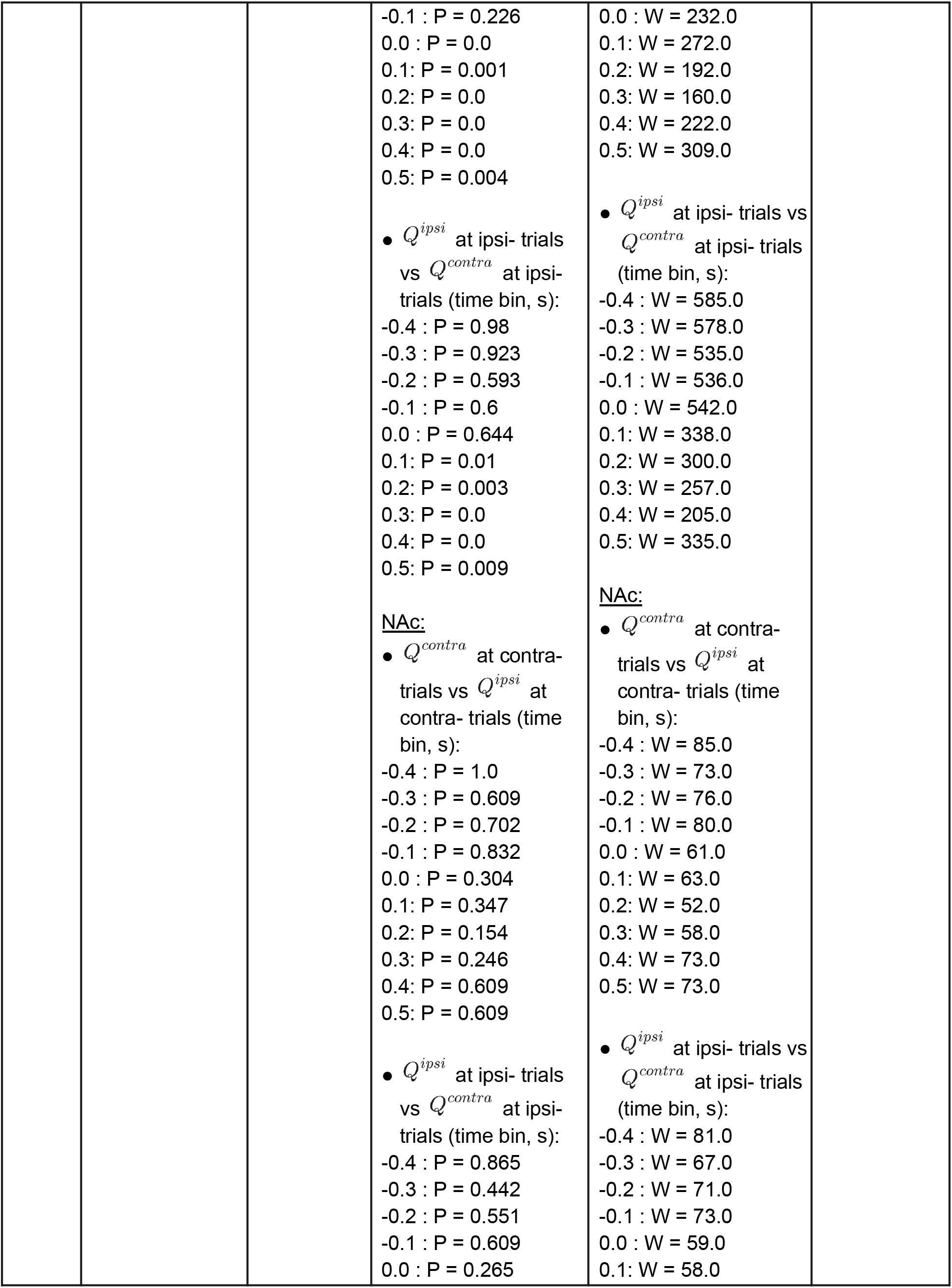
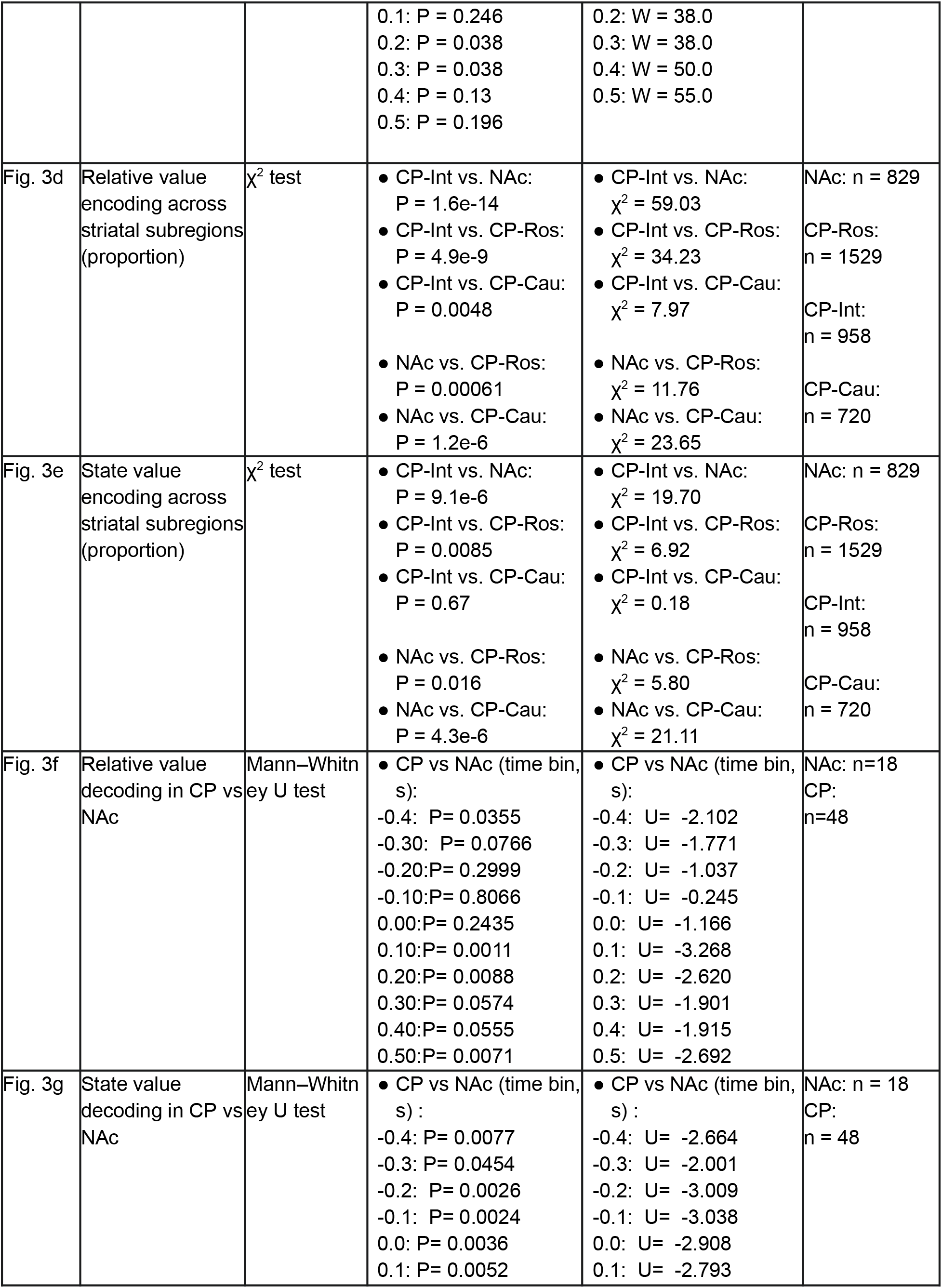
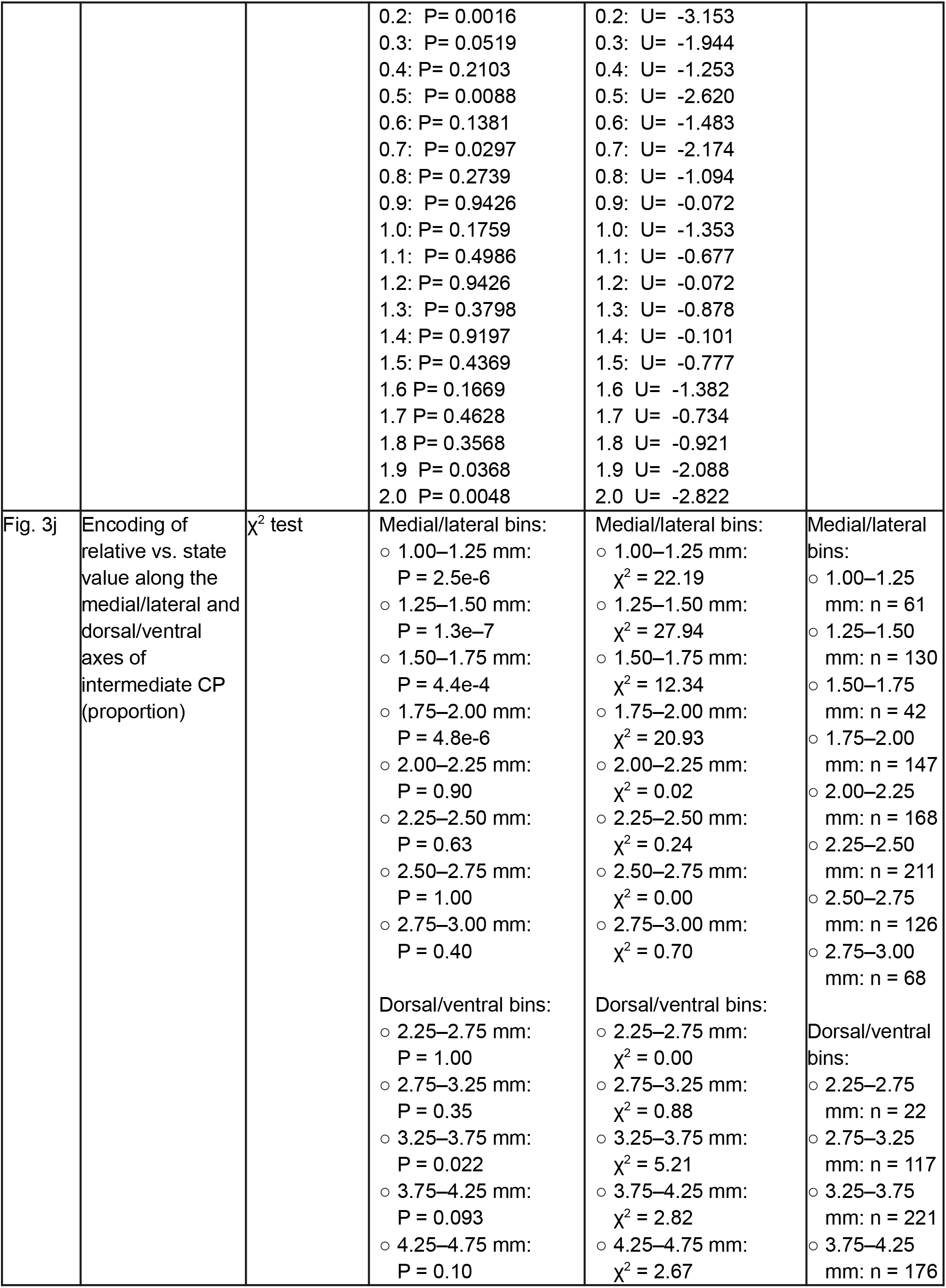
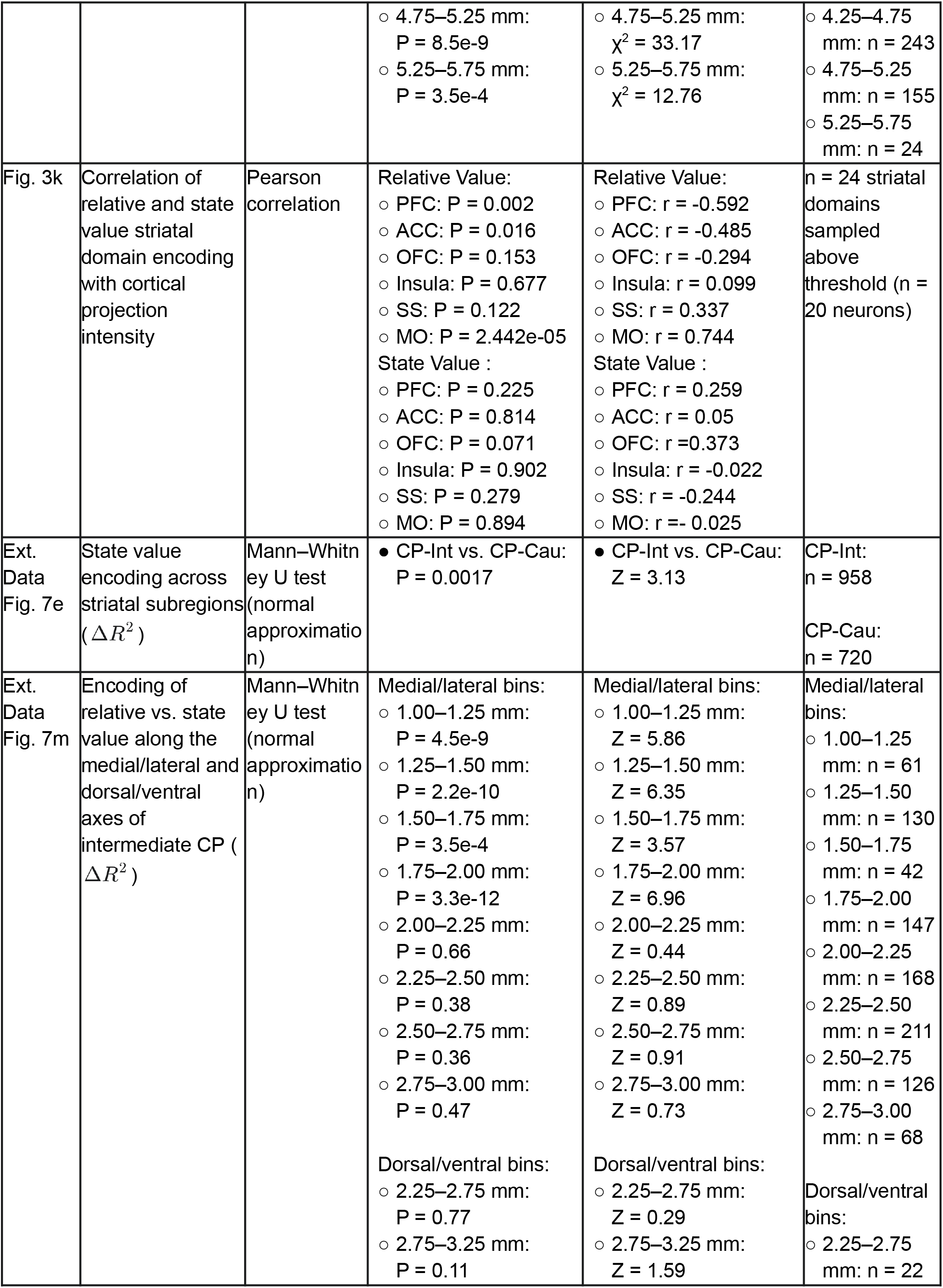
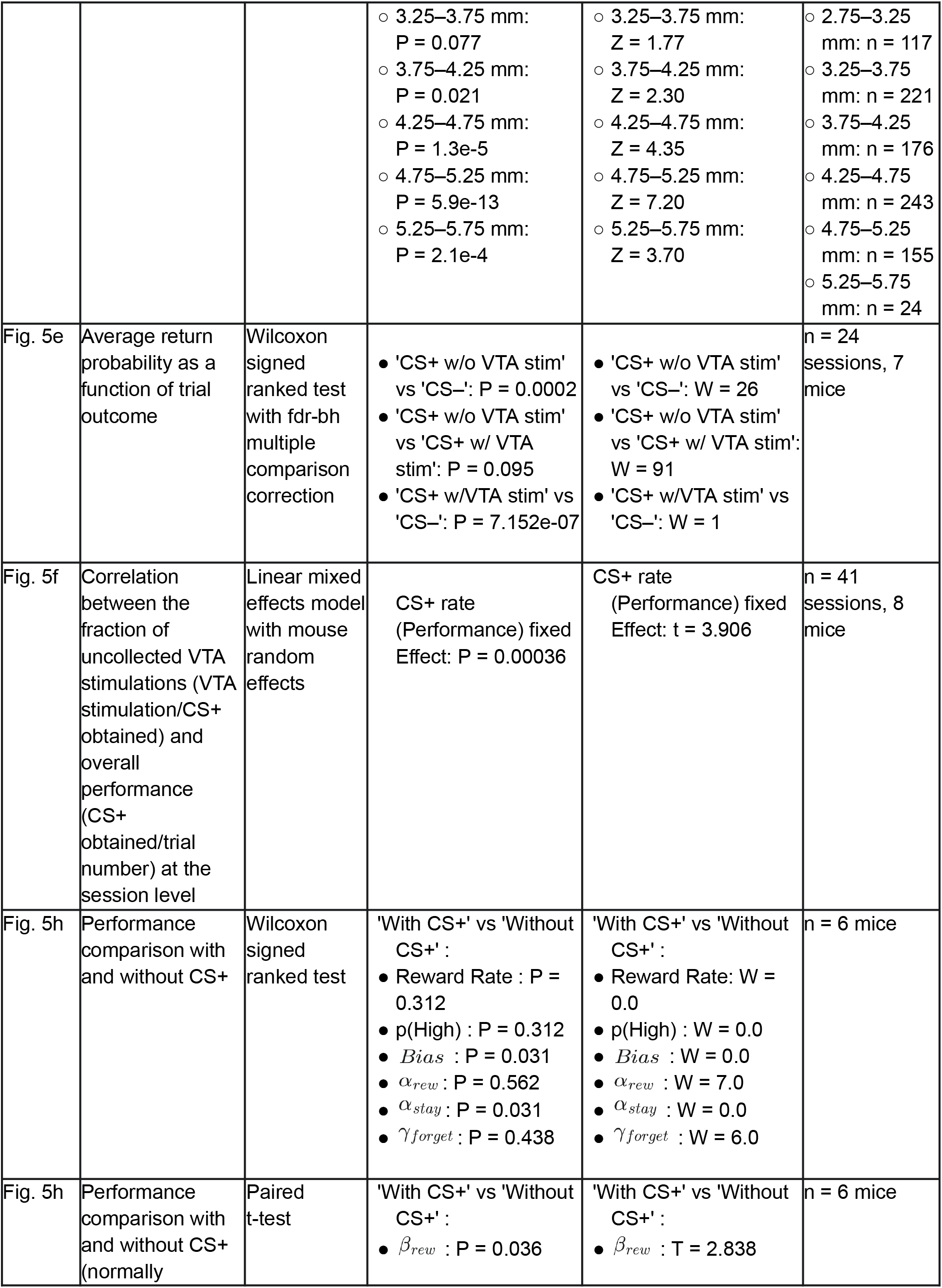
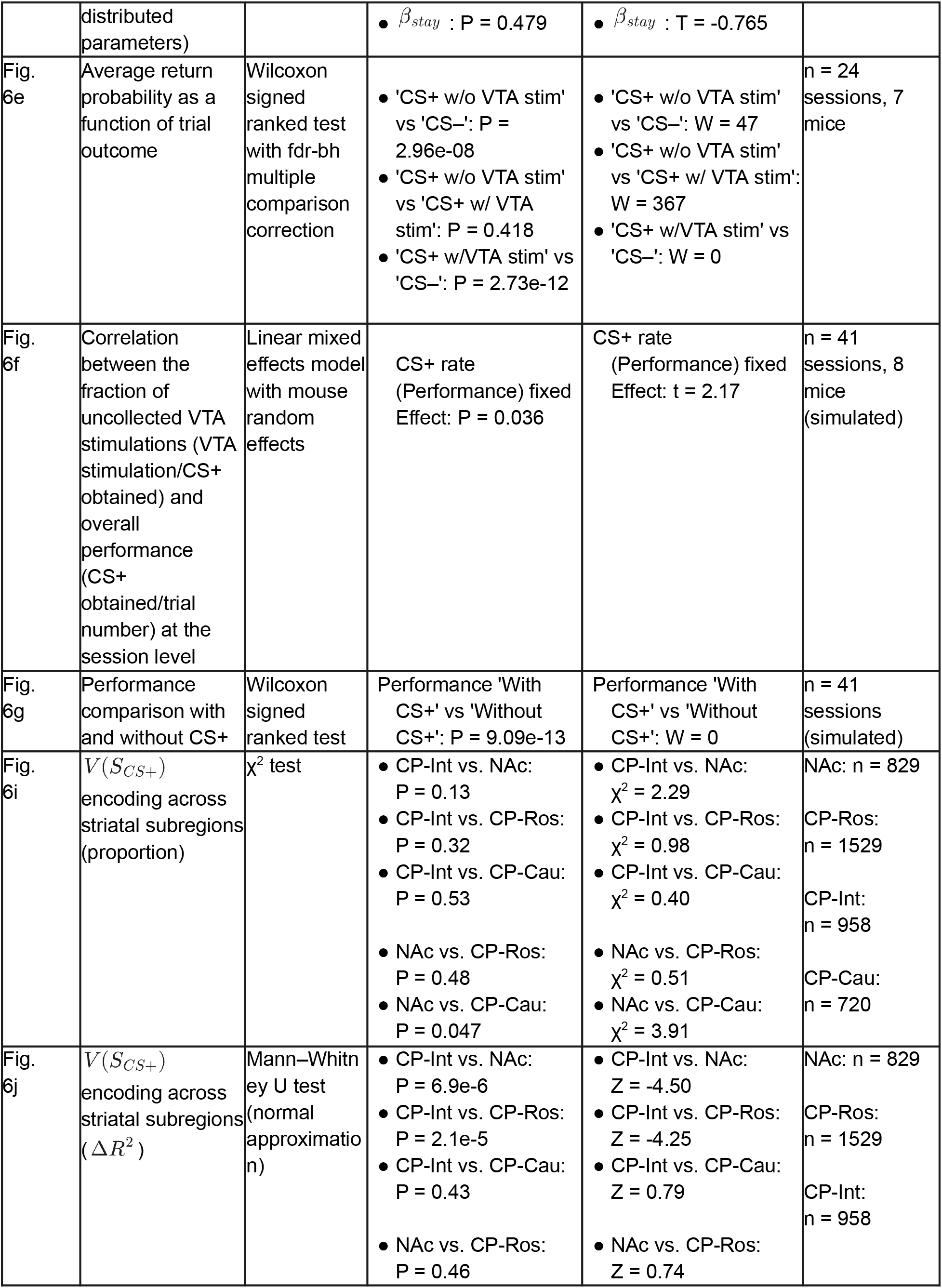
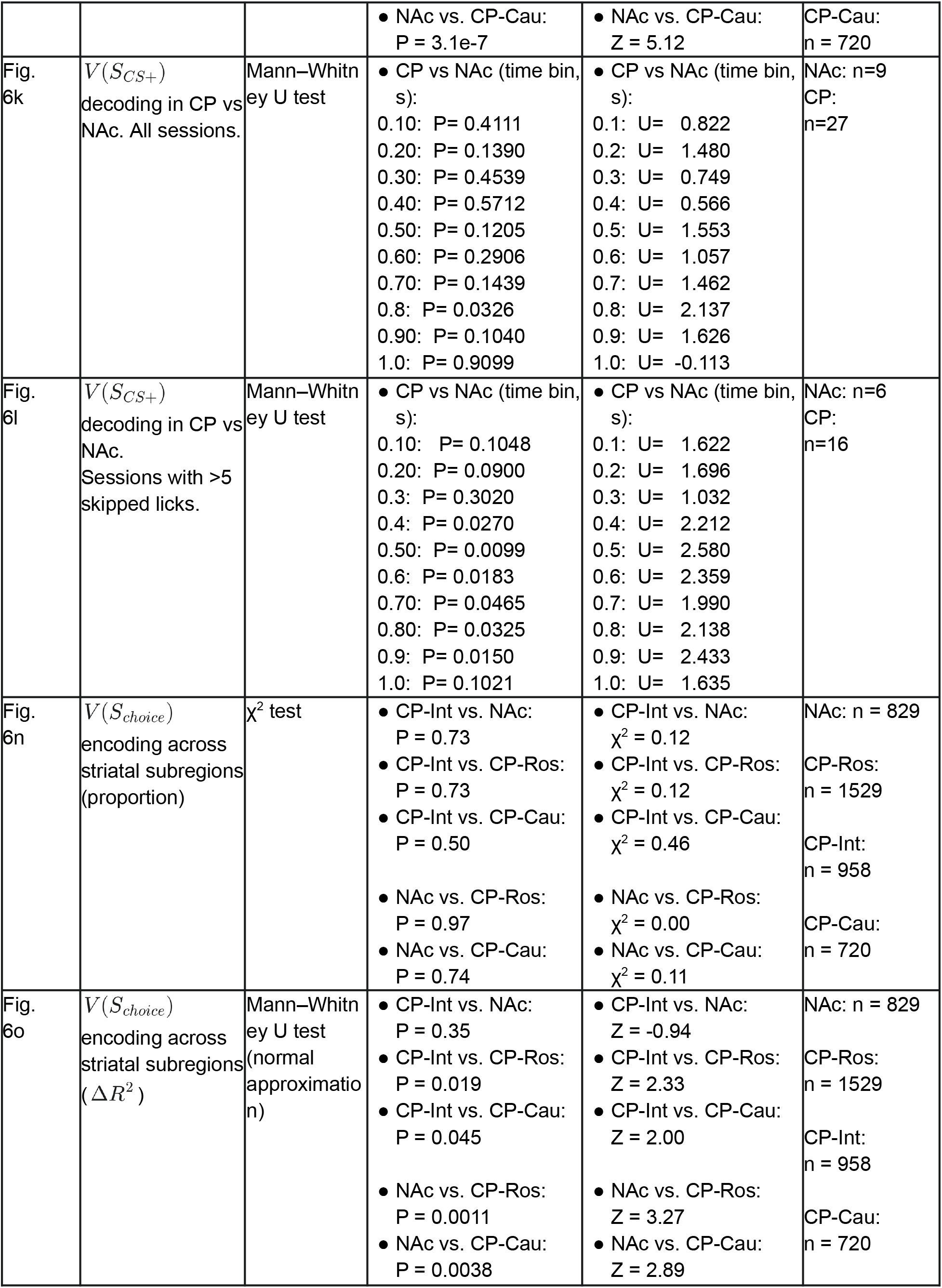
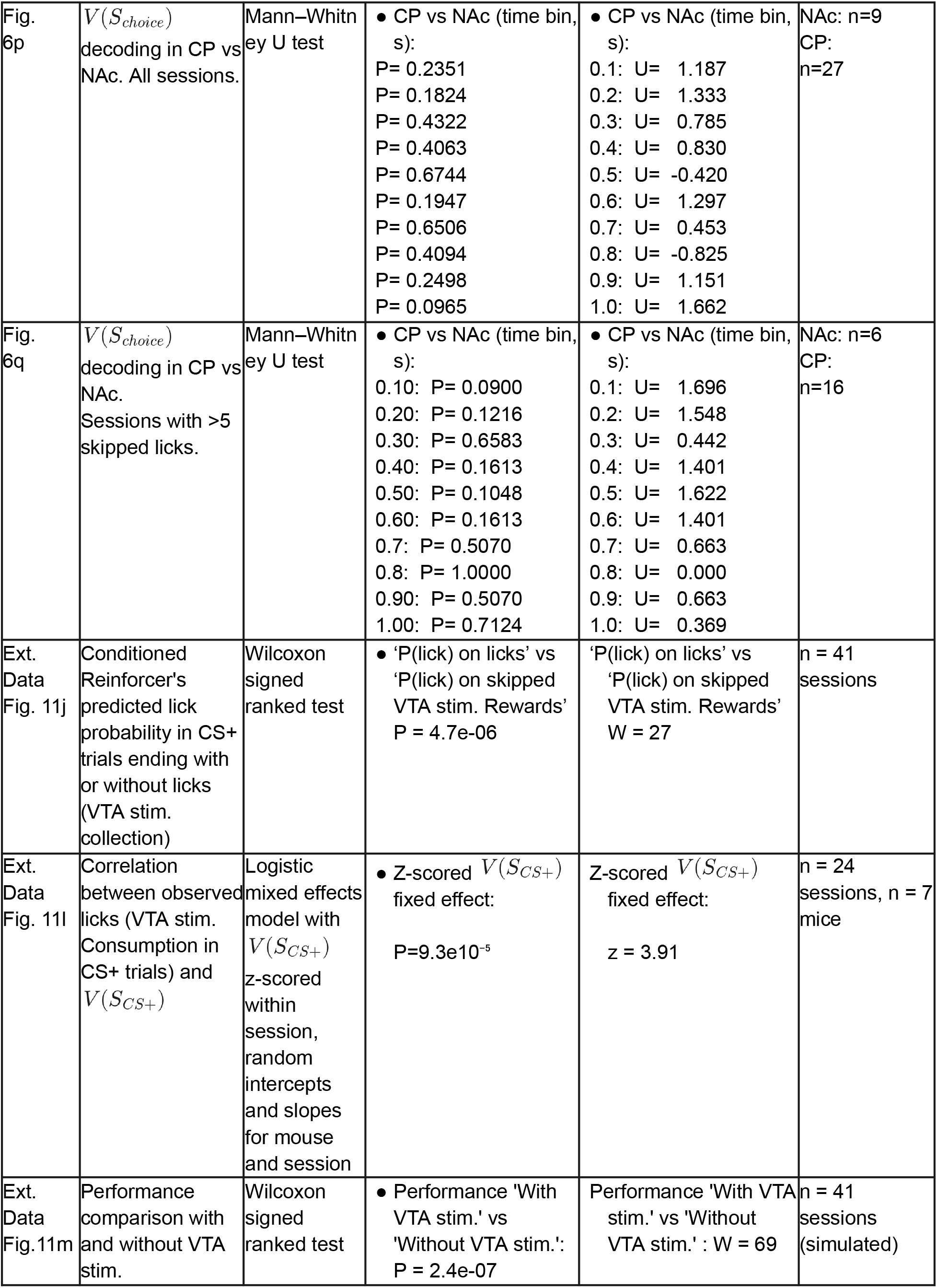
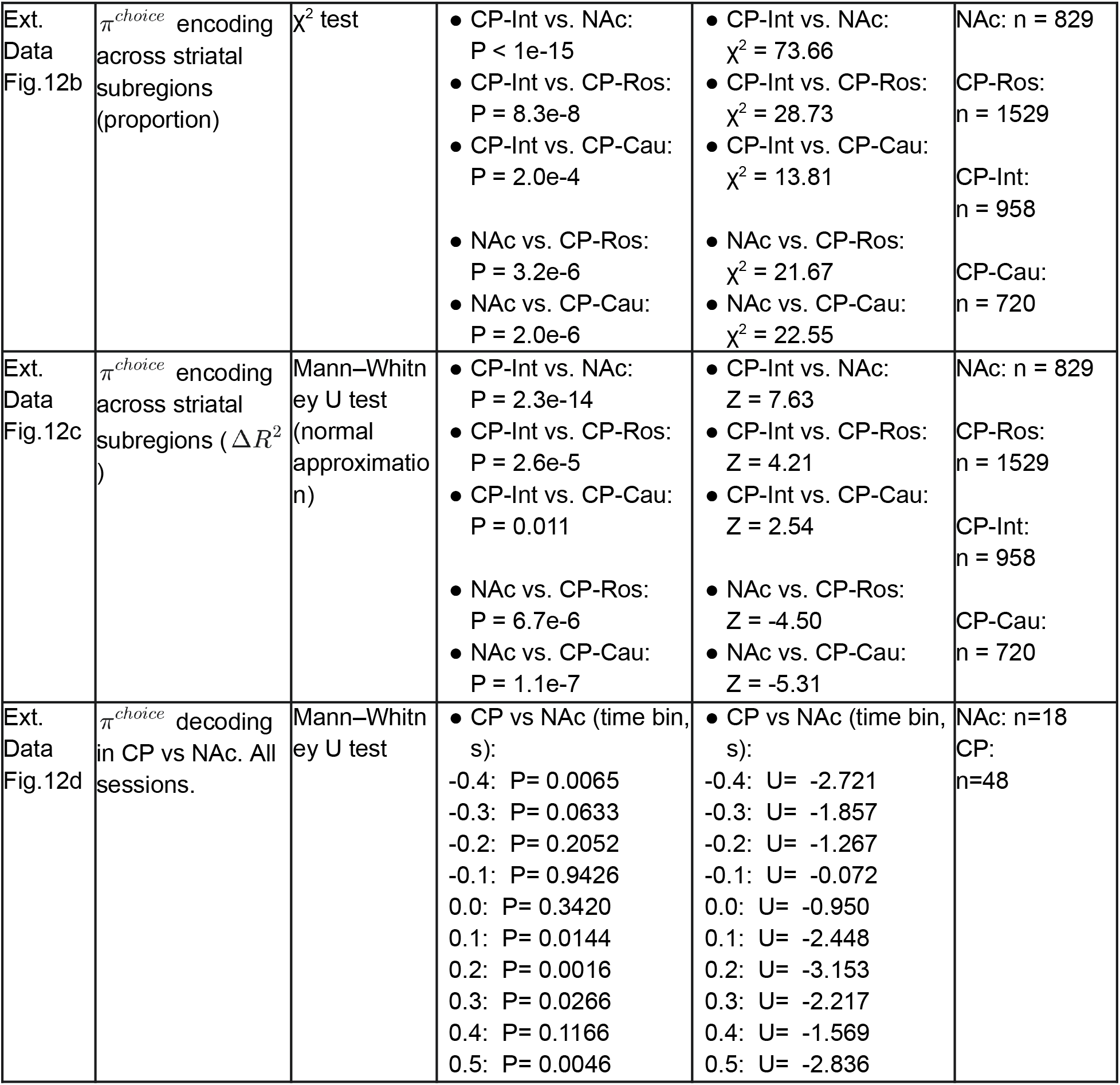
Summary of statistical comparisons.

